# Native and regenerated cellulose show similar environmental biodegradation behavior across global terrestrial and aquatic ecosystems

**DOI:** 10.64898/2025.12.24.696393

**Authors:** Christian Lott, Alisa Berning, Victoria Eckerle, Ann-Sophie Rupp, Julian Rau, Giuseppe Suaria, Mireno Borghini, Stefano Miserocchi, Francesco Paladini de Mendoza, Federico Giglio, Markus Lasut, Andreas Eich, Miriam Weber

## Abstract

Cellulose is the most abundant natural polymer and serves as the structural scaffolding molecule of plants, and materials from cellulose fibers are considered important to the global shift toward renewable materials. Yet, uncertainty remains about their persistence in natural environments. Here we show that native and regenerated cellulose, ranging from cotton and linen to viscose, modal, and lyocell, despite minor structural differences, are biodegraded at comparable rates, indicating that there is no scientifically justified distinction regarding environmental behavior. We assessed the biodegradation of diverse cellulosic materials, including powders, loose fibers, fabrics, and nonwovens, under technical and natural conditions across soil, home compost, freshwater, and marine coastal and deep-sea environments. Our study combined a total of 152 scenarios with laboratory tests, mesocosm and field experiments, spanning from polar to tropical regions and temperatures from -1.8 °C in the high Arctic Ocean to 38.4 °C in a marine beach in the Mediterranean Sea, and between -6.1 to 54.1 °C in agricultural soil. All neat cellulosic fibers showed inherent biodegradability, with biodegradation half-lives typically ranging from weeks to months. Biodegradation rates were primarily driven by water availability, temperature, and nutrient levels, while the role of oxygen was indifferent. Standardized lab tests aligned well with field observations, confirming their validity for assessing inherent biodegradability with environmental relevance. However, biodegradation of biodegradable polymers in real-world scenarios is influenced by product-level modifications such as dyeing and finishing. By distinguishing the effects of fiber chemistry separately from those of finishing treatments, this study clarifies the role of cellulose in a sustainable materials economy and supports evidence-based regulation of fiber biodegradability claims and product labeling.

## 1. INTRODUCTION

Cellulose is the most abundant natural polymer and serves as the structural scaffolding molecule of plants. Plant fibers have been used since early human history, and still today flax/linen, hemp, abacá, sisal, ramie, coir, and cotton (Müssig, 2010) continue to be cultivated and used for technical applications and clothing. Cellulose-based materials such a paper are increasingly used to replace, for example, plastic packaging. Natural plant fibers contain cellulose, hemicelluloses and lignin (lignocellulose) and are traditionally and industrially treated to optimize the properties of the final product. Cellulose is a polysaccharide composed of glucose monomers that are linked by ß-1,4-glucosidic bonds (Huang and Fu, 2013), which assemble into crystalline and amorphous domains stabilized by an extensive hydrogen-bonding network. The cellulose molecule can be dissolved using suitable chemicals and made into a viscous paste. This regeneration of cellulose is a technical process that enables its use as a base material for fiber wet spinning to achieve standardized material properties. The final regenerated cellulose is chemically identical to the original cellulose, with slight differences in crystallinity, molecular orientation, and the adsorption properties of water and dye stuffs (Mahltig, 2018). Native plant fibers consist mainly of the native cellulose I, which can be converted into cellulose II, for example by mercerization, the treatment with concentrated aqueous sodium hydroxide (NaOH) to improve functional properties. Regenerated cellulosic fibers (RCFs) such as viscose, modal, or lyocell are composed of cellulose II, which results from the dissolution and subsequent regeneration of native cellulose. Today, viscose, and modal with a slightly modified production process, and lyocell with a different dissolution process, are the main generic types of regenerated cellulose. For an in-depth review, see Huber et al., 2026.

This study only considered the grades of the industrial raw fiber as they leave the production plant for further processing. For these natural and regenerated cellulosic fibers as base material we use the term ‘*neat’* throughout the manuscript to differentiate them from fibers or products that may have undergone further treatment steps along the value chain until the final product. The focus of this study was on the biodegradability and environmental biodegradation rates of natural cellulose and RCF, as part of the so-called man-made cellulosic fibers (MMCF), excluding, for example, cellulose acetate, which is chemically modified.

Biodegradation of a polymer is defined as the complete microbial conversion of all its organic constituents to carbon dioxide, new microbial biomass and mineral salts under oxic conditions or to carbon dioxide, methane, new microbial biomass and mineral salts under anoxic conditions (SAPEA, 2020). The natural decay of plant debris, such as leaves, straw, and wood, in the environment can provide insight into the biodegradation of cellulosic materials in nature. Biodegradation of cellulose in the aquatic environment has been studied historically as far back as the 1970s in the context of paper mill effluents (Hofsten and Edberg, 1972). Biopolymers, i.e. polymers produced by living organisms, such as cellulose are biodegradable in natural environments. The rate of degradation depends on both the polymer type and the environmental conditions (Erdal and Hakkarainen 2022). Because polymers are too large for direct uptake into microbial cells, they must be broken down first by abiotic processes or extracellular microbial enzymes into smaller compounds, typically below a molecular weight of 500 to 1000 g mol^-1^. These dissolved compounds can then be assimilated by microorganisms and further metabolized intracellularly.

In nature, cellulose biodegradation is carried out by microorganisms through cellulases secreted by bacteria and fungi. Cellulose can be directly degraded by cellulosomes, large extracellular enzyme complexes (Warren, 1996). These include endoglucanases, which hydrolyze β-1,4-glycosidic linkages in amorphous cellulose, and cellobiohydrolases, which act on cellulose chain ends (Wilson 2008). Lytic polysaccharide monooxygenases also contribute by oxidatively cleaving glycosidic bonds. Upon assimilation and metabolization by microbes, biodegradation of cellulose ultimately leads to full mineralization. Some cellulases exhibit processivity, which allows for the continuous degradation of crystalline cellulose without detaching from the substrate. Fungal and bacterial cellobiohydrolases independently evolved this ability through similar structural strategies, providing an example of convergent protein evolution (Uchiyama et al. 2020).

Classically, there is an (assumed) hierarchy of biodegradation activity across different environmental compartments with respect to biodegradable polymers where these materials biodegrade faster in industrial compost than in soil, faster in soil than in freshwater, and faster in freshwater than in marine water (De Wilde 2020; Erdal and Hakkarainen 2022). This concept was based on a wealth of observations from standard laboratory biodegradability tests and has been adopted for the biodegradation potential of natural settings:

Soil >>>> freshwater >>> estuarine >> marine coastal > deep-sea

However, results from field and mesocosm studies by us and others suggest that under conditions in the open environment the interplay of abiotic and biotic factors is more complex to justify such hierarchical simplification. Factors influencing the biodegradation of organic matter in general include water availability, temperature, the concentration of oxygen and of nutrients - mainly nitrogen and phosphorous compounds essential for the formation of new microbial biomass - and the abundance of capable microbes that possess the appropriate enzymes.

The biodegradation of natural polymers is known to have limits, which, at extremes, can lead to the preservation of plant fibers, animal hair, collagen or melanin over thousands or even millions of years as fossils. Natural preservation can be caused by dryness or anoxia.

To give a comprehensive overview of the biodegradation behavior of native and regenerated cellulosic fibers in technical and natural scenarios, we present evidence from experimental data from standard biodegradability tests in laboratory respirometers in the form of mineralization endpoints, and from additional disintegration tests. We also conducted mesocosm tests, where natural scenarios were simulated in tanks with natural soil, freshwater, or seawater and aquatic sediments of different origins and properties. The disintegration of sheets of cellulose filter paper, protected from mechanical stress, and their gradual loss over time was measured as a proxy for biodegradation. To cover the environmental fate of cellulose as broadly as possible, we included qualitative observational evidence from several *in-situ* studies on the disintegration behavior of textiles and natural materials under marine conditions, in shallow coastal habitats and for the first time from the deep sea.

The main questions of this study were: Is the biodegradability of regenerated cellulose comparable to that of native cellulose? Are there differences in biodegradability between the different types and forms? Does environmental biodegradability differ in technical and natural environments? What are the typical biodegradation rates or lifetimes? Which factors determine the rate and extent of the biodegradation of cellulosic fibers in the environment?

Addressing these questions has implications far beyond cellulosic fibers: it informs how we define and assess “biodegradability” of natural and synthetic biodegradable materials in sustainability frameworks, and how material design, processing, and environmental policy should converge to ensure truly circular bio-based economies.

## 2. MATERIALS AND METHODS

### 2.1 Test Scheme

Biodegradation was assessed using three complementary experimental approaches in laboratory, mesocosm (tank) and field tests (fig. 1) according to standard and adapted methods (tab.1). The first set of experiments comprised standardized laboratory biodegradability tests under controlled conditions to measure mineralization in respirometers. Another set of experiments evaluated environmental biodegradation, ideally conducted *in situ* to capture rates specific to the prevailing environmental scenario. To bridge the methodological gap between laboratory and field experiments, a set of mesocosm (i.e., tank) tests were employed. These allowed the use of larger amounts of soil, sediment or water, so-called environmental matrices, while offering the controlled adjustment of parameters such as temperature, oxygen availability, pH, and the nutrient regime. Both field and tank tests represented open systems in which the direct quantification of gas exchange (CO_2_ evolution or O_2_ consumption) is not feasible. Instead, the progression of material disintegration was monitored over time and used as a proxy for biodegradation. This approach is only valid if a) the material has previously demonstrated its biodegradability in standardized laboratory tests, and b) suitable measures are taken to avoid physical abrasion or premature fragment loss (Lott et al. 2020). Mere mass or material loss alone cannot be considered proof of biodegradation.

**Figure 1:**
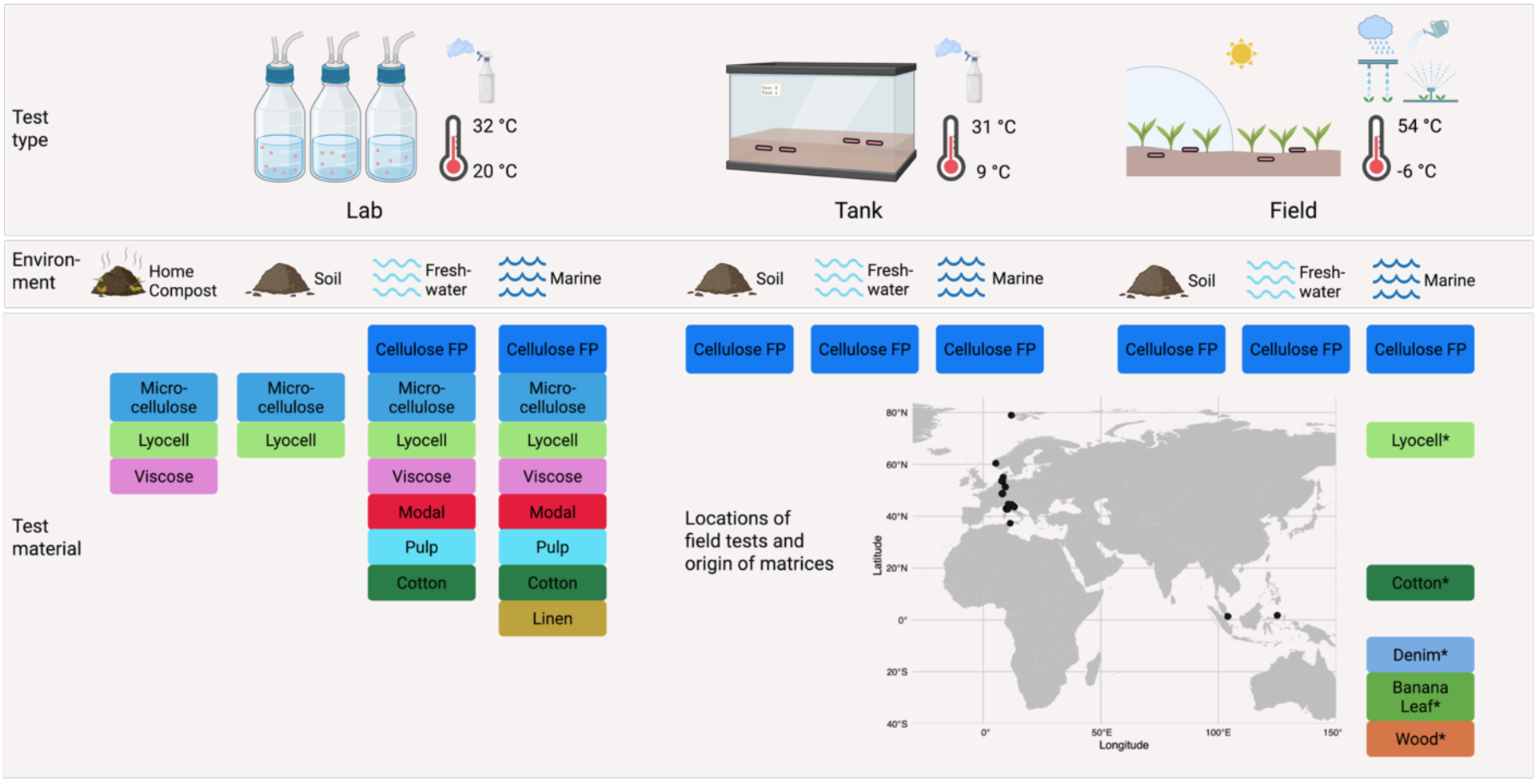
**Overview of all experiments** by test type (lab, tank, field), environmental scenario (home compost, soil, freshwater, marine), material types and geographical positions of field tests and provenance of environmental matrices as inocula (soil, freshwater and marine sediments) ranging from the polar High Arctic over cold- and warm-temperate Europe to tropical Southeast Asia. For details see table SI1. Asterisks (*) indicate scenarios of field tests which provided observational data only.

#### Experimental set 1: Standard biodegradability and disintegration tests in home compost, soil, water with activated sludge, freshwater sediment, and marine water and sediment

Most standardized laboratory tests (tab. 1) were contracted to Normec OWS (Ghent, Belgium). Field, tank and additional lab tests were conducted at HYDRA Marine Sciences. Generally, a standard biodegradability lab test of a carbon-based polymer is performed through respirometry via an incubation in a closed vessel with an inoculum (here called ‘matrix’), e.g. compost, water, or soil, containing natural microbes. In oxic conditions, carbon dioxide as the end-product of the metabolic conversion of the organic carbon (mineralization) from the test material is quantified over time. This provides information about the amount of carbon that has been completely converted at a given time point, i.e., the degree of mineralization, as well as the biodegradation rate. Alternatively, some standards also allow the use of oxygen consumption over time as a parameter to measure biodegradation. Standard laboratory tests are usually run under optimized conditions to address biodegrad*ability*, i.e., the *potential* of a material to be degraded at all by naturally occurring microorganisms (bacteria, archaea and fungi).

**Table 1:**
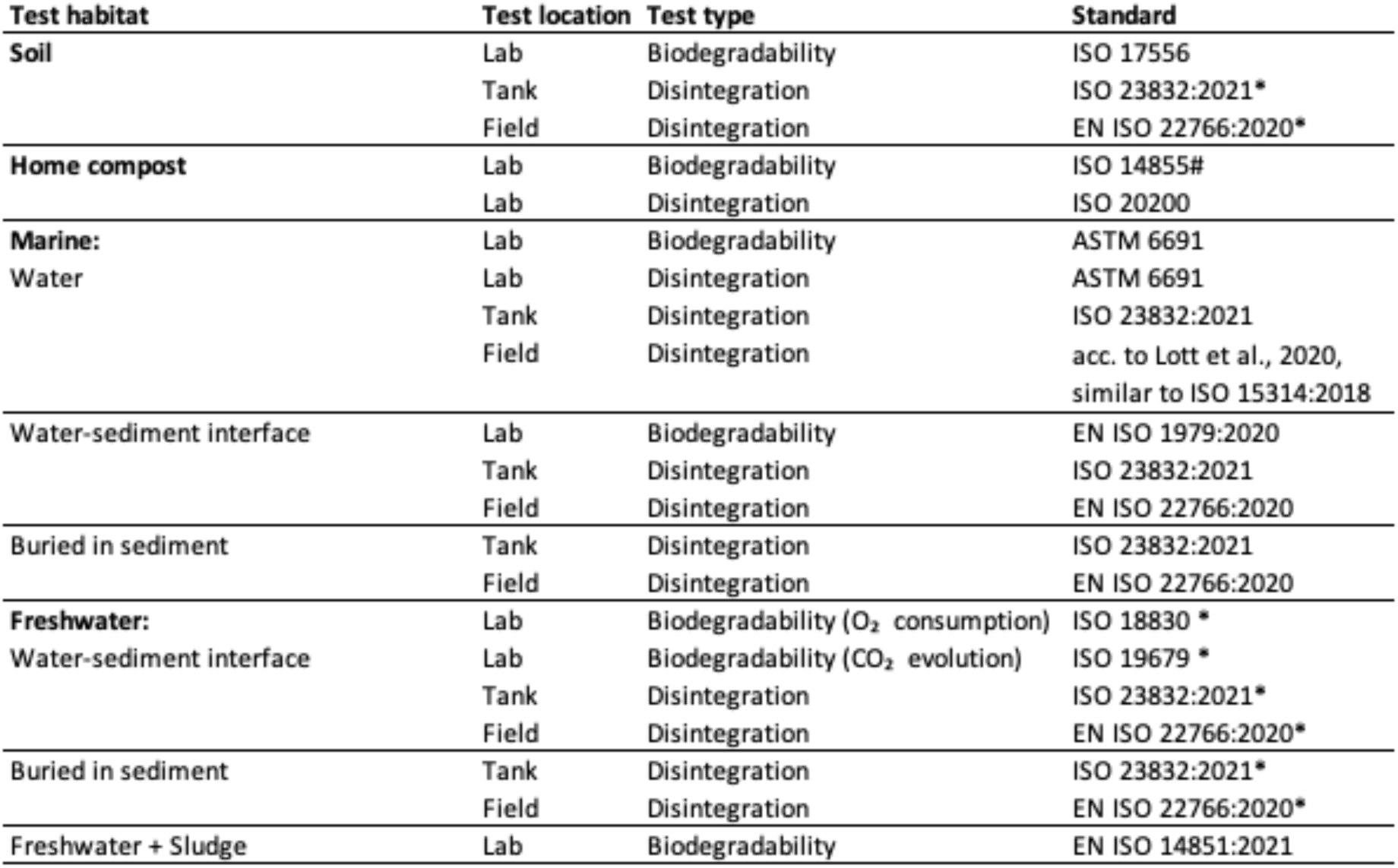
List of standard tests conducted under laboratory conditions, in open-system tanks and in the field. Test habitats, test locations, test types and standards are given. Test materials were prepared in the form of sheets, fibers or powder. The hash symbol (#) indicates that the temperature was adapted from the standard intended for industrial compostability to 28 ± 2 °C to simulate home composting conditions. The asterisk (*) indicates the cases for which the standards were adapted to freshwater or soil conditions due to the lack of appropriate standards. For open-system tank and field tests, the test materials were prepared according to Lott et al., 2020 or Eich et al., 2021.

The materials tested were microcrystalline cellulose powder (Avicell, Merck), ash-free cellulose filter paper (Whatman no. 4 or Macherey-Nagel MN 640 d), loose fibers, woven and nonwoven sheets of different forms of viscose, modal, lyocell, organic cotton, softwood paper pulp, and abacá pulp, all provided by Lenzing AG, Austria, and linen fiber (loose and woven) provided by the Alliance for European Flax-Linen & Hemp, Paris. For material details see tables SI1 – SI4 and for temperatures see tab. SI7.

#### Experimental set 2: Disintegration time series of cellulose filter paper in soil, freshwater and marine mesocosm tank and field tests

Field tests were conducted in the High Arctic ocean, in European agricultural fields and lakes, the Mediterranean Sea both coastal and in the deep sea, and in tropical Southeast Asia, mainly according to marine standards or adapted to freshwater and soil. Mesocosm tank tests were conducted using soil, sediment and water from some of these field sites. Temperature was recorded with automated temperature loggers (Tinytag Aquatic 2 –TG, Gemini Data Loggers, UK; HOBO Pendant, Onset Computer Corp., Bourne, MA, USA; or CTD sensors at the deep-sea moorings) near the samples (see tab. SI7).

In soil tests, cellulose filter paper was exposed for disintegration in tank and field experiments, laid on the soil either openly (surface) or covered with silage (fermented hay) or buried in the soil, in the open field or in a greenhouse tunnel without heating. The mean temperatures of the experiments ranged from 11 °C to 24 °C (min. -6.1 °C, max. 54.1 °C). The different soil types were agricultural soil, garden soil, forest soil and a mix of garden and forest soil from SW Germany. Different moisture levels from air-dry to wet were tested, with samples either not irrigated, exposed to natural precipitation, irrigated with water, or irrigated with water and compost tea prepared at the test site by the farmer. Compost tea is often used in organic farming and horticulture to enhance the microbial activity of the soil and consists of an exudate from compost, containing active microbes and nutrients.

Freshwater tests were conducted in a garden pond, two lakes and one creek in Germany, and in an irrigation pond in Italy or in tanks with matrices retrieved thereof, with a range of mean temperatures between 12 °C and 28 °C (min. 2.0 °C, max. 32.9 °C).

Marine tests covered a polar site in the High Arctic ocean at Spitsbergen, a public aquarium in Bergen, both Norway and several North Sea coastal sites in Germany, sites in the Mediterranean Sea (several beaches in the Tyrrhenian and Adriatic Sea, a lagoon, coastal seafloor sites, and deep-sea sites), and sites in tropical Southeast Asia (Singapore and Indonesia) covering beaches, mangroves, and coastal seafloor. The exact locations are described below (2.2 and tab. SI6). Marine mean temperatures ranged from 9 °C to 29 °C (min. -1.8 °C, max. 38.4 °C). For the temperatures in individual experiments see table SI7.

To distinguish experimental settings from each other, we termed a specific test scenario, i.e. a combination of material and test environment as ‘treatment’ and gave it an individual code T1, T2, T3, etc., nominated in the respective data graphs for identification. Different environmental scenarios were experimentally assessed for the biodegradation of cellulose filter paper which is regularly being used as a positive control and reference material in many biodegradation studies, and prescribed in standard methods. Ash-free cellulose filter paper (MN 640 d, 85 g/m², 170 µm thickness, Macherey-Nagel, Germany) was exposed to soil (tab. SI2), freshwater (tab. SI3) and marine (tab. SI4) conditions, in tank and field experiments.

Sheets of cellulose filter paper were cut to the size of 260 x 200 mm (field), 60 x 80 mm or 30 x 80 mm (field and tank), mounted in plastic frames (HDPE) and protected from physical damage by polyester mesh (SEFAR, Switzerland) as described before (Lott et al. 2020, Eich et al., 2021). After retrieval, the samples were carefully unpacked and cleaned from adhering soil or sediment in a shallow tray filled with water to avoid potential fragment loss, photographed in a standardized way and analyzed photogrammetrically (see below). Disintegration over time as a proxy for biodegradation was determined from time series of photographs.

The disintegration dynamics were statistically modelled to obtain a specific half-life (see below) for each combination of material and test scenario. The specific half-lives allowed to numerically compare the biodegradation behavior of a material under different conditions, or different materials under the same conditions and statistically rank them.

#### Experimental set 3: Semi-quantitative, observational tests of disintegration of textile samples and natural materials under marine shallow-water and deep-sea conditions

In addition to the systematic time series, textile samples of organic cotton fabric and lyocell fabric (provided by VAUDE, Tettnang, Germany), a used organic cotton t-shirt (C&A, bought from a warehouse), a used denim jeans (LEVIS, bought from a warehouse) along with the same cellulose filter paper described above (Macherey & Nagel) and natural cellulosic materials (e.g., fresh banana leaves cut directly from the tree; sheets of oak wood veneer, untreated, from a furniture carpenter), with no or only a few replicates per treatment each, were exposed in marine field experiments in tropical Southeast Asia (Bangka Island, Sulawesi, Indonesia), in coastal and deep-sea locations of the Mediterranean Sea, and in the High Arctic. Additionally, a set of face masks denoted as made from cotton, hemp, linen, viscose and polyester was exposed in a mesocosm simulation experiment on marine mud. The end point, and in some case the intermediate status of the degree of disintegration over time was estimated from photographs of the samples, as a first information on the specific lifetimes of the articles in the respective test scenario. The materials and the different environmental conditions are described in table SI5 and SI7.

#### Experiment 4: Field experiment on the influence of temperature on cellulose biodegradation in a seasonal lake

To address temperature as the main driver of environmental biodegradation rate, the disintegration of cellulose filter paper was studied in a field experiment at the bottom of a freshwater lake (Adamsee, 48°43′41.9″ N, 8°4′57.7″) in the Upper Rhine Valley of south-western Germany at about 4 m water depth for over two years. The ambient temperature seasonally ranged between 5 °C and 26 °C. The rather sticky sediment consisted of sandy silt with a high organic content. Sediment properties were analysed with standard methods (see data not shown). Cellulose filter paper samples were prepared as described above according to Lott et al., 2020. One set of samples (n = 3 per interval) was laid on top of the lake sediment (fig. 9, blue columns) and another set was buried ∼15 cm below the sediment surface (fig. 9, red columns). Samples were retrieved approx. every two months and replaced by new samples. The retrieved samples were analysed for the degree of disintegration (see below). Temperature was recorded with automated temperature loggers (Tinytag) near the samples (see tab. SI7).

### 2.2 Geographical coverage of all experiments

Fig. 1 shows a map with the locations of the field experiments and the provenance of environmental matrices (soil, freshwater and marine sediments) on a global scale.

Tank experiments were performed at HYDRA labs, with soils from a garden, a forest and an agricultural field nearby, and with freshwater sediments from nearby lakes and a creek, and with marine sediments from beach locations at the German Wadden Sea, the Italian coasts of the Adriatic and the Tyrrhenian Seas, and from Singapore. In Norway, marine tank experiments were conducted in a flow-through tank of a public aquarium of Bergen.

Soil field experiments occurred at an organic farm (Biogärtnerei Schmälzle & Sohn, Sinzheim) and freshwater field experiments in ponds in Kassel (Germany), Casalgrande (Italy), and in lakes Adamsee and Weitenung, Bühl (Germany). Marine samples were exposed in the High Arctic in the Kongsfjorden off the coast of Spitsbergen, Svalbard, Norway in the water column at a depth of 91 m. In the Mediterranean Sea, the benthic marine field tests were performed in the protected area of the National Park Tuscan Archipelago off the island of Pianosa and the eulittoral and lagoon test sites were set up in a former salina on the island of Elba, Terme di San Giovanni, Portoferraio, Italy as described in Lott et al. (2020). In the tropical sea of Indonesia, the pelagic and benthic field tests were performed in Sahaong Bay, Pulau Bangka, NE Sulawesi. The eulittoral test site was a sandy beach protected by a rim of mangrove forest. Deep-sea tests were conducted in the Mediterranean Sea in the water column of the Sicily Channel at 532 m and the Corsica Channel at 400 m by attaching a stainless-steel frame to instrumented mooring lines used for permanent oceanographic monitoring, similar to the pelagic tests described in Lott et al. (2020).

### 2.3 Data analysis

#### Biodegradation analysis

Biodegradation was calculated from the raw data of CO_2_ evolution over time from respirometer tests conducted by HYDRA on a 12-channel flow-through respirometer (ECHO Instruments, Slovenia) as described by Kintzi et al. (2025). For experiments conducted in standard tests (see tab. 1) by the contracted laboratory (Normec OWS), biodegradation was assessed using either CO_2_ production or O_2_ consumption data, depending on data quality. Raw data was not available numerically for all experiments and if necessary, extracted from plots in test reports using WebPlotDigitizer (https://automeris.io).

#### Disintegration analysis

Disintegration of sheet samples (experimental sets 2, 3 and 4) was determined photogrammetrically from standardized images which were taken right after sampling as previously described (Eich et al., 2021). A convolutional neural network (U-Net model) was implemented using TensorFlow 2.12.0 (Abadi et al., 2016) and keras (Chollet et al., 2015) to perform semantic image segmentation, classifying each pixel as ‘hole’ or ‘no hole’. The neural network was trained using photos of samples along with corresponding masks, which were manually created in Affinity Photo (version 1.10.6). The masks consisted of color-coded regions marking the disintegrated area and the area with the remaining sample. The deep learning segmentation model provided a high-resolution pixel-wise classification image, from which the degree of disintegration (% area loss) was quantified.

#### Statistical analysis and calculation of half-life

The modeling of disintegration data of film samples from tank and field tests and biodegradation data from respirometer tests, as well as half-life (t_0.5_) calculations, were carried out using R (R Core Team, 2025) as described in Lott et al. (2021). For film samples deployed in mesocosm tanks and in the field, the disintegration over time was modelled using beta regression (*betareg* package) and the appropriate link-function (*logit, cloglog, cauchit*, or *loglog*) selected by comparing the Root Mean Square Deviation (package *caret*). Half-life was calculated by back-transforming link-functions for 50% material disintegration and solving the formula for the time (t_0.5_) using the coefficients estimated by the linear model. A reliable half-life estimation requires sufficiently high data quality. Treatments in which the last measured data point fell below 50%, and/or for which only endpoint measurements were available, resulted in highly uncertain predictions and were therefore considered to lack statistical robustness. For these cases, half-life calculations were not performed. Instead, the extent of final disintegration was either visualized separately or reported descriptively within the graphs of half-life plots.

The half-life of the materials deployed in respirometer tests was analyzed using Three Parameter Logistic Regression (3PL) by fitting the data with a non-linear model (package *nlme*) (formula adapted from Junker et al., 2016). To account for the lack of independence from repeated measurements of the sample in respirometer tests, replicate ID was included as a random factor in the analysis.

Monte-Carlo simulations were used to calculate 500,000 half-life values considering the variance of the model coefficients, which allowed statistical tests with empirical p-values that were adjusted for multiple comparisons using the method from Holm (1979). The distributions of the re-sampled half-life values were visualized by violin plots in which the spread along the *x* axis represents the frequency of values on the *y* axis (half-life). Different letters indicate significantly different groups.

#### Calculation of the Specific Surface Degradation Rate (SSDR)

Biodegradation of solid polymers occurs via enzymatic attack at the surface of the material leading to surface erosion (Lott et al., 2021). Specific Surface Degradation Rates (SSDRs) (Chamas et al., 2020) provide a normalized measure to numerically compare materials or exposure scenarios and is also used in life cycle assessments such as the MariLCA approach (www.marilca.org) as part of the effect factor for environmental impact.

In our study, SSDRs were calculated as *ν_d_* based on the initial diameter of the test item (*d*_0_, in µm), exposure time (*t*), shape to determine the power (*a*), and the relative mass loss over time (Δ*m*/*m*_0_), following (Maga et al. 2022). For infinite films, the power *a* equals 1, and for fibers, it equals 2. In accordance with their recommendation to estimate SSDRs at approximately 50% degradation, we derived SSDRs using the modelled half-lives of the test materials. To account for variability and uncertainty in our estimates, we further calculated empirical 95% confidence intervals for the SSDRs based on the Monte-Carlo simulations of modelled half-lives.

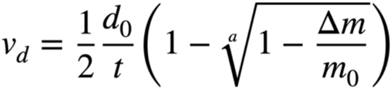

When classified as a “film”, following Maga et al. (2022), sheet-like fibrous materials showed unrealistically high SSDRs, suggesting that this classification did not adequately reflect the surface accessibility and biodegradation dynamics in these materials in contrast to the uniform body of a plastic film. In fibrous materials, individual fibers are exposed to microbial attack from all directions, unlike true films, where biodegradation occurs from only one or two exposed surfaces. We therefore reclassified all materials as a “fiber”-type, yielding more realistic and comparable SSDR values. However, biodegradation in sheet-like fibrous materials, such as nonwoven and woven fabric, is slower than in free fibers most probably due to some shielding effect and diffusion limitation within the interwoven structure.

For materials for which only the dtex value was available the fiber diameter was calculated using the following formula, where *d* is the fiber diameter, dtex the linear density in decitex (g/10,000 m) and *ρ* the density of the specific material (Viscose: 1.5045, Modal: 1.5141, Lyocell: 1.5205, Cotton: 1.5) in g/cm³.

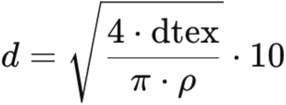

## 3. RESULTS

### 3.1 Standard biodegradability and disintegration lab tests in home compost, soil, water with activated sludge, freshwater and sediment, seawater, seawater and sediment

Biodegradability and disintegration tests were conducted according to international standard test methods (tab. 1) at temperatures between 21 ± 1 and 30 ± 2 °C. The degree of biodegradation in % was calculated at the end of the experiment from the conversion of polymer carbon into carbon dioxide (mineralization) or in some cases the amount of oxygen consumed. For the standard disintegration tests, a material was determined as completely disintegrated if after the termination of the experiment no particles larger than 2 mm were retained by sieving the content of the incubation vessel. Figure 2 shows that all materials had reached at least 70% of mineralization within 1 to 4 months. For the monospecific materials tested here, the carbon which is not balanced by the evolved carbon dioxide is assumed to have been used by the microbial community for built-up of biomass. Generally, the form and shape of the materials submitted to such tests influence the accessibility of the single fiber for biodegrading microbes thus modulate the biodegradation *rate* or the lifetimes of products, respectively. The use of a powder (microcellulose or cryomilled from fiber material) with a higher surface-to-volume ratio enhanced the biodegradation rate compared to the fibrous material as can be seen in the difference between the values for microcellulose (cyan dots, fig. 2) and cellulose filter paper (dark blue dots, fig. 2), and in the case of linen, the differentiation between loose fiber and a piece of woven textile (# and §, fig. 2). All tested materials, i.e. the different variants, forms, and qualities of cellulosic fibers, met the requirements for home compostability, soil, water, seawater, and marine biodegradability tests. The data from a repetition of independent experiments were largely consistent, with slight variations explained by differences in the test materials’ physical form (e.g., fiber thickness, loose fiber vs. nonwoven). All cellulosic materials were biodegraded similarly, with the regenerated cellulose variants of neat viscose, modal, and lyocell fibers being biodegraded as completely as the native cellulose forms filter paper and linen fiber.

**Figure 2:**
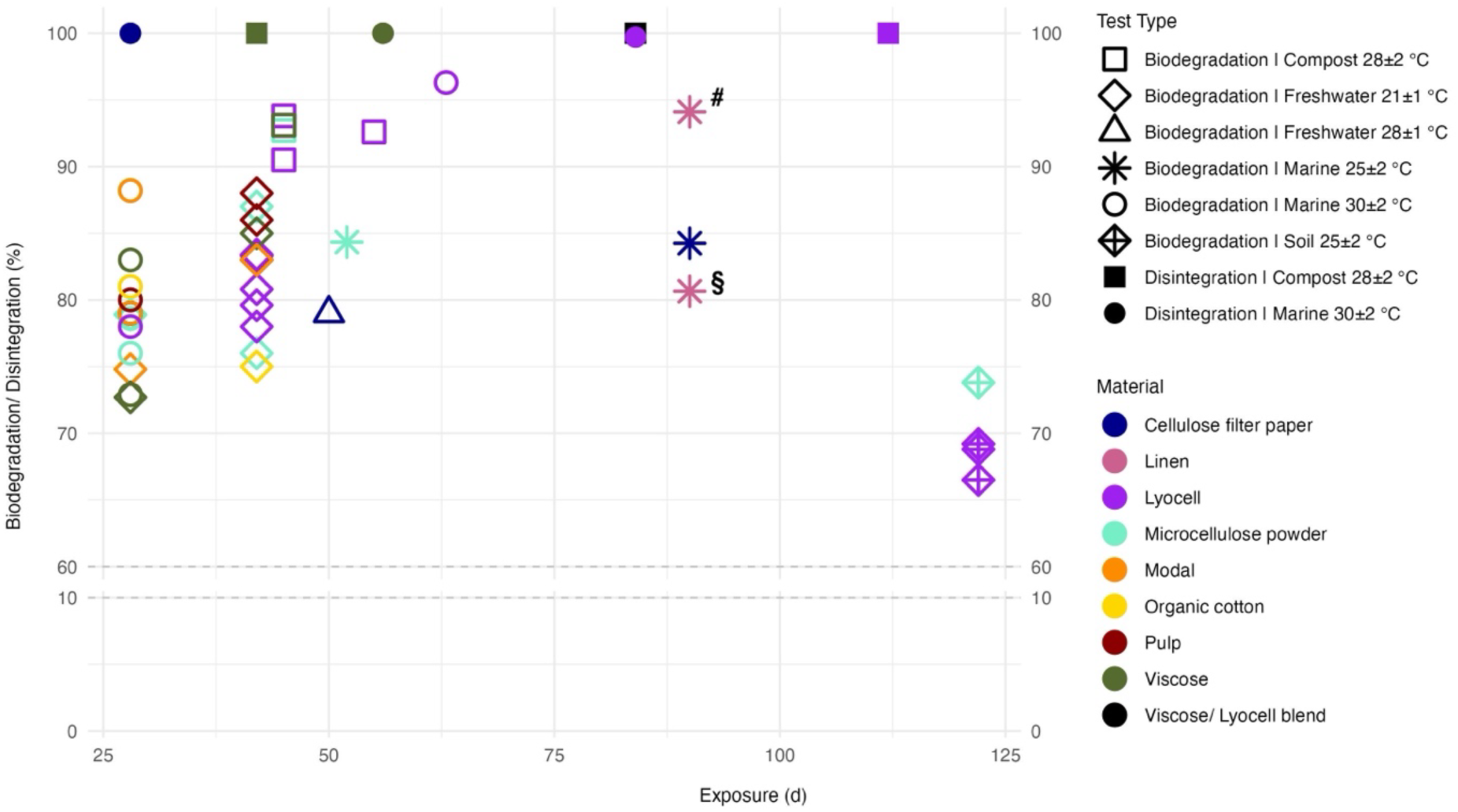
**Biodegradation and disintegration of different forms of native cellulose, linen, viscose, modal and lyocell in 57 standard laboratory tests** under various conditions: home compost 28 ± 2 °C, soil 25 ± 2 °C, water with sludge inoculum 21 ± 1 °C, freshwater 28 ± 2 °C, marine sediment 25 ± 2 °C and seawater 30 ± 2 °C. Biodegradation is given as degree of mineralization at the plateau phase at the end of the tests. Disintegration values in % are given as the end-point of the test duration (filled dots). The plot shows the data coded by color for material type and by shape for the test scenario. # loose linen fibers, § linen fabric.

Although it is known that temperature generally matters for the rate at which biodegradation processes occur there was no clear trend in the comparative overview of different lab test scenarios. An effect of an assumed natural heterogeneity of the matrices water, soil, sediment used in the various tests could not be differentiated from a possible temperature effect.

### 3.2. Environmental relevance: Cellulose filter paper in mesocosm and field tests

Different environmental scenarios were experimentally assessed for the biodegradation of cellulose filter paper which is being used as a positive control and reference material in many biodegradation studies. In our study, the disintegration as a proxy for biodegradation of ash-free cellulose filter paper was determined in time series of several standard tests and adaptations thereof. The data was statistically modeled to obtain a specific half-life (Lott et al. 2021) for each combination of material and test scenario. The specific half-lives allow to numerically compare the biodegradation behavior of a specific object under different conditions, or different objects under the same conditions and rank them.

#### 3.2.1 Compost experiments

The tests conducted according to the standard for industrial compostability (*EN ISO 14855 – Determination of the ultimate aerobic biodegradability of plastic materials under controlled composting conditions*) were kept at a lower temperature of 28 ± 2 °C to simulate home composting conditions. The moisture content of the matrix was controlled as prescribed in the standard. The half-lives modeled from the original mineralization data of lab biodegradability tests were about 1 to 2 weeks for microcellulose powder and 3 to 4 weeks for cryomilled lyocell or viscose. The matrix consisting of a mix of green compost and vegetable, garden and fruit waste (VGF in fig. 3) seemed to have led to slightly higher biodegradation rates than inocula denoted as stabilized and mature compost derived from municipal solid waste (MSW in fig. 3). However, replication was too low for statistical analysis (n = 1), and the variation was most likely due to the different particle sizes of commercially procured microcellulose on the one hand, and lyocell and viscose that were individually cryomilled in the lab on the other.

**Figure 3:**
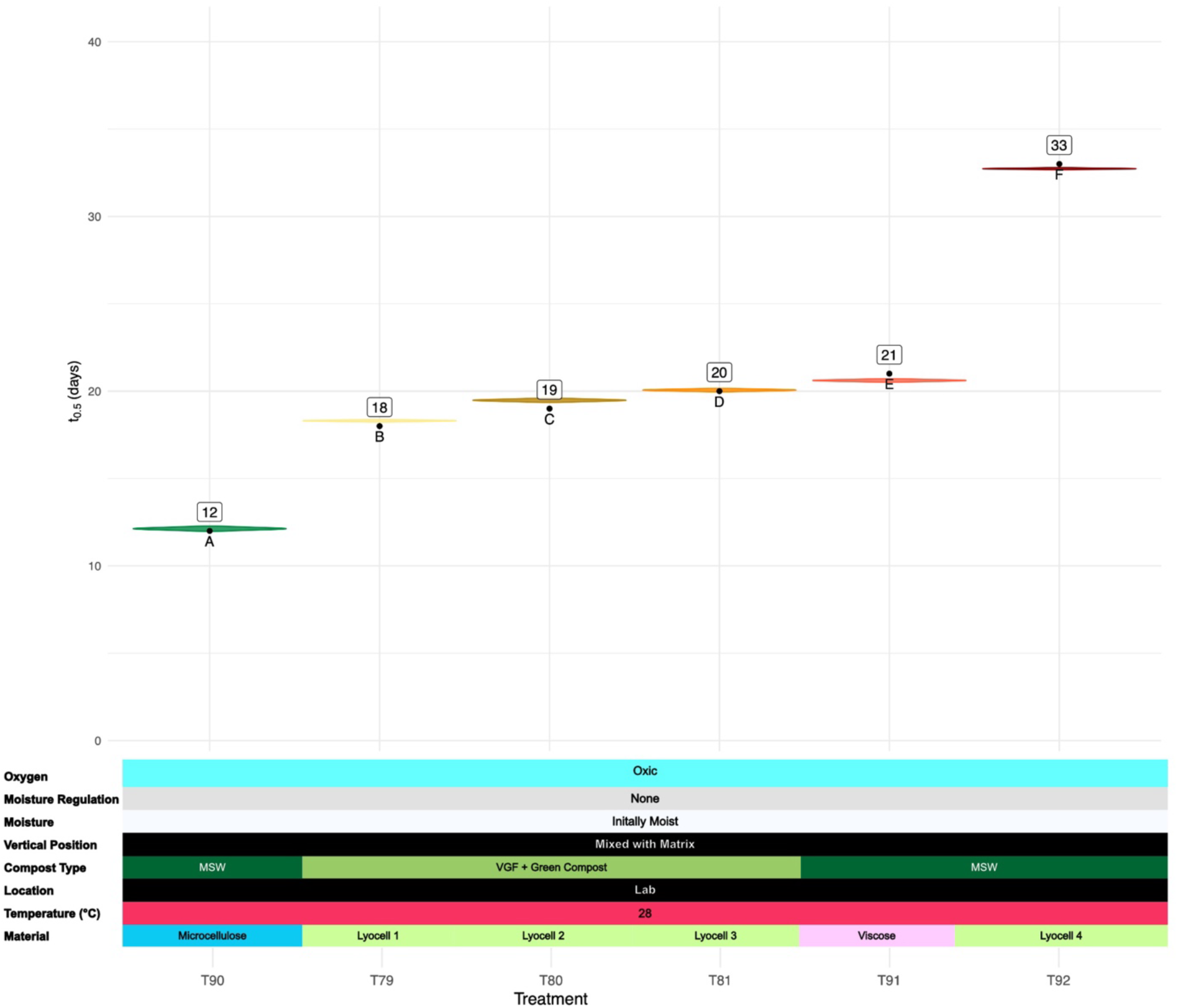
**Biodegradation half-lives of microcellulose, viscose, modal and lyocell**, all in powder form, in 7 standard home compostability tests (EN ISO 14855) at 28 °C. VGF = vegetable, garden and fruit waste, MSW = stabilized and mature compost derived from the green waste fraction of municipal solid waste.

#### 3.2.2 Soil experiments

Cellulose filter paper was exposed for disintegration in tank and field experiments, laid on the soil either openly (surface) or covered with silage (fermented hay) or buried in the soil, in the open field or in a greenhouse tunnel. In most of the 24 scenarios tested, cellulose filter paper was disintegrated within weeks to months (fig. 4). The fastest disintegration with a modelled t_0.5_ of 23 d was measured for the T40 samples which were buried in the open field, with rainfall occurring during the exposure period at temperatures between 2.6 and 28.3 °C. The slowest disintegration with sufficient data for modelling the half-life (t_0.5_ = 245 d) was measured for T38, where samples were exposed in a greenhouse tunnel on the surface of agricultural soil covered with silage, away from the reach of the drip irrigation at a temperature range from -2.2 °C to 31.6 °C. In very dry soil treatments, disintegration was very low to not measurable within the exposure time of up to 621 days (T21-T23, T27, T28b, T35, grey boxes, fig. 4). While the driest samples were still visually unchanged and clean white, in some samples a strong discoloration with orange, brown and black spots, similar to what was frequently observed at the onset of disintegration in faster-degrading samples, indicated the presence of microbes presumably linked to biodegradation activity. Water availability was likely to be the most important determining factor for biodegradation-driven disintegration of cellulose filter paper in soil settings. Temperature seemed to play a less important role, and high temperatures alone did not lead to a higher disintegration rate or shorter half-life. The addition of compost tea (T31, T32, T43, fig. 4) accelerated biodegradation.

**Figure 4:**
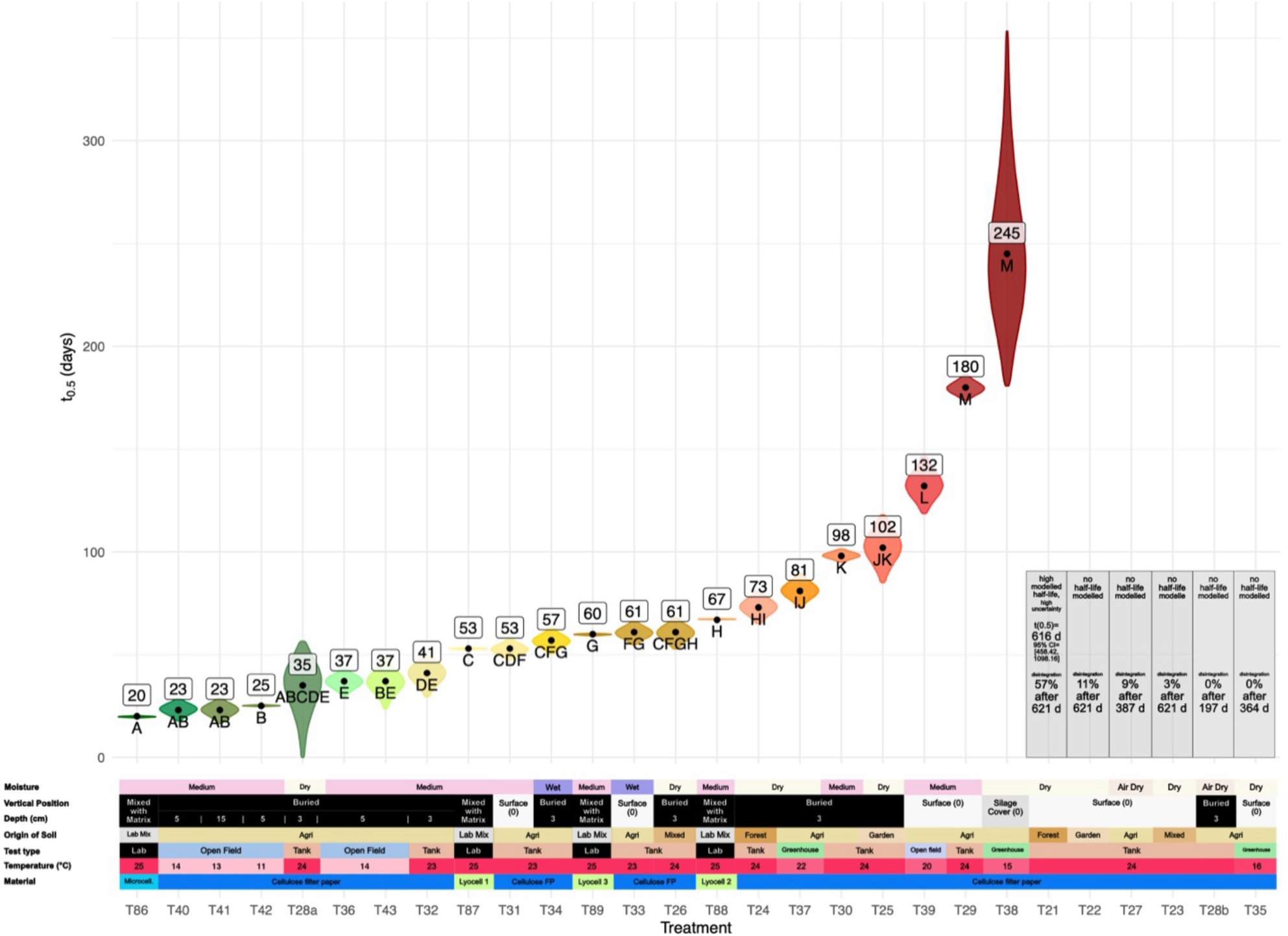
**Soil disintegration half-lives of cellulose filter paper compared to biodegradation half-lives of lyocell** (T87 - T89, marked with light green bar). Values for cellulose filter paper modelled from disintegration experiments at a constant temperature of 23 or 24 °C in mesoscom tests (tank) and ambient temperatures in field tests under open conditions with mean temperatures from 11 °C to 22 °C and fluctuations from -2.1 °C to 54.1 °C and in the unheated greenhouse from -6.1 °C to 47.7 °C. Values for lyocell were modelled from biodegradability standard laboratory tests at 25 °C.

#### 3.2.3 Freshwater experiments

Cellulose filter paper was exposed for disintegration tests in tank and field experiments at mean temperatures between 12 °C and 28 °C (min. 2 °C; max. 32.9 °C). Samples were laid on the sediment-water interface (benthic scenario) or buried in sediment (buried). Sediments were collected from a creek and two lakes in southwestern Germany. Field experiments were conducted in one of the lakes at 4 m water depth. In all tested freshwater systems, in or on lake and creek sediment, biodegradation was occurring and the disintegration half-lives of cellulose filter paper ranged from a few weeks (T10: t_0.5_ = 25 d; fig. 5) to some months (T2: t_0.5_ = 250 d; fig. 5). Besides temperature, the sediment property seemed to be the most inportant factor influencing the biodegradation rate. When buried in the sediment, the half-life of cellulose filter paper was slightly shorter than when laid on the sediment surface. Our field studies revealed that the seasonal variation of temperature led to variable disintegration rates over the course of the year. This was subsequently addressed with a dedicated experiment (see below, fig. 9).

**Figure 5:**
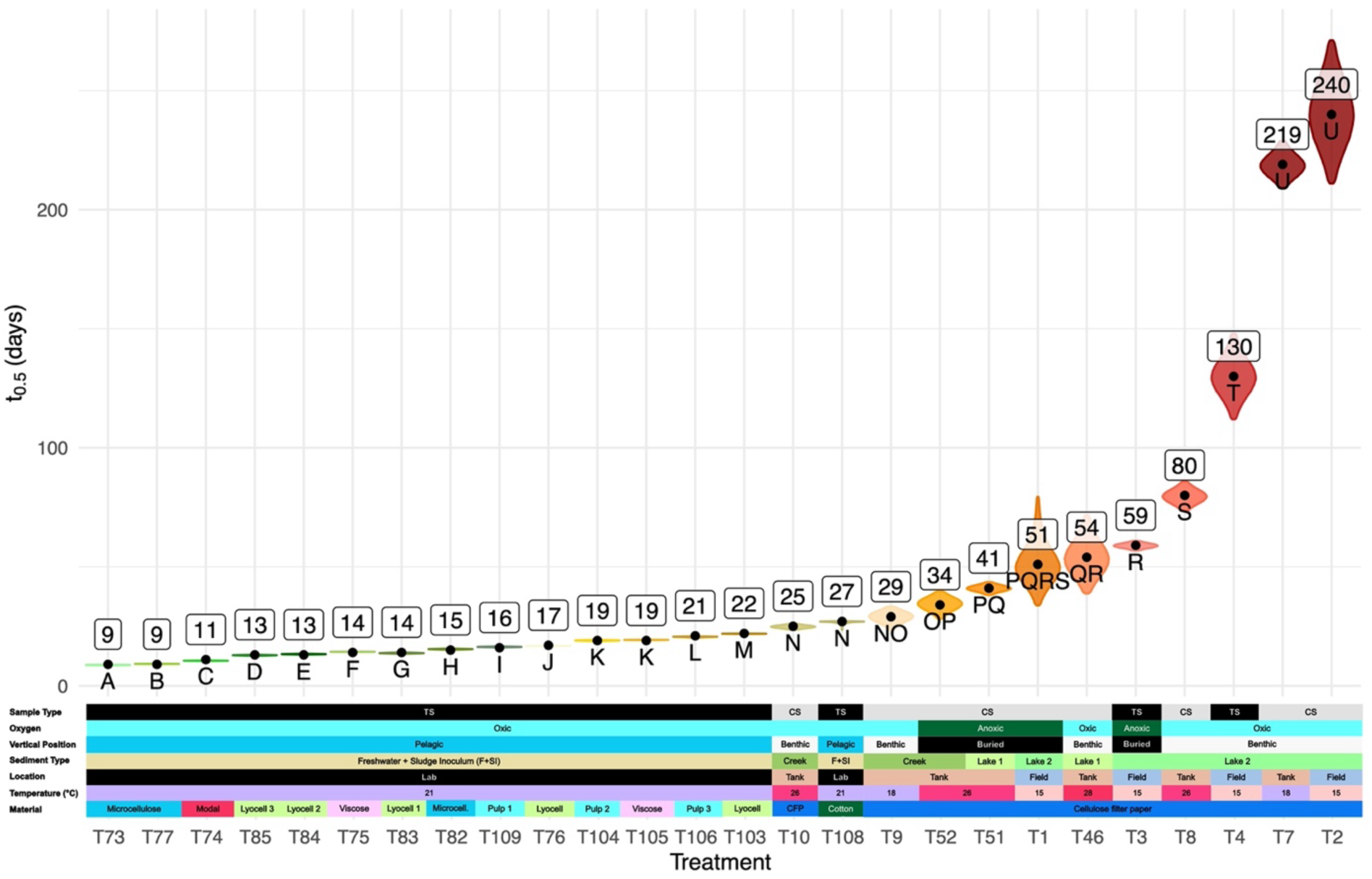
Freshwater disintegration half-lives of cellulose filter paper (CFP, dark blue) compared to biodegradation half-lives of modal (T74, red in ‘Material’, bottom line), lyocell (T76, T83-T85, T103, light green), viscose (T75, T105, pink), paper pulp (T104, T106, T109, light blue) and cotton (T108, dark green). Values for cellulose filter paper were modelled from disintegration at temperatures between 18 °C and 28 °C in mesoscom tests with sediment and water from two lakes and one creek, and at ambient temperatures (4.8 – 25.8°C) in a field test in a lake. Values for microcellulose, modal, lyocell, viscose, paper pulp and cotton were modelled from biodegradability standard laboratory tests with activated sludge-amended freshwater (F+SI) at 21 °C.

#### 3.2.4. Marine experiments

Cellulose filter paper was exposed for disintegration in tank and field experiments at mean temperatures between 9 °C and 29 °C (min. -1.8 °C; max. 38.4 °C). Samples were either pending in the open seawater (pelagic scenario), laid on the sediment-water interface (benthic scenario) or buried in sediment (beach or anoxic). Sediments used for tank tests were from beaches from a Norwegian fjord, the German North Sea, the Mediterranean Sea and from tropical Southeast Asia. Field experiments were conducted in the High Arctic ocean, under tropical coastal conditions in Indonesia in a sandy beach, in the water column and on the seafloor at 32 m, and in the Mediterranean Sea in a sandy beach, on the seafloor at 40 m water depth, and in two deep-sea settings.

Cellulose filter paper was biodegraded under marine conditions (fig. 6) comparable to freshwater or soil test scenarios, in contrast to a widespread belief that the marine environment is low in biodegradation activity towards polymers. The shortest half-life of only 15 days was measured with sand from a North-Sea beach in a sediment-water interface test in a mesocosm tank at 23 °C, a temperature comparable to the *in-situ* summer maximum. Even at a low temperature of 9 °C cellulose filter paper tested in a public flow-through aquarium in Norway on a sand surface had a disintegration half-life of 58 days. A half-life of 389 d (∼13 months) was modeled from T50 in an experiment with sediment ultralow in nutrients at 14 °C. Surprisingly, although run at 23 °C, tests with three beach sediments (T13, T14 and T16, grey boxes, fig. 6) showed a very low biodegradation activity towards the cellulose filter paper and the data was not sufficient to model disintegration half-lives after an experiment duration of almost 7 months (202 d). Although no disintegration could be measured within the observation time, the discoloration of the samples with brown and black spots, similar to the observations in the soil experiments, indicated the presence of microbes and the onset of biodegradation. Sediment quality seemed to be a strong determinant of the biodegradation activity, leading to a great variability of specific half-lives, as it was also shown before for biodegradable plastic polymers (Eich et al. 2021). The chemical analysis of the sediments in T13, T14 and T16 did not show any unusual properties (e.g., nutrient concentration, organic matter content, toxic chemicals etc., data not shown) to give any hints as to what could have caused such low activity. Further studies on the microbial community are pending and were not subject to this study. One sediment (NOR4, fig. 6) showed an almost 5 times higher biodegradation rate with t_0.5_ = 34 d if tested under oxic conditions at 23 °C (T20) than in the absence of oxygen at 26 °C (t_0.5_ = 166 d; T53).

**Figure 6:**
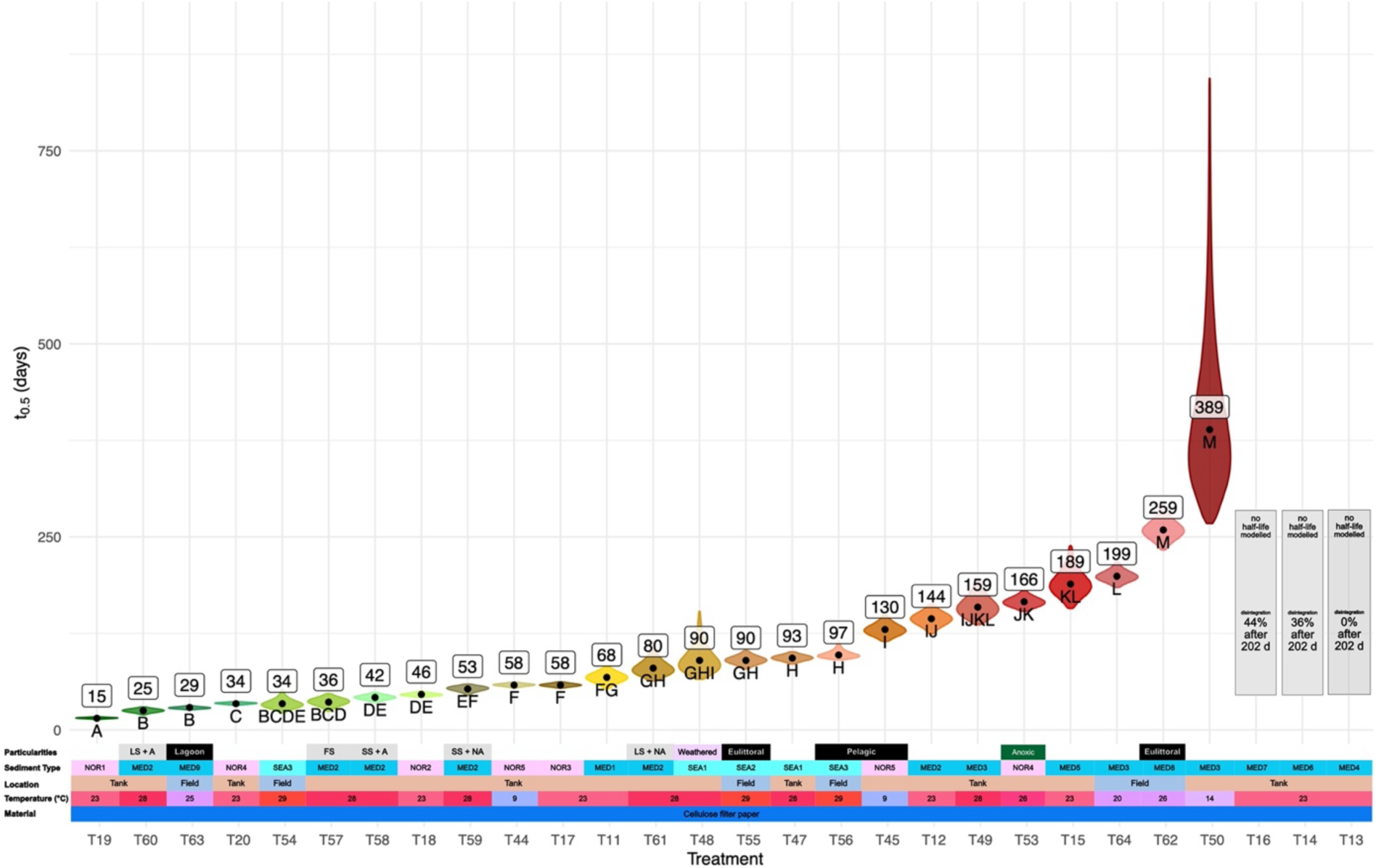
**Marine disintegration half-lives in field and tank tests**, of cellulose filter paper modeled from disintegration experiments in mesoscoms with sediment and water from the North Sea, the Mediterranean Sea and from tropical Southeast Asia, and in field tests at mean temperatures between 9 °C and 29 °C (min. -1 °C; max. 38.4 °C). T53 shows data from a mesocosm test with beach sand kept free of oxygen (anoxic).

As a comparison to results from the open-system disintegration tests in field and mesocosms, fig. 7 shows the half-lives modeled from the mineralization in standard laboratory tests with seawater (30 ± 2 °C) or at the seawater-sediment interface (26 ± 1 °C). Most of the treatments resulted in half-lives below 2 weeks, and treatments of pulp varieties, microcellulose, modal, viscose and cotton, with seawater and nutrients added, showed half-lives of only 7 days or less. The deviation of the half-lives of the same linen fiber material by a factor of 3 could be explained by the form of the material. For loose linen fibers of 9 µm diameter t_0.5_ was 11 d (T110, fig. 7) and for an intact piece of woven linen fabric of ∼335 µm thickness t_0.5_ was 34 d (T112, fig. 7). Exemplarily, photo panels with time series from a tropical beach experiment (fig. SI18), in the tropical water column (fig. SI19) and at the tropical seafloor (fig. SI20) are shown in the Supplementary Information.

**Figure 7:**
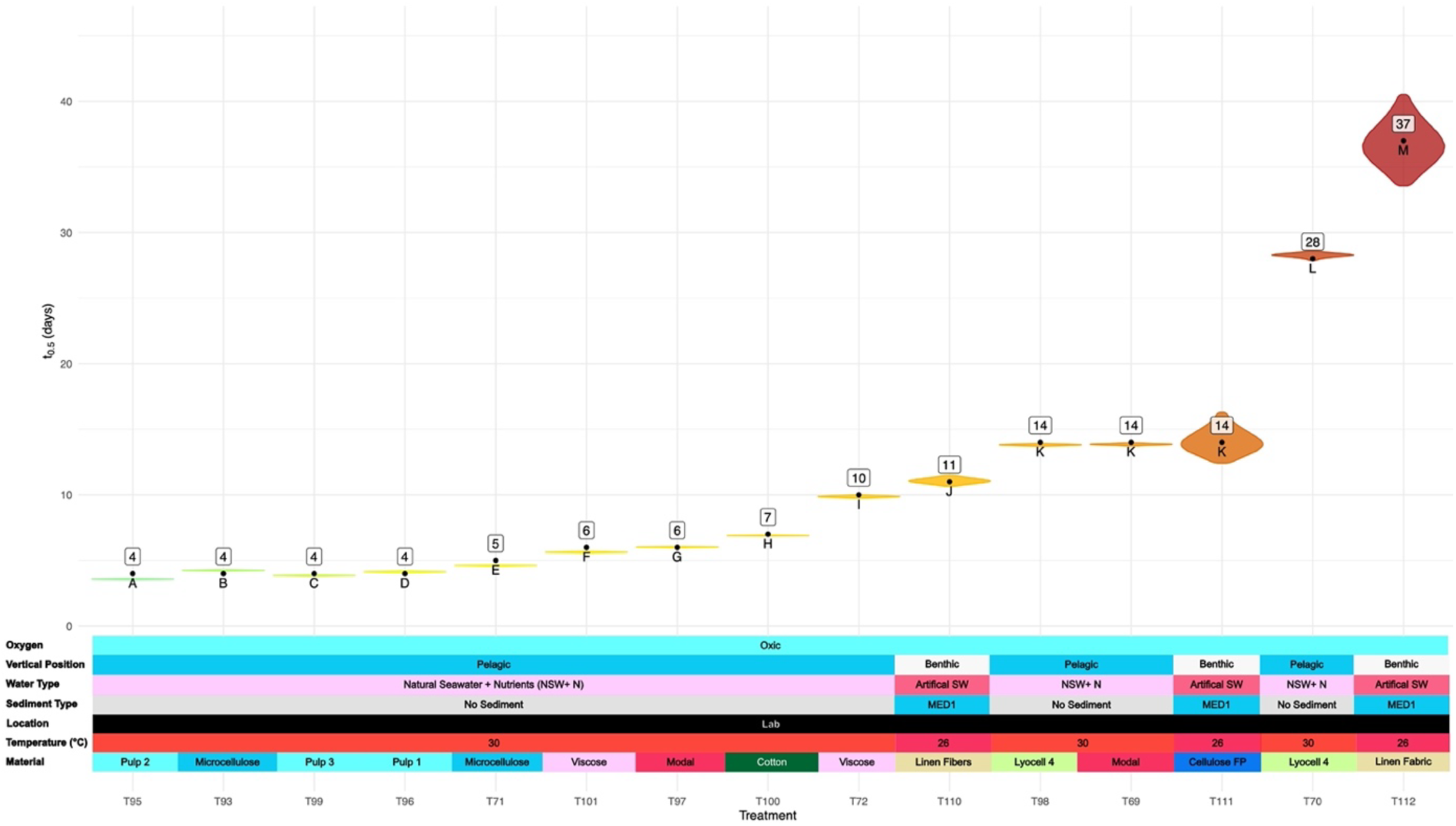
Marine biodegradation half-lives of microcellulose (T93, T71), cellulose filter paper (T111, dark blue), paper pulp (T95, T96, T99, light blue), modal (T69, T97, red), lyocell (T70, T98, light green), viscose (T72, T101, pink), cotton (T100, dark green) and linen fibers (T110,T112, beige). Values were modelled from standard biodegradability tests with seawater at 30 °C (pelagic) or with seawater and marine sediment (benthic) at 26 °C.

### 3.3 Specific Surface Degradation Rates (SSDR)

In addition to the half-lives for the fibrous materials also the SSDRs were calculated from the biodegradation and disintegration data (tables SI1-SI4). In home compost most tests were done with powder instead of fibers and thus only a few SSDRs could be calculated. The highest SSDR was measured for viscose fibers of 12 µm diameter in home compost at 28 ± 2°C with a rate of 0.0836 µm d^-1^ (no CI, 1 data point only), and a deduced t₀.₅ = 21 d. Cellulose filter paper fibers of 20 µm diameter were eroded in soil at 26 ± 2 °C with the highest surface-specific degradation rate (SSDR) of 0.127 µm d⁻¹ (CI: 0.102–0.177 µm d⁻¹) and a half-life (t₀.₅) of 23 d (CI: 16.52–28.74 d). In freshwater at 26 °C, the highest SSDR was 0.118 µm d⁻¹ (CI: 0.112–0.124 µm d⁻¹) with t₀.₅ = 25 d (CI: 23.54–26.13 d). Under marine conditions at 25 ± 1 °C, the highest SSDR reached 0.209 µm d⁻¹ (CI: 0.180–0.236 µm d⁻¹) with t₀.₅ = 14 d (CI: 12.41–16.29 d).

The slowest measurable SSDRs for cellulose filter paper were 0.00474 µm d⁻¹ in soil (CI: 0.00263–0.00638 µm d⁻¹), 0.0122 µm d⁻¹ in freshwater (CI: 0.0108–0.0139 µm d⁻¹), and 0.00753 µm d⁻¹ in marine tests (CI: 0.00337–0.0109 µm d⁻¹).

### 3.4. Observational marine disintegration experiments *in situ*

The disintegration of fabric samples of cotton and lyocell compared to cellulose filter paper, of an organic cotton t-shirt, denim blue jeans, wood (oak veneer) and of banana leaves was tested, with 1-3 replicates and one sampling event only, under coastal tropical marine conditions (Pulau Bangka, Sulawesi Utara, Indonesia) and of fabric samples in the High Arctic, in the Mediterranean Sea in a coastal and in a deep-sea setting (Sicily Channel, 532 m depth and Corsica Channel, 400 m depth).

In the Arctic water column (−91 m, T = -1.8 - 6.8 °C) cotton t-shirt samples were almost completely (> 98 %) gone after ∼2 years, whereas veneer sheets of oak wood were still largely intact after ∼3 years in the same setting. In both deep-sea experiments in the Mediterranean Sea water column, at 400 and 532 meters water depth respectively, cellulose filter paper, cotton and lyocell fabric were also completely disintegrated after 20 to 24 months (fig. 8), and there were no material residues visible (fig. SI11). In the coastal Mediterranean Sea, complete disintegration was reached in the lagoon mud after 9 months, on the seafloor at 40 m water depth after 14 months, and in the beach sand after almost 2 years. The woven cotton and lyocell samples disintegrated slightly slower than the cellulose filter paper, supposedly due to their denser geometry. Note the imprint of the fiber structure in the cotton and lyocell samples in the eulittoral experiment after 673 d, likely due to intense biofilm formation (fig. SI12, left column).

**Figure 8:**
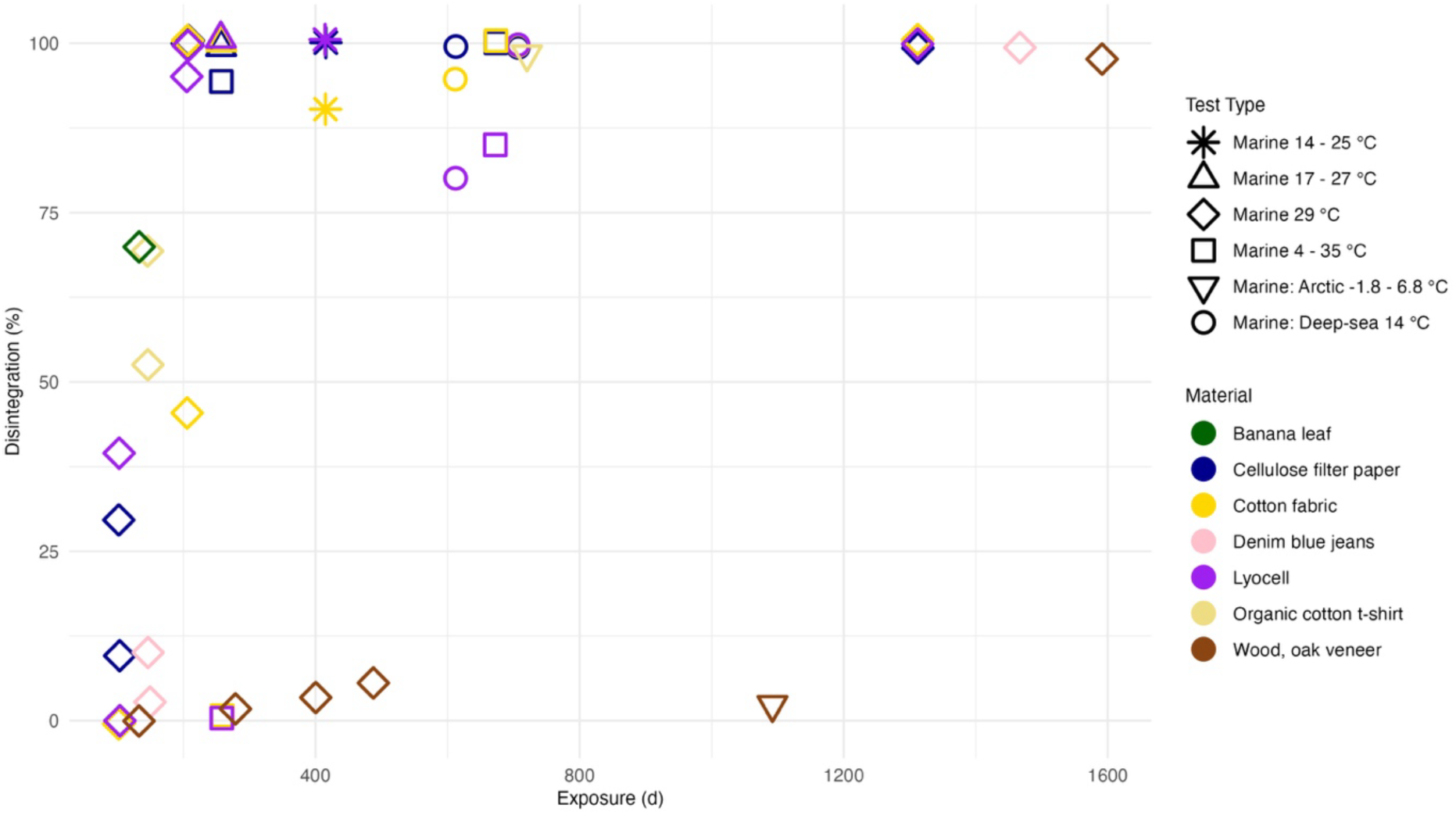
Marine disintegration of cellulose filter paper, cotton and lyocell fabric, cotton t-shirt, denim jeans, banana leaves and wood. Observational experiments with few replicates (n=1-3) in deep-sea and shallow-water settings. The 44 data points show the intermediate or endpoint degree of disintegration. The 100% values represent the maximum lifetime under the respective conditions and were most likely reached months earlier, but due to accessibility constraints samples could not be recovered in time. For details see table SI5.

In the tropical marine coastal settings of Indonesia, in the beach experiment, complete disintegration was reached for all three materials after 207 d, whereas at the seafloor in 32 m water depth, a substantial amount of the cotton fabric and few remains of the lyocell fabric were still left, supposedly due to their denser geometrical structure compared to the filter paper (fig. SI13). The sediment in the beach experiments showed higher activity towards cotton and lyocell than in the seafloor experiments. Disintegration of organic cotton t-shirt samples was far advanced but not completed yet after 5 months in both the benthic and eulittoral setup and thin remains of the fabric were still found. Visually, it was hard to distinguish between biofilm and fiber residues after 146 days (fig. SI14). For the Denim blue jeans samples, disintegration was noticeable but not completed yet after 5 months in the eulittoral and the benthic scenario. The seafloor test was terminated after 149 days, and the beach samples were redeployed. After a total of 4 years, only the fine protective gauze originally placed around the samples remained, and no trace of the test material was left in the eulittoral (fig. SI15). The wood samples from an oak veneer (380 µm thick) only started to visibly disintegrate after 13 months on the sea floor. After 4.4 years, a few fragments of the material were still left (fig. SI16). After the exposure of 4.5 months on the seafloor at 32 m depth, fresh banana leaves were strongly but not yet completely disintegrated (fig. SI17).

### 3.5. Temperature-dependence: Seasonality of cellulose biodegradation in a freshwater lake

Disintegration of cellulose filter paper was studied in a lake in southern Germany at ambient temperatures ranging seasonally between 4.8 °C and 25.8 °C. One set of samples was laid on top of the lake sediment (fig. 9, blue columns) and another set was buried ∼15 cm below the sediment surface (fig. 9, red columns) under apparently anoxic conditions. After an exposure time of approximately two months, samples were replaced and the retrieved samples were analysed for the degree of disintegration. In the warmer period of the year, the 2-month-disintegration had reached over 90% in the buried samples and above 80% in the samples on top of the sediment. In the cold season, disintegration of the buried samples was between 3 and 44 % but the samples laid on the sediment were found still intact after 2 months. Clearly, temperature was the driving factor for the biodegradation of cellulose filter paper in this field experiment. The disintegration was faster for buried samples than for samples at the sediment surface.

**Figure 9:**
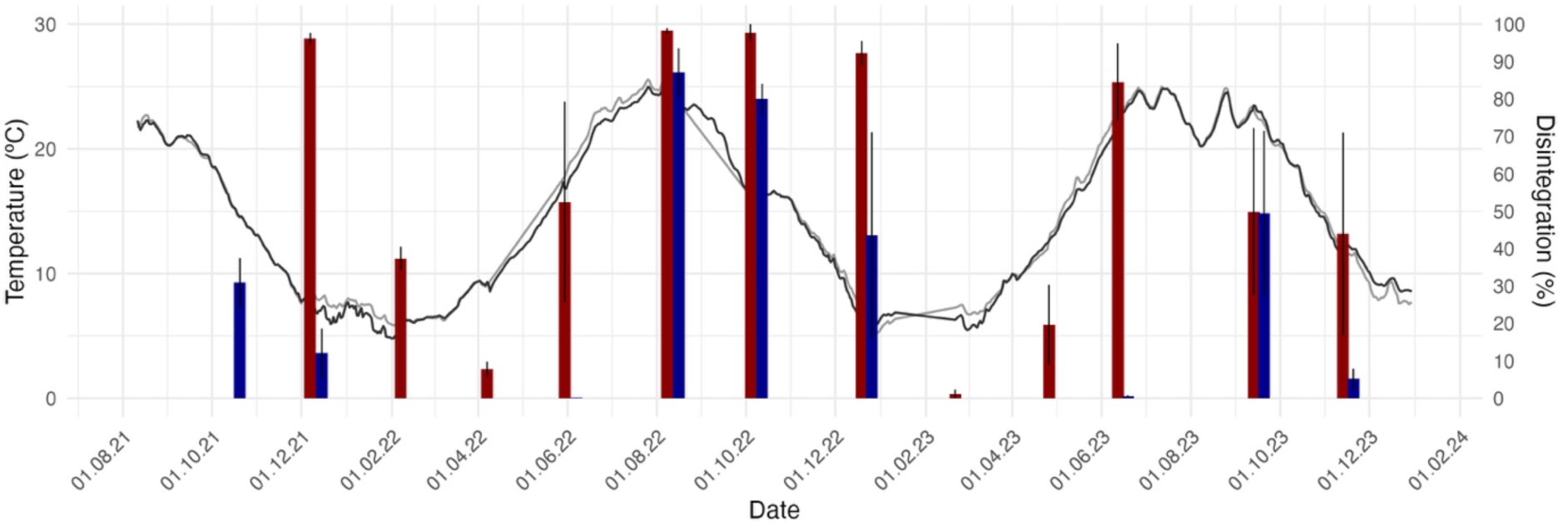
**Freshwater disintegration** of cellulose filter paper in a lake in southwestern Germany at ambient temperatures ranging seasonally between 4.8 °C and 24.8 °C in two scenarios: Samples (n = 3 per interval) laid on top of the lake sediment (blue columns) and samples buried ∼15 cm below the sediment surface (red columns), repeatedly exposed and retrieved after approx. 2 months. Error bars indicate the standard deviation. Temperature profiles from 2 independent sensors.

## 4 Discussion and Conclusion

### 4.1 The biodegradability of cotton, linen, cellulose filter paper and regenerated cellulose

All cellulosic materials examined in this study were biodegradable and showed gradual or complete biodegradation within the observation period under all tested environmental conditions. However, even pure cellulose filter paper showed a limited degree of biodegradation in certain soil, freshwater and marine settings within the duration of the experiments. This is a good example for the difference between the intrinsic material property of biodegrad*ability* as a potential and the actual behavior under natural conditions. Only if the conditions are favorable, the potential can unfold and the process of biodegrad*ation* occurs. The biodegradation *rate* strongly depends on the conditions as well and can be too low to be measured within the time frame of the experiment. As an example, in experiment 4, which examined the temperature-dependence of biodegradation in a freshwater lake, disintegration during the colder months was so minimal that for some samples it could not be reliably resolved with the applied method. A longer exposure under the same conditions would have been necessary, but this was incompatible with the experimental design. Modeling of the data with an adapted Arrhenius equation (fig. 10) suggests a possible threshold temperature below which no biodegradation occurs. However, this interpretation should be made cautiously, and more sensitive methods capable of detecting very low degradation rates would be needed to determine whether such a threshold truly exists and, if so, where it lies.

**Figure 10:**
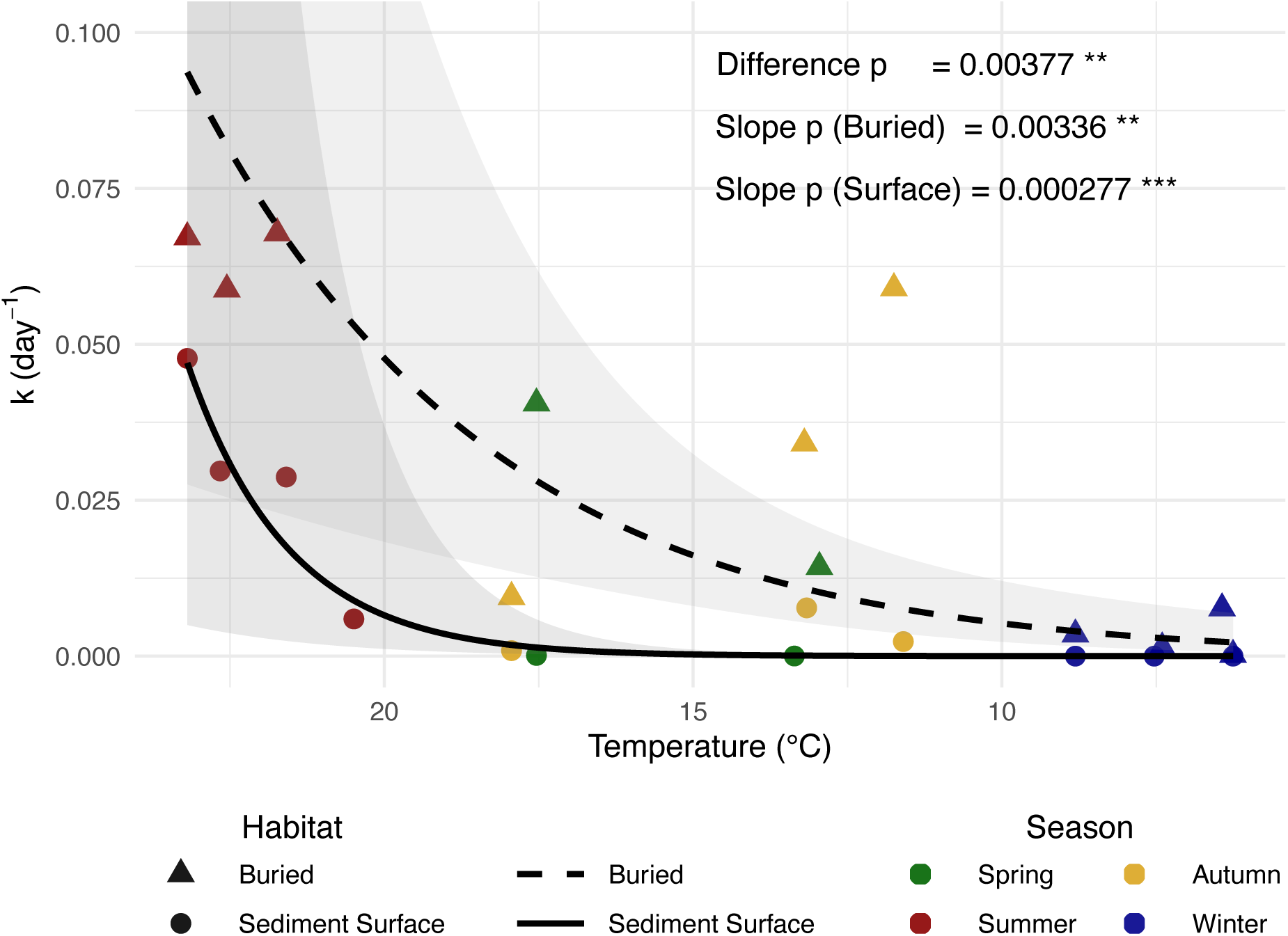
Temperature dependence of the daily disintegration rate of cellulose filter paper in a seasonal freshwater lake. Data points give the daily mean disintegration rate. The x-axis shows the sum of the daily mean temperatures. The data for the samples buried in the lake mud (triangles, dashed line) were modeled separately from the data for the samples laid on top of the sediment (dots, full line). Blue: winter months, green: spring, red: summer, yellow: autumn.

Figure 10 shows the same data from fig. 9 converted into a daily disintegration rate (% d^-1^) relative to the mean daily temperature. The daily mean temperatures were summed up over the whole exposure time. There is a highly significant correlation (p buried: 0.00336; p surface: 0.000277) between the mean daily temperature and the daily disintegration rate.

Although the temperature dependence of cellulose biodegradation in a freshwater lake could nicely be demonstrated in our experiment for both scenarios, different biodegradation rates between samples laid on and the samples buried in the sediment were expected but could not be explained by temperature differences. That the samples within the sediment were significantly (p = 0.00377) faster disintegrated than the samples laid on the sediment surface was counterintuitive. The mud of the lake floor was very likely free of oxygen at the depth where the samples were buried (15 cm), deducing from tank experiments done in another context with the same sediment (data not shown). The lack of oxygen within aquatic sediments is widely recognized as a constraint on rapid biodegradation, as anoxic conditions slow organic matter mineralization and promote carbon preservation in sediments (Canfield 1994). This biogeochemical control is also reflected in the archaeological record, where reduced oxygen availability in aquatic muds contributes to the long-term preservation of artifacts (Bengtsson 1975; Gregory 1995; Caple and Dungworth 1998), see also below. As a potential explanation for the faster disintegration of the buried samples in experiment 4, we speculate that the readily biodegradable cellulose introduced into deeper sediment layers was rapidly metabolized by a microbial community that, while enzymatically capable, is typically constrained by limited availability of labile organic carbon. In lake sediments, the vertical distribution of organic matter is strongly structured by sedimentation, with fresh, bioavailable carbon concentrated at the sediment–water interface and progressively depleted with depth due to prior microbial processing (Canfield, 1994; Middelburg, 2018). As a result, microorganisms in deeper sediment layers often experience carbon limitation despite remaining metabolically active (Arndt et al., 2013).

The artificial introduction of cellulose at depth may therefore have relieved this limitation by providing a high-quality substrate, triggering enhanced microbial activity and accelerated degradation. Such a response is consistent with concepts of carbon limitation and positive priming, whereby the addition of labile organic matter stimulates microbial metabolism and extracellular enzyme production, leading to rapid substrate turnover (Kuzyakov et al. 2000; Bengtson et al. 2012; Bianchi and Ward 2019).

In contrast, at the sediment surface, microbial communities are continuously supplied with diverse and abundant organic substrates derived from plant litter, benthic production, and sedimented planktonic debris (Middelburg et al. 1997; Burdige 2007). Under these conditions, the cellulose of the experimental samples may have represented only one of many available carbon sources and therefore experienced lower relative degradation rates due to substrate competition and reduced selective pressure for cellulolytic activity. Together, these observations highlight how depth-dependent carbon availability and priming dynamics can strongly modulate the rate of degradation of biodegradable materials in sediments, independent of their intrinsic biodegradability.

Cellulose occurs in different structural forms. Native cotton fiber consists mainly of cellulose I, but it can be converted into cellulose II, for example by mercerization, the treatment with concentrated aqueous sodium hydroxide (NaOH) to improve functional properties. Regenerated cellulosic fibers (RCFs) such as viscose, modal, or lyocell are composed of cellulose II, which results from the dissolution and subsequent regeneration of native cellulose. Despite these structural and processing differences, our results showed no systematic differences in biodegradation behavior between natural fibers (e.g., linen), processed native cellulose fibers (e.g., filter paper, cotton fabric), and regenerated cellulose fibers. Consequently, distinguishing between regenerated cellulosic fibers and natural cellulose in terms of environmental behavior appears unwarranted, and after mercerization also not possible.

From a mechanistic perspective, the consideration of “biodegradation” should not be equated simply with disintegration or reduced to compliance with pass levels in standardized tests, as these may not always capture environmental relevance. Instead, whether and at which rate biodegradation occurs depends on a combination of fiber structure, treatment history, and environmental conditions consistent with laboratory findings (Park et al., 2004; Zambrano et al., 2020). Environmental samples from deep-sea sediments often show high abundances of colored cellulosic fibers, indicating anthropogenic origin (Sanchez-Vidal et al., 2018; Adams et al., 2021). Fibers used in clothing undergo multiple chemical and structural treatments during manufacture (Lam et al. 2012; Zanchettin et al. 2023), which can increase their resistance to biodegradation. Processes such as mordanting, dyeing, coating, or antimicrobial finishing can limit enzyme access to polymer chains and may account for the persistent occurrence of cellulosic fibers in environmental samples, the so-called “cellulose enigma” (Suaria et al. 2020; Lott et al. 2022; Pasterk et al. 2024). An illustrative example is the case of early denim blue jeans recovered from a deep-sea shipwreck sunk in 1857, which were preserved for over a century and a half. Their remarkable state of conservation has been attributed not to the cellulose fibers themselves, but to zinc salts used as mordants in the dyeing process (Chen and Jakes 2001). Such cases show that the persistence of fiber products cannot be understood without considering the chemical and structural modifications applied during manufacture (Lykaki et al. 2021).

### 4.2 Standardized lab vs. mesocosm and open field tests

Standardized laboratory tests for assessing biodegradability are often criticized for being optimized and not reflecting the real scenario in the open environment nor testing a product as is (SAPEA 2020). This criticism seems to ignore the actual purpose of such tests. Optimized, controlled and standardized conditions are essential for ensuring comparability and reproducibility of results across laboratories and geographic regions. The purpose of standardized tests is to provide proof of biodegradability as an inherent property, demonstrating that the polymer can, in principle, be biodegraded by naturally occurring microorganisms. In contrast, environmental biodegradability is a system property determined by the specific conditions of the habitat to which the object is exposed. It is therefore advisable to always specify the relevant environment alongside any biodegradability claim, for example, “biodegradable in soil” or “biodegradable under marine conditions.”

Our study also showed that the biodegradation *rate* for the exact same material can differ by orders of magnitude depending on environmental conditions. Nonetheless, if a polymer such as cellulose is inherently biodegradable, it will biodegrade completely under most natural conditions. Only under extreme circumstances, such as in the absence of liquid water in hot or cold deserts, can biodegradation be so slow or cease that cellulose persists for long periods.

An example is 33,000-year-old dyed flax fibers found in a dry cave in Georgia (Kvavadze et al. 2009). Another example concerns fragments of sails from the wooden ship *Wasa*, likely made of linen and hemp, which sank in Stockholm harbor mud and were retrieved after 333 years (Bengtsson 1975). In contrast, in our study, cellulose filter paper buried in anoxic lake sediment showed faster disintegration than samples placed on the sediment surface. A possible explanation is that the lake sediment receives seasonal leaf litter from surrounding vegetation and therefore harbors a microbial community highly adapted to cellulose biodegradation, a hypothesis that requires further investigation.

The relatively slow biodegradation in the soil tests (fig. 2, lower right) could point to the fact that in a triphasic (gas-water-solid) medium such as soil the water availability is of great importance for microbes to be active (Kleyer 2020). A lower water content can dramatically slow down biodegradation. It can be assumed, that an optimized moisture content of the soil during the tests could have accelerated the biodegradation. This methodological shortcoming is a known complication of this test and is also addressed in the description of the respective standard ISO 17556. It is also remarkable that the often-promoted hierarchy of microbial activity toward biodegradable materials (e.g., De Wilde, 2020; Erdal and Hakkarainen, 2022), with compost and soil regarded as the most active, and freshwater and marine environments as the least active, is not supported by our experimental data.

### 4.3 Biodegradable fibers along environmental pathways

Regarding environmental fate, it is crucial not only to examine individual environmental conditions in isolation but to consider the entire system that a fiber fragment passes through, from its release to its potential arrival in the marine environment, as the final sink. This has also been acknowledged in the context of regulation such as the EU policy framework on biobased, biodegradable and compostable plastics (European Commission 2022). Most fiber fragments, including those from biodegradable fibers such as cotton or viscose are released to the air during regular use of textiles (de Falco et al., 2020) or to the water during domestic laundering (Napper and Thompson 2016; De Falco et al. 2019; Zambrano et al. 2019). If not released directly into nature, these fibers enter wastewater treatment plants (WWTPs), where a large proportion (often >90%) is retained in sewage sludge, though a measurable fraction still escapes with effluent (Ziajahromi et al. 2017; Talvitie et al. 2017).

Fibers retained in sludge may later enter soils through land application as fertilizer, where biodegradation proceeds depending on moisture, temperature, and microbial activity (Liu et al. 2021). Meanwhile, the fraction discharged with treated effluent may enter freshwater and marine systems, where fiber fragments are subject to hydrolysis, further microbial attack, and mechanical abrasion, leading to further fragmentation.

Thus, biodegradable fibers are unlikely to remain unchanged during this transport. Each compartment, WWTP, river, estuary, offers distinct biogeochemical conditions (oxygen, nutrient availability, microbial communities) that promote continued (bio)degradation. Alike ours, several studies on cellulose-based fibers demonstrated that significant biodegradation occurs in freshwater sediments and marine surface waters within weeks to months (Woodall et al. 2014; Alvarez-Zeferino et al. 2015; Royer et al. 2021, 2023; Erdal and Hakkarainen 2022). Therefore, by the time these fragments reach marine environments, they are often smaller, microbially well colonized, and structurally altered.

However, assuming that biodegradable fibers immediately enter the ocean and sink straight to the seafloor may oversimplify the process. Fiber transport in the ocean is highly dynamic, controlled by currents, density differences, turbulence, and aggregation phenomena (Kowalski et al. 2016; Karkanorachaki et al. 2021). Sinking is rarely linear: fibers can oscillate vertically, sinking and being resuspended repeatedly, over long timescales. This means it can take months to years for a distinct particle to reach deep-sea conditions (Choy et al. 2019). Given this, it is plausible that most biodegradable fiber fragments undergo substantial biodegradation before reaching the deep ocean.

Nonetheless, natural transport mechanisms, such as the sinking of marine snow, aggregates of organic detritus, mucus, plankton debris, and fine mineral particles, can accelerate vertical export (Turner 2015; Kvale et al. 2020). Micro- and nanoplastics, including fibrous particles, have been observed embedded within such agglomerates that can sink rapidly to the seafloor (Porter et al. 2018). Hence, while most biodegradable fibers may degrade during prolonged residence in the upper water column, some may become entrained in fast-sinking organic aggregates, reaching the benthic environment more quickly.

For neat cellulose fibers, our study clearly showed that they are well biodegradable in all environmental compartments, even in very cold and/or deep-sea conditions.

### 4.4 Polymer versus Product: Persistence and Environmental Relevance

A crucial aspect of assessing environmental impact is distinguishing between the neat polymer and the final product. While pure cellulose as a polymer is inherently biodegradable, the transformation into a product can substantially alter its environmental behavior. This principle is not limited to cellulosics but applies equally to biodegradable synthetic polymers and plastics derived from them.

In this context, persistence is the defining factor of environmental impact. Persistence refers not only to resistance against microbial mineralization but is also related to processes such as fragmentation and dispersal, which together shape the exposure potential and ecological consequences of fiber residues. The more stable a fiber is, the farther it can be transported, and the more likely it is to accumulate in its final sink. A polymer that is biodegradable in principle may still contribute to long-term environmental impact if modified in ways that hinder biodegradation.

Therefore, for impact assessments it is not only relevant whether a polymer class (e.g., cellulose, polyester) is inherently biodegradable but how the final product performs in real-world environments. As emphasized by Hartmann et al. 2019a and Stark 2019, and the ensuing debate (Hartmann et al. 2019b), the terminology surrounding biodegradability and persistence must be applied carefully. The distinction between polymer and product becomes decisive: a “well-biodegradable” neat polymer is of little concern, but once transformed and chemically treated, its persistence and environmental footprint can change drastically. It is therefore the final product that must be assessed for its environmental fate and impact (Stanton et al. 2021).

The SSDR as a normalized measure for fiber (bio-)degradation is used to integrate the formation and impact of solid polymers and plastic particles into a life cycle assessment (LCA), e.g. by the MariLCA initiative (www.marilca.org). The above discussion about the high abundance of treated cellulose fibers in environmental samples is also relevant to all cellulose used for fashion and technical textile applications, underscoring the need to distinguish the environmental biodegradation rates of neat cellulosic materials from those of treated cellulose fibers, rather than considering them equally. Here, using SSDRs can bring great progress to compare biodegradable and non-biodegradable materials fairly, given that the base data to calculate SSDR are reliable and originate from scientifically sound studies. The SSDR concept works well for cellulose and other surface-eroding biodegradable materials but has its limits in its application for bulk-degrading (Burkersroda et al. 2002) polymers such as poly(lactic acid), where the initial step of environmental degradation is chemical hydrolysis throughout the bulk of the object and no material loss is measurable, e.g. via thinning or surface erosion, for a long time although molecular degradation is ongoing. This fact is often ignored and can lead to wrong expectations or conclusions of experimental results (e.g., Royer et al., 2023).

With regard to environmental fate and impact, the extensive evidence presented in this study shows that distinguishing between native and regenerated cellulose is neither justified nor advisable. Efforts to reduce fiber accumulation in the environment, whether practical or regulatory, should focus on the final products entering the market, as these pose the greatest risk of release into nature regardless of their base polymers. Meaningful progress requires a life-cycle perspective that links renewable sourcing with responsible design, manufacturing, and end-of-life management. Only by doing so can we advance towards more effective solutions with less impact and avoid impeding innovation through misguided assumptions or requirements.

## Acknowledgements

We thank Daniele Fontana, and Hanna Kuhfuß for assistance during field work and access to their ponds, and Johanna Wiedling and Lena Löschel for sample preparation and field experiment maintenance. We are grateful to the administration of the Parco Nazionale Arcipelago Toscano, Portoferraio for continuous support and granting access to the protected area of the island of Pianosa (permit n.3063/19.05.2014 and following), and to dott. Emiliano Somigli and his staff for granting access to Terme San Giovanni to perform the lagoon tests. The Government of the Republic of Indonesia, Ministry of Research, Technology and Higher Education, RISTEK-DIKTI, Jakarta is gratefully thanked for granting the research permits no. 71 and 72/SIP/FRP/E5/Dit.KI/III/2017 and extensions to CL and MW, CL, and MW express their thanks to University Sam Ratulangi, Manado welcoming them as guest researchers. Thanks to Marco Segre Reinach and staff from Coral Eye Resort, Pulau Bangka, for kind support, and Gabriele ‘Lele’ Bellosta for remotely assisted sampling and maintaining field experiments during the COVID pandemic. Parts of the studies were conducted within the EU FP7 project Open-Bio and have received partial funding under the grant agreement no KBBE/FP7EN/613677 and by BASF SA, Ludwigshafen, Germany which we also thank for the allowance to use sediments and data from the positive control samples of several field trials. We are grateful to the town of Bühl administration and the county of Rastatt Landratsamt (permits Az 4.2/692.21 4.23.13 and 4.2/692.21 4.23.13) for the allowance to conduct experiments in the lakes of Weitenung and Oberbruch; to the licensees of the quarry and their workers for friendly assistance during diving operations, and to the owners of the Adamsee campground for access to their premises; to Georg and Moritz Schmälzle and staff from Biogärtnerei Schmälzle & Sohn for continuous support and assistance with field trials; to the Bergen Aquarium staff for maintenance of the tank experiment, and Gunhild Bødtker from NORCE for sampling. Thanks to Karin Schroeder CNR-ISMAR for coordinating the deep-sea mooring activities in the Mediterranean Sea. Thanks are also due to captains and crews of the Italian research vessels R/V G. DALLAPORTA and R/V GAIA BLU for technical and logistical support. Access to the moorings was provided by the EC-H2020 FP JERICO-S3 (GA No. 871153) Transnational Access Agreements N° 21/1001635 and 21/1001606. GS received financial support under the NRRP, funded by the European Union – NextGenerationEU– Project ‘MICROplastic effects on marine BEnthic Ecosystems Functioning (MICROBEEF)’ – CUP F53D23004170006 - Grant Assignment Decree No. 1015/2023 by the Italian Ministry of Ministry of University and Research. MDI mooring activities in the Arctic were supported by the ITINERIS project, funded through the EU – Next Generation PNRR program, and by the logistical assistance of the CNR Arctic Station Dirigibile Italia. Thanks to Robert Klauer, VAUDE, Germany for providing fabric samples and to Matejka Turel, Echo Instruments, Slovenia for providing different face masks. Lenzing AG is acknowledged for providing data from lab tests contracted to Normec OWS and for funding the compilation of the whole data set. Thanks to Larissa Budin, Michaela Kogler and K. Christian Schuster for background information on materials and fruitful discussions.

## SUPPLEMENTARY INFORMATION

### Part 1: Tables

**Table SI1:** Overview of materials and experimental treatments under compost conditions. The materials were exposed to various compost matrices to investigate the effects of environmental factors on their degree of disintegration or biodegradation (%). Each treatment (coded as Tn) represented a unique combination of temperature (°C), location, material, test type, matrix, vertical position, oxygen availability, and water regime. The half-life and SSDR were determined where data quality allowed. Information on sample exposure (days), fiber diameter (µm) and the applied standard is also provided.

| Treat | Code | T [°C] | Location | Material | Test Type | Matrix | Vertical position | Oxygen | Water Regime | Disintegration or Biodegradation [%] | Half-Life [d (CI)] | Exposure [d] | Fiber Ø [µm] | SSDR [µm yr<sup>-1</sup> (CI)] | Standard |
| --- | --- | --- | --- | --- | --- | --- | --- | --- | --- | --- | --- | --- | --- | --- | --- |
| Compost | T79 | 28±2 | LAB 1 | LENZING™ Lyocell A100 (crosslinked fiber), white, cryogenically milled (< 1 mm) | Lab: Biodegradability | Compost (VGF + Green Compost): Mix of mature VGF (Vegetable, Garden, Fruit waste) and green compost (garden waste). | Mixed with Matrix | Oxic | Standard | 93.1±1.7 | 18 (18.24-18.37) | 45 | 10.23 | - | ISO 14855-1 |
| Compost | T80 | 28±2 | LAB 1 | LENZING™ Lyocell Fill (siliconized fiber), white, cryogenically milled (< 1 mm) | Lab: Biodegradability | Compost (VGF + Green Compost): Mix of mature VGF (Vegetable, Garden, Fruit waste) and green compost (garden waste). | Mixed with Matrix | Oxic | Standard | 93.8±0.3 | 19 (19.35-19.60) | 45 | 23.69 | - | ISO 14855-1 |
| Compost | T81 | 28±2 | LAB 1 | LENZING™<br>Lyocell LF<br>(crosslinked<br>fiber), white,<br>cryogenically<br>milled (< 1 mm) | Lab:<br>Biodegradability | Compost<br>(VGF +<br>Green<br>Compost):<br>Mix of<br>mature VGF<br>(Vegetable,<br>Garden,<br>Fruit waste)<br>and green<br>compost<br>(garden<br>waste). | Mixed<br>with<br>Matrix | Oxic | Standard | 90.5±0.9 | 20<br>(19.96-20.15) | 45 | 10.43 | - | ISO<br>14855-1 |
| Compost | T90 | 28±2 | LAB 1 | Cellulose:<br>Native cellulose<br>powder for thin<br>layer<br>chromatography<br>(Avicel),<br>powder, white,<br>Merck Art. Nr.<br>2331 | Lab:<br>Biodegradability | Compost<br>(MSW):<br>Stabilized<br>and mature<br>compost<br>from<br>organic<br>fraction of<br>municipal<br>solid waste. | Mixed<br>with<br>Matrix | Oxic | Standard | 92.7±0.5 | 12<br>(11.98-12.28) | 45 | - | - | ISO<br>14855,<br>EN<br>13432 |
| Compost | T91 | 28±2 | LAB 1 | LENZING™<br>Viscose, white,<br>1.7/40,<br>cryogenically<br>milled (< 1 mm) | Lab:<br>Biodegradability | Compost<br>(MSW):<br>Stabilized<br>and mature<br>compost<br>from<br>organic<br>fraction of<br>municipal<br>solid waste. | Mixed<br>with<br>Matrix | Oxic | Standard | 93.1±1.9 | 21<br>(20.51-20.71) | 45 | 11.99 | - | ISO<br>14855,<br>EN<br>13432 |
| Compost | T92 | 28±2 | LAB 1 | LENZING™<br>Lyocell, white,<br>1.7/38,<br>cryogenically<br>milled (< 1 mm) | Lab:<br>Biodegradability | Compost<br>(MSW):<br>Stabilized<br>and mature<br>compost<br>from<br>organic<br>fraction of<br>municipal<br>solid waste. | Mixed<br>with<br>Matrix | Oxic | Standard | 92.6±0.9 | 33<br>(32.67-32.79) | 55 | 11.93 | - | ISO<br>14855,<br>EN<br>13432 |
| Compost | TLy-1/3 | 28±2 | LAB 1 | LENZING™<br>Lyocell, white,<br>1.7/38,<br>nonwoven<br>sheet, thickness:<br>348 µm, 62<br>g/m2 | Lab:<br>Disintegration | Compost:<br>80/20 mix<br>of < 10 mm<br>mature<br>compost<br>(aged 17<br>weeks) &<br>fresh milled<br>VGF<br>(Vegetable,<br>Garden,<br>Fruit waste) | Mixed<br>with<br>Matrix | Oxic | Moistened<br>if needed | 100 | - | 112 | 11.93 | 11.39 | ISO<br>20200 |
| Compost | Vi-1/3 | 28±2 | LAB 1 | LENZING™<br>Viscose, white,<br>1.7/40,<br>nonwoven<br>sheet, thickness:<br>273 µm, 55<br>g/m2 | Lab:<br>Disintegration | Compost:<br>80/20 mix<br>of < 10 mm<br>mature<br>compost<br>(aged 17<br>weeks) &<br>fresh milled<br>VGF<br>(Vegetable,<br>Garden,<br>Fruit waste) | Mixed<br>with<br>Matrix | Oxic | Moistened<br>if needed | 100 | - | 42 | 11.99 | 30.53 | ISO<br>20200 |
| Compost | ViLy-1/3 | 28±2 | LAB 1 | LENZING™<br>Viscose/Lyocell<br>(50%/50%),<br>white, Viscose:<br>1.7/40, Lyocell:<br>1.7/38,<br>nonwoven<br>sheet, thickness:<br>289 µm, 57<br>g/m2 | Lab:<br>Disintegration | Compost:<br>80/20 mix<br>of < 10 mm<br>mature<br>compost<br>(aged 17<br>weeks) &<br>fresh milled<br>VGF<br>(Vegetable,<br>Garden,<br>Fruit waste) | Mixed<br>with<br>Matrix | Oxic | Moistened<br>if needed | 100 | - | 84 | 11.96 | 15.22 | ISO<br>20200 |

**Table SI2:** Overview of materials and experimental treatments under soil conditions. The materials were exposed to various soil environments to investigate the effects of environmental factors on their degree of disintegration or biodegradation (%). Each treatment (coded as Tn) represented a unique combination of temperature (°C), location, material, test type, matrix, vertical position, water availability or type of irrigation, and nutrient addition. The half-life and SSDR were determined where data quality allowed. Information on sample exposure (days), fiber diameter (µm) and the applied standard is also provided.

| Treat | Code | T [°C] | Location | Material | Test Type | Matrix | Vertical position | Water availability or type of irrigation | Nutrient addition | Disintegration or Biodegradation [%] | Half-Life [d (CI)] | Exposure [d] | Fiber Ø [µm] | SSDR [µm yr<sup>-1</sup> (CI)] | Standard |
| --- | --- | --- | --- | --- | --- | --- | --- | --- | --- | --- | --- | --- | --- | --- | --- |
| Soil | T21 | 24 (18.6-30.8) | LAB 2 | Cellulose filter paper (MN 640 d, 85 g/m<sup>2</sup>, 170 µm thickness, Macherey-Nagel, Germany) | Tank: Disintegration | Forest Soil (F) | Surface (0 cm) | None | None |  | 616 (458.42-1098.16) | 621 | 20 | 1.73 (0.96–2.33) | ISO 23832:2021, adapted to soil |
| Soil | T22 | 24 (18.6-30.8) | LAB 2 | Cellulose filter paper (MN 640 d, 85 g/m<sup>2</sup>, 170 µm thickness, Macherey-Nagel, Germany) | Tank: Disintegration | Garden Soil (G) | Surface (0 cm) | None | None | 11% after 621 d | - | 621 | 20 | - | ISO 23832:2021, adapted to soil |
| Soil | T23 | 24 (18.6-30.8) | LAB 2 | Cellulose filter paper (MN 640 d, 85 g/m<sup>2</sup>, 170 µm thickness, Macherey-Nagel, Germany) | Tank: Disintegration | Mixed Soil (F+G) | Surface (0 cm) | None | None | 3% after 621 d | - | 621 | 20 | - | ISO 23832:2021, adapted to soil |
| Soil | T24 | 24 (18.6-30.5) | LAB 2 | Cellulose filter paper (MN 640 d, 85 g/m<sup>2</sup>, 170 µm thickness, Macherey-Nagel, Germany) | Tank: Disintegration | Forest Soil (F) | Buried (3 cm) | None | None |  | 73 (65.22-80.96) | 529 | 20 | 14.59 (13.2–16.39) | ISO 23832:2021, adapted to soil |
| Soil | T25 | 24<br>(18.6-30.5) | LAB 2 | Cellulose filter paper (MN 640 d, 85 g/m<sup>2</sup>, 170 µm thickness, Macherey-Nagel, Germany) | Tank: Disintegration | Garden Soil (G) | Buried (3 cm) | None | None |  | 102<br>(85.11-117.88) | 207 | 20 | 10.44<br>(9.07–12.56) | ISO 23832:2021, adapted to soil |
| Soil | T26 | 24<br>(18.6-30.5) | LAB 2 | Cellulose filter paper (MN 640 d, 85 g/m<sup>2</sup>, 170 µm thickness, Macherey-Nagel, Germany) | Tank: Disintegration | Mixed Soil (F+G) | Buried (3 cm) | None | None |  | 61<br>(52.78-68.06) | 145 | 20 | 17.64<br>(15.71–20.26) | ISO 23832:2021, adapted to soil |
| Soil | T27 | 24<br>(18.6-29.8) | LAB 2 | Cellulose filter paper (MN 640 d, 85 g/m<sup>2</sup>, 170 µm thickness, Macherey-Nagel, Germany) | Tank: Disintegration | Agricultural Soil | Surface (0 cm) | None | None | 9% after 387 d | - | 387 | 20 | - | ISO 23832:2021, adapted to soil |
| Soil | T28a | 24<br>(13.6-26.3) | LAB 2 | Cellulose filter paper (MN 640 d, 85 g/m<sup>2</sup>, 170 µm thickness, Macherey-Nagel, Germany) | Tank: Disintegration | Agricultural Soil | Buried (3 cm) | None | None |  | 35 (0-56.55) | 153 | 20 | 30.04<br>(-42.57–146.13) | ISO 23832:2021, adapted to soil |
| Soil | T28b | 24<br>(13.8-26) | LAB 2 | Cellulose filter paper (MN 640 d, 85 g/m<sup>2</sup>, 170 µm thickness, Macherey-Nagel, Germany) | Tank: Disintegration | Agricultural Soil | Buried (3 cm) | None | None | 0% after 197 d | - | 197 | 20 | - | ISO 23832:2021, adapted to soil |
| Soil | T29 | 24<br>(13.8-26) | LAB 2 | Cellulose filter paper (MN 640 d, 85 g/m<sup>2</sup>, 170 µm thickness, Macherey-Nagel, Germany) | Tank: Disintegration | Agricultural Soil | Surface (0 cm) | Irrigation, Medium | None |  | 180<br>(173.86-186.17) | 197 | 20 | 5.95<br>(5.74–6.15) | ISO 23832:2021, adapted to soil |
| Soil | T30 | 24<br>(13.8-26) | LAB 2 | Cellulose filter paper (MN 640 d, 85 g/m<sup>2</sup>, 170 µm thickness, Macherey-Nagel, Germany) | Tank: Disintegration | Agricultural Soil | Buried (3 cm) | Irrigation, Medium | None |  | 98<br>(95.1-101.35) | 197 | 20 | 10.88<br>(10.55–11.24) | ISO 23832:2021, adapted to soil |
| Soil | T31 | 23<br>(13.9-26) | LAB 2 | Cellulose filter paper (MN 640 d, 85 g/m<sup>2</sup>, 170 µm thickness, Macherey-Nagel, Germany) | Tank: Disintegration | Agricultural Soil | Surface (0 cm) | Irrigation, Medium | Compost Tea |  | 53<br>(47.6-57.68) | 197 | 20 | 20.33<br>(18.53–22.46) | ISO 23832:2021, adapted to soil |
| Soil | T32 | 23<br>(13.9-26) | LAB 2 | Cellulose filter paper (MN 640 d, 85 g/m<sup>2</sup>, 170 µm thickness, Macherey-Nagel, Germany) | Tank: Disintegration | Agricultural Soil | Buried (3 cm) | Irrigation, Medium | Compost Tea |  | 41<br>(29.49-51.6) | 197 | 20 | 26.16<br>(20.72–36.25) | ISO 23832:2021, adapted to soil |
| Soil | T33 | 23<br>(13.8-25) | LAB 2 | Cellulose filter paper (MN 640 d, 85 g/m<sup>2</sup>, 170 µm thickness, Macherey-Nagel, Germany) | Tank: Disintegration | Agricultural Soil | Surface (0 cm) | Irrigation, Wet | None |  | 61 (55-66.45) | 197 | 20 | 17.62<br>(16.09–19.44) | ISO 23832:2021, adapted to soil |
| Soil | T34 | 23<br>(13.8-25) | LAB 2 | Cellulose filter paper (MN 640 d, 85 g/m<sup>2</sup>, 170 µm thickness, Macherey-Nagel, Germany) | Tank: Disintegration | Agricultural Soil | Buried (3 cm) | Irrigation, Wet | None |  | 57<br>(50.64-62.83) | 197 | 20 | 18.86<br>(17.01–21.11) | ISO 23832:2021, adapted to soil |
| Soil | T35 | 16 (-6.1-47.7) | Agriculture | Cellulose filter paper (MN 640 d, 85 g/m<sup>2</sup>, 170 µm thickness, Macherey-Nagel, Germany) | Field (Greenhouse): Disintegration | Agricultural Soil | Surface (0 cm) | Drip-Line Irrigation | None | 0% after 364 d | - | 364 | 20 | - | EN ISO 22766:2020, adapted to soil |
| Soil | T36 | 14<br>(2.6-28.3) | Agriculture | Cellulose filter paper (MN 640 d, 85 g/m<sup>2</sup>, 170 µm thickness, Macherey-Nagel, Germany) | Field: Disintegration | Agricultural Soil | Buried (5 cm) | Rain, Natural precipitation | None |  | 37<br>(29.76-43.51) | 74 | 20 | 29.05<br>(24.57–35.92) | EN ISO 22766:2020, adapted to soil |
| Soil | T37 | 22<br>(2.9-47.7) | Agriculture | Cellulose filter paper (MN 640 d, 85 g/m<sup>2</sup>, 170 µm thickness, Macherey-Nagel, Germany) | Field (Greenhouse): Disintegration | Agricultural Soil | Buried (3 cm) | Drip-Line Irrigation | None |  | 81<br>(73.35-89.56) | 155 | 20 | 13.15<br>(11.94–14.57) | EN ISO 22766:2020, adapted to soil |
| Soil | T38 | 15 (-2.2-31.6) | Agriculture | Cellulose filter paper (MN 640 d, 85 g/m<sup>2</sup>, 170 µm thickness, Macherey-Nagel, Germany) | Field (Greenhouse): Disintegration | Agricultural Soil | Surface, Silage cover (10 cm) | Drip-Line Irrigation | None |  | 245<br>(180.78-353.32) | 364 | 20 | 4.37<br>(3.02–5.91) | EN ISO 22766:2020, adapted to soil |
| Soil | T39 | 20 (-2.1-54.1) | Agriculture | Cellulose filter paper (MN 640 d, 85 g/m<sup>2</sup>, 170 µm thickness, Macherey-Nagel, Germany) | Field: Disintegration | Agricultural Soil | Surface (0 cm) | Rain, Natural precipitation | None |  | 132<br>(118.61-146.56) | 286 | 20 | 8.07<br>(7.29–9.01) | EN ISO 22766:2020, adapted to soil |
| Soil | T40 | 14<br>(2.6-28.3) | Agriculture | Cellulose filter paper (MN 640 d, 85 g/m<sup>2</sup>, 170 µm thickness, Macherey-Nagel, Germany) | Field: Disintegration | Agricultural Soil | Buried (5 cm) | Rain, Natural precipitation | None |  | 23<br>(17.77-28.47) | 74 | 20 | 45.69<br>(37.55–60.16) | EN ISO 22766:2020, adapted to soil |
| Soil | T41 | 13<br>(5.1-31.6) | Agriculture | Cellulose filter paper (MN 640 d, 85 g/m<sup>2</sup>, 170 µm thickness, Macherey-Nagel, Germany) | Field: Disintegration | Agricultural Soil | Buried (15 cm) | Rain, Natural precipitation | None |  | 23<br>(16.52-28.74) | 74 | 20 | 46.47<br>(37.2–64.72) | EN ISO 22766:2020, adapted to soil |
| Soil | T42 | 11<br>(2.6-21.9) | Agriculture | Cellulose filter paper (MN 640 d, 85 g/m<sup>2</sup>, 170 µm thickness, Macherey-Nagel, Germany) | Field: Disintegration | Agricultural Soil | Buried (5 cm) | Irrigation | None |  | 25<br>(24.66-25.53) | 40 | 20 | 42.59<br>(41.88–43.35) | EN ISO 22766:2020, adapted to soil |
| Soil | T43 | 14<br>(2.6-28.3) | Agriculture | Cellulose filter paper (MN 640 d, 85 g/m<sup>2</sup>, 170 µm thickness, Macherey-Nagel, Germany) | Field: Disintegration | Agricultural Soil | Buried (5 cm) | Irrigation | Compost Tea |  | 37<br>(23.65-48.77) | 74 | 20 | 29.26<br>(21.9–45.11) | EN ISO 22766:2020, adapted to soil |
| Soil | T86 | 25±2 | LAB 1 | Cellulose: Native cellulose powder for thin layer chromatography (Avicel), powder, white, Merck Art. Nr. 2331 | Lab: Biodegradability | Lab Mix: 70% industrial quartz sand, 10% kaolinite clay, 16% natural soil and 4% mature compost | Mixed with Matrix | Humidified to compensate evaporation | 4% mature compost | 73.8±0.8 | 20<br>(19.27-20.24) | 122 | - | - | ISO 17556 |
| Soil | T87 | 25±2 | LAB 1 | LENZING™ Lyocell A100 (crosslinked fiber), white, cryogenically milled (< 1 mm) | Lab: Biodegradability | Lab Mix: 70% industrial quartz sand, 10% kaolinite clay, 16% natural soil and 4% mature compost | Mixed with Matrix | Humidified to compensate evaporation | 4% mature compost | 68.8±3.4 | 53<br>(52.39-53.35) | 122 | 10.23 | - | ISO 17556 |
| Soil | T88 | 25±2 | LAB 1 | LENZING™ Lyocell Fill (siliconized fiber), white, cryogenically milled (< 1 mm) | Lab: Biodegradability | Lab Mix: 70% industrial quartz sand, 10% kaolinite clay, 16% natural soil and 4% mature compost | Mixed with Matrix | Humidified to compensate evaporation | 4% mature compost | 66.5±0.3 | 67<br>(67.21-67.54) | 122 | 23.69 | - | ISO 17556 |
| Soil | T89 | 25±2 | LAB 1 | LENZING™<br>Lyocell LF<br>(crosslinked<br>fiber), white,<br>cryogenically<br>milled (< 1 mm) | Lab:<br>Biodegradability | Lab Mix:<br>70%<br>industrial<br>quartz<br>sand, 10%<br>kaolinite<br>clay, 16%<br>natural soil<br>and 4%<br>mature<br>compost | Mixed<br>with<br>Matrix | Humidified<br>to<br>compensate<br>evaporation | 4%<br>mature<br>compost | 69.2±2.1 | 60<br>(59.23-60.45) | 122 | 10.43 | - | ISO 17556 |

**Table SI3:** Overview of materials and experimentalt under freshwater conditions. The materials were exposed to various freshwater environments to investigate the effects of environmental factors on their degree of disintegration or biodegradation (%). Each treatment (coded as Tn) represented a unique combination of temperature (°C), location, material, test type, matrix, vertical position, and oxygen availability. The half-life and SSDR were determined where data quality allowed. Information on sample exposure (days), fiber diameter (µm) and the applied standard is also provided.

| Treat | Code | T [°C] | Location | Material | Test Type | Matrix | Vertical position | Oxygen | Sample Type | Disintegration or Biodegradation [%] | Half-Life [d (CI)] | Exposure [d] | Fiber Ø [µm] | SSDR [µm yr<sup>-1</sup> (CI)] | Standard |
| --- | --- | --- | --- | --- | --- | --- | --- | --- | --- | --- | --- | --- | --- | --- | --- |
| Freshwater | T1 | 15 (4.8-25.8) | Adamsee | Cellulose filter paper (MN 640 d, 85 g/m<sup>2</sup>, 170 µm thickness, Macherey-Nagel, Germany) | Field: Disintegration | Lake 2 | Buried | Anoxic | Check |  | 51 (33.78-79.4) | 62 | 20 | 21.01 (13.34-31.41) | EN ISO 22766:202, adapted to freshwater |
| Freshwater | T2 | 15 (4.8-25.8) | Adamsee | Cellulose filter paper (MN 640 d, 85 g/m<sup>2</sup>, 170 µm thickness, Macherey-Nagel, Germany) | Field: Disintegration | Lake 2 | Benthic | Oxic | Check |  | 240 (210.98-271.12) | 425 | 20 | 4.45 (3.94-5.07) | EN ISO 22766:202, adapted to freshwater |
| Freshwater | T3 | 15 (4.8-25.8) | Adamsee | Cellulose filter paper (MN 640 d, 85 g/m<sup>2</sup>, 170 µm thickness, Macherey-Nagel, Germany) | Field: Disintegration | Lake 2 | Buried | Anoxic | True |  | 59 (56.83-60.94) | 120 | 20 | 18.16 (17.54-18.81) | EN ISO 22766:202, adapted to freshwater |
| Freshwater | T4 | 15 (4.8-25.8) | Adamsee | Cellulose filter paper (MN 640 d, 85 g/m<sup>2</sup>, 170 µm thickness, Macherey-Nagel, Germany) | Field: Disintegration | Lake 2 | Benthic | Oxic | True |  | 130 (111.98-148.01) | 293 | 20 | 8.25 (7.22-9.55) | EN ISO 22766:202, adapted to freshwater |
| Freshwater | T7 | 18 | LAB 2 | Cellulose filter paper (MN 640 d, 85 g/m<sup>2</sup>, 170 µm thickness, Macherey-Nagel, Germany) | Tank: Disintegration | Lake 2 | Benthic | Oxic | Check |  | 219<br>(209.22-228.9) | 315 | 20 | 4.89<br>(4.67–5.11) | ISO 23832:2021, adapted to freshwater |
| Freshwater | T8 | 26 | LAB 2 | Cellulose filter paper (MN 640 d, 85 g/m<sup>2</sup>, 170 µm thickness, Macherey-Nagel, Germany) | Tank: Disintegration | Lake 2 | Benthic | Oxic | Check |  | 80<br>(73.41-85.81) | 147 | 20 | 13.44<br>(12.46–14.56) | ISO 23832:2021, adapted to freshwater |
| Freshwater | T9 | 18 | LAB 2 | Cellulose filter paper (MN 640 d, 85 g/m<sup>2</sup>, 170 µm thickness, Macherey-Nagel, Germany) | Tank: Disintegration | Creek | Benthic | Oxic | Check |  | 29<br>(23.42-33.31) | 67 | 20 | 36.96<br>(32.09–45.66) | ISO 23832:2021, adapted to freshwater |
| Freshwater | T10 | 26 | LAB 2 | Cellulose filter paper (MN 640 d, 85 g/m<sup>2</sup>, 170 µm thickness, Macherey-Nagel, Germany) | Tank: Disintegration | Creek | Benthic | Oxic | Check |  | 25<br>(23.54-26.13) | 42 | 20 | 43.06<br>(40.91–45.41) | ISO 23832:2021, adapted to freshwater |
| Freshwater | T46 | 28 | LAB 2 | Cellulose filter paper (MN 640 d, 85 g/m<sup>2</sup>, 170 µm thickness, Macherey-Nagel, Germany) | Tank: Disintegration | Lake 1 | Benthic | Oxic | Check |  | 54<br>(38.82-71.81) | 138 | 20 | 19.91<br>(14.89–27.54) | ISO 23832:2021, adapted to freshwater |
| Freshwater | T51 | 26 | LAB 2 | Cellulose filter paper (MN 640 d, 85 g/m<sup>2</sup>, 170 µm thickness, Macherey-Nagel, Germany) | Tank: Disintegration | Lake 1 | Buried | Anoxic | Check |  | 41<br>(38.2-43.56) | 90 | 20 | 26.13<br>(24.54–27.99) | ISO 23832:2021, adapted to freshwater |
| Freshwater | T52 | 26 | LAB 2 | Cellulose filter paper (MN 640 d, 85 g/m<sup>2</sup>, 170 µm thickness, Macherey- | Tank: Disintegration | Creek | Buried | Anoxic | Check |  | 34<br>(28.13-39.73) | 119 | 20 | 31.14<br>(26.91–38) | ISO 23832:2021, adapted to freshwater |
|  |  |  |  | Nagel,<br>Germany) |  |  |  |  |  |  |  |  |  |  |  |
| Freshwater | T65 | 12<br>(2-21.6) | LAB 2 | Cellulose filter<br>paper (MN 640<br>d, 85 g/m<sup>2</sup>, 170<br>µm thickness,<br>Macherey-Nagel,<br>Germany) | Field:<br>Disintegration | Lake 3 | Benthic | Oxic | True | 2% after 70 d<br>100% after 115<br>d | - | 115 | 20 | - | EN ISO<br>22766:202,<br>adapted to<br>freshwater |
| Freshwater | T66 | 12<br>(7-26.3) | LAB 2 | Cellulose filter<br>paper (MN 640<br>d, 85 g/m<sup>2</sup>, 170<br>µm thickness,<br>Macherey-Nagel,<br>Germany) | Field:<br>Disintegration | Lake 1 | Benthic | Oxic | True | 96% after 245<br>d | - | 245 | 20 | - | EN ISO<br>22766:202,<br>adapted to<br>freshwater |
| Freshwater | T67 | 15<br>(3.9-32.9) | LAB 2 | Cellulose filter<br>paper (MN 640<br>d, 85 g/m<sup>2</sup>, 170<br>µm thickness,<br>Macherey-Nagel,<br>Germany) | Field:<br>Disintegration | Lake 4 | Benthic | Oxic | True | 100% after 328<br>d | - | 328 | 20 | - | EN ISO<br>22766:202,<br>adapted to<br>freshwater |
| Freshwater | T73 | 21±1 | LAB 1 | Cellulose:<br>Native cellulose<br>powder for thin<br>layer<br>chromatography<br>(Avicel),<br>powder, white,<br>Merck Art. Nr.<br>2331 | Lab:<br>Biodegradability | Freshwat<br>er +<br>Sludge<br>Inoculum<br>(F+SI) | Pelagic | Oxic | True | 78.9±2.0 | 9 (8.64-8.87) | 28 | 20 | - | ISO 14851 |
| Freshwater | T74 | 21±1 | LAB 1 | LENZING™<br>Modal, white,<br>1.7/39, tow, cut<br>into small pieces<br>with scissors | Lab:<br>Biodegradability | Freshwat<br>er +<br>Sludge<br>Inoculum<br>(F+SI) | Pelagic | Oxic | True | 74.8 ± 0.6 | 11<br>(10.39-10.70) | 28 | 11.96 | 60.58<br>(59.7–<br>61.49) | ISO 14851 |
| Freshwater | T75 | 21±1 | LAB 1 | LENZING™<br>Viscose, white,<br>1.7/40, tow, cut<br>into small pieces<br>with scissors | Lab:<br>Biodegradability | Freshwat<br>er +<br>Sludge<br>Inoculum<br>(F+SI) | Pelagic | Oxic | True | 72.7 ± 0.6 | 14<br>(14.22-14.34) | 28 | 11.99 | 44.87<br>(44.69–<br>45.06) | ISO 14851 |
| Freshwater | T76 | 21±1 | LAB 1 | LENZING™<br>Lyocell, white,<br>1.7/38, tow, cut<br>into small pieces<br>with scissors | Lab:<br>Biodegradability | Freshwat<br>er +<br>Sludge<br>Inoculum<br>(F+SI) | Pelagic | Oxic | True | 83.3 ± 3.1 | 17<br>(16.74-16.85) | 42 | 11.93 | 37.97<br>(37.85–<br>38.09) | ISO 14851 |
| Freshwater | T77 | 21±1 | LAB 1 | Cellulose:<br>Native cellulose<br>powder for thin<br>layer<br>chromatography<br>(Avicel),<br>powder, white,<br>Merck Art. Nr.<br>2331 | Lab:<br>Biodegradability | Freshwat<br>er +<br>Sludge<br>Inoculum<br>(F+SI) | Pelagic | Oxic | True | 85 | 9 (9.14-9.30) | 42 | - | - | ISO 14851 |
| Freshwater | T82 | 21±1 | LAB 1 | Cellulose:<br>Native cellulose<br>powder for thin<br>layer<br>chromatography<br>(Avicel),<br>powder, white,<br>Merck Art. Nr.<br>2331 | Lab:<br>Biodegradability | Freshwat<br>er +<br>Sludge<br>Inoculum<br>(F+SI) | Pelagic | Oxic | True | 87 | 15<br>(15.09-15.66) | 42 | - | - | ISO 14851 |
| Freshwater | T83 | 21±1 | LAB 1 | LENZING™<br>Lyocell A100<br>(crosslinked<br>fiber), white,<br>cryogenically<br>milled (< 1 mm) | Lab:<br>Biodegradability | Freshwat<br>er +<br>Sludge<br>Inoculum<br>(F+SI) | Pelagic | Oxic | True | 79.6 ± 6.3 | 14<br>(13.42-14.01) | 42 | 10.23 | - | ISO 14851 |
| Freshwater | T84 | 21±1 | LAB 1 | LENZING™<br>Lyocell Fill<br>(siliconized<br>fiber), white,<br>cryogenically<br>milled (< 1 mm),<br>6.7/60 | Lab:<br>Biodegradability | Freshwat<br>er +<br>Sludge<br>Inoculum<br>(F+SI) | Pelagic | Oxic | True | 80.8 ± 1.9 | 13<br>(13.07-13.49) | 42 | 23.69 | - | ISO 14851 |
| Freshwater | T85 | 21±1 | LAB 1 | LENZING™<br>Lyocell LF<br>(crosslinked<br>fiber), white,<br>cryogenically<br>milled (< 1 mm),<br>1.3/38 | Lab:<br>Biodegradability | Freshwat<br>er +<br>Sludge<br>Inoculum<br>(F+SI) | Pelagic | Oxic | True | 83.4 ± 1.0 | 13<br>(12.59-13.09) | 42 | 10.43 | - | ISO 14851 |
| Freshwater | T102 | 21±1 | LAB 1 | Cellulose:<br>Native cellulose powder for thin layer chromatography (Avicel), powder, white, Merck Art. Nr. 2331 | Lab:<br>Biodegradability | Freshwater + Sludge Inoculum (F+SI) | Pelagic | Oxic | True | 76 | - | 42 | - | - | ISO 14851 |
| Freshwater | T103 | 21±1 | LAB 1 | LENZING™<br>Lyocell, white, 1.7/38, cryogenically milled (< 1mm) | Lab:<br>Biodegradability | Freshwater + Sludge Inoculum (F+SI) | Pelagic | Oxic | True | 78 | 22 (21.78-21.97) | 42 | 11.93 | 29.15 (29.02–29.28) | ISO 14851 |
| Freshwater | T104 | 21±1 | LAB 1 | Svilosa Kraft<br>Paper pulp, grated, reduced in size with scissors | Lab:<br>Biodegradability | Freshwater + Sludge Inoculum (F+SI) | Pelagic | Oxic | True | 83 | 19 (18.5-19.29) | 42 | - | - | ISO 14851 |
| Freshwater | T105 | 21±1 | LAB 1 | LENZING™<br>Viscose, white, 1,7/40, cryogenically milled (< 1 mm) | Lab:<br>Biodegradability | Freshwater + Sludge Inoculum (F+SI) | Pelagic | Oxic | True | 85 | 19 (19.15-19.39) | 42 | 11.99 | - | ISO 14851 |
| Freshwater | T106 | 21±1 | LAB 1 | Canfor ECF 90, grated, reduced in size with scissors | Lab:<br>Biodegradability | Freshwater + Sludge Inoculum (F+SI) | Pelagic | Oxic | True | 88 | 21 (19.41-20.74) | 42 | - | - | ISO 14851 |
| Freshwater | T107 | 21±1 | LAB 1 | LENZING™<br>Modal, white, 1,7/39, cryogenically milled (< 1 mm) | Lab:<br>Biodegradability | Freshwater + Sludge Inoculum (F+SI) | Pelagic | Oxic | True | 83 | - | 42 | 11.96 | - | ISO 14851 |
| Freshwater | T108 | 21±1 | LAB 1 | Organic cotton, white, 2.2/25, cryogenically milled (< 1 mm) | Lab:<br>Biodegradability | Freshwater + Sludge Inoculum (F+SI) | Pelagic | Oxic | True | 75 | 27 (23.51-27.17) | 42 | 15.68 | - | ISO 14851 |
| Freshwater | T109 | 21±1 | LAB 1 | Abaca pulp (PSPI-100), grated, reduced in size with scissors | Lab: Biodegradability | Freshwater + Sludge Inoculum (F+SI) | Pelagic | Oxic | True | 86 | 16 (16.10-16.52) | 42 | - | - | ISO 14851 |
| Freshwater | T113 | 28±1 | LAB 2 | Cellulose filter paper, Macherey-Nagel, MN 640 d, sheet thickness 135 µm, fiber diameter nominally 20 µm, ash free, white, cut into small pieces | Lab: Biodegradability | Freshwater + Lake 1 Sediment | Benthic | Oxic | True | 79 | - | 50 | 20 | 46.35 (44.77–47.97) | ISO 18830, adapted to freshwater, modified (stirred), O2 consumption |

**Table SI4:**
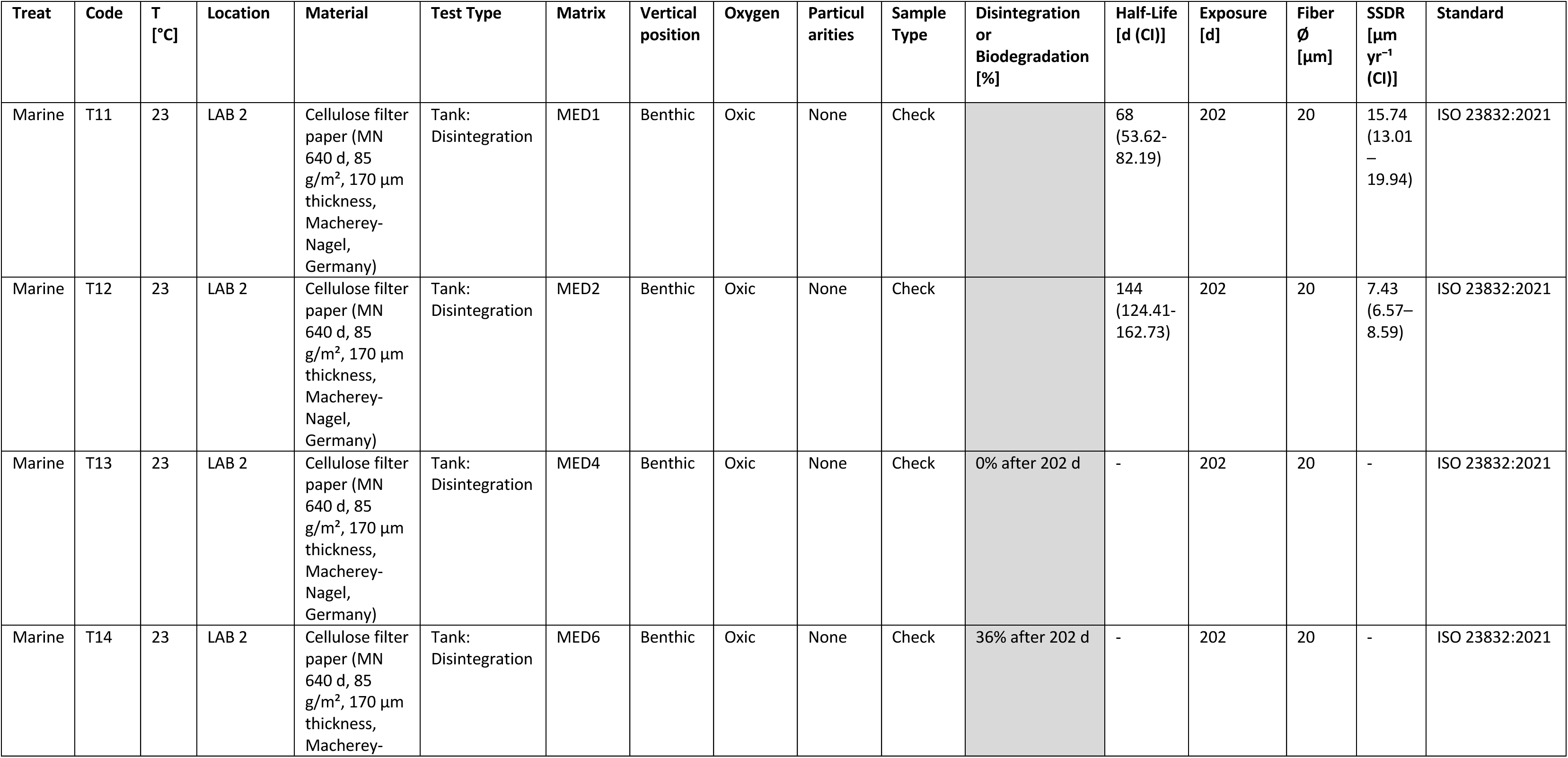
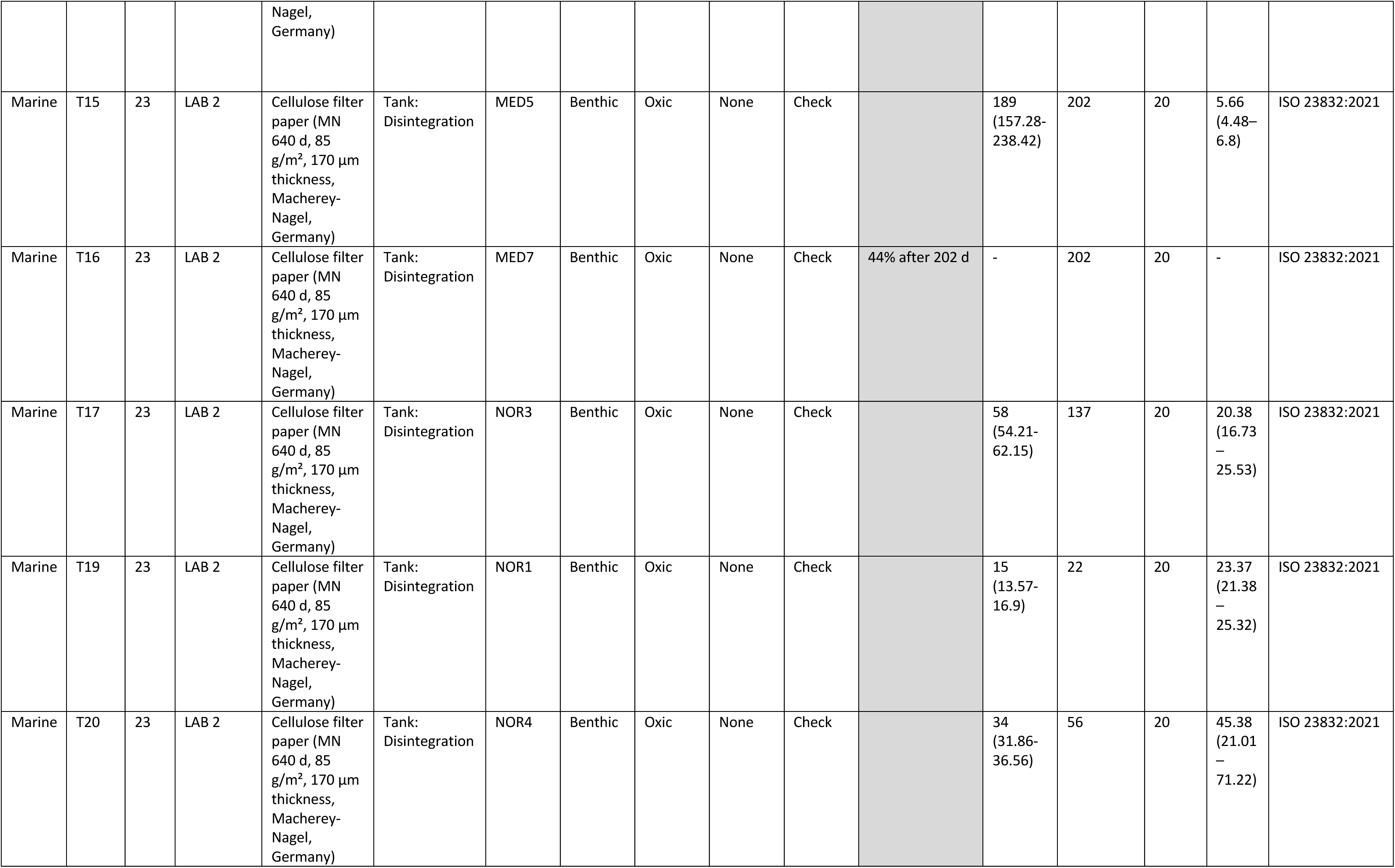
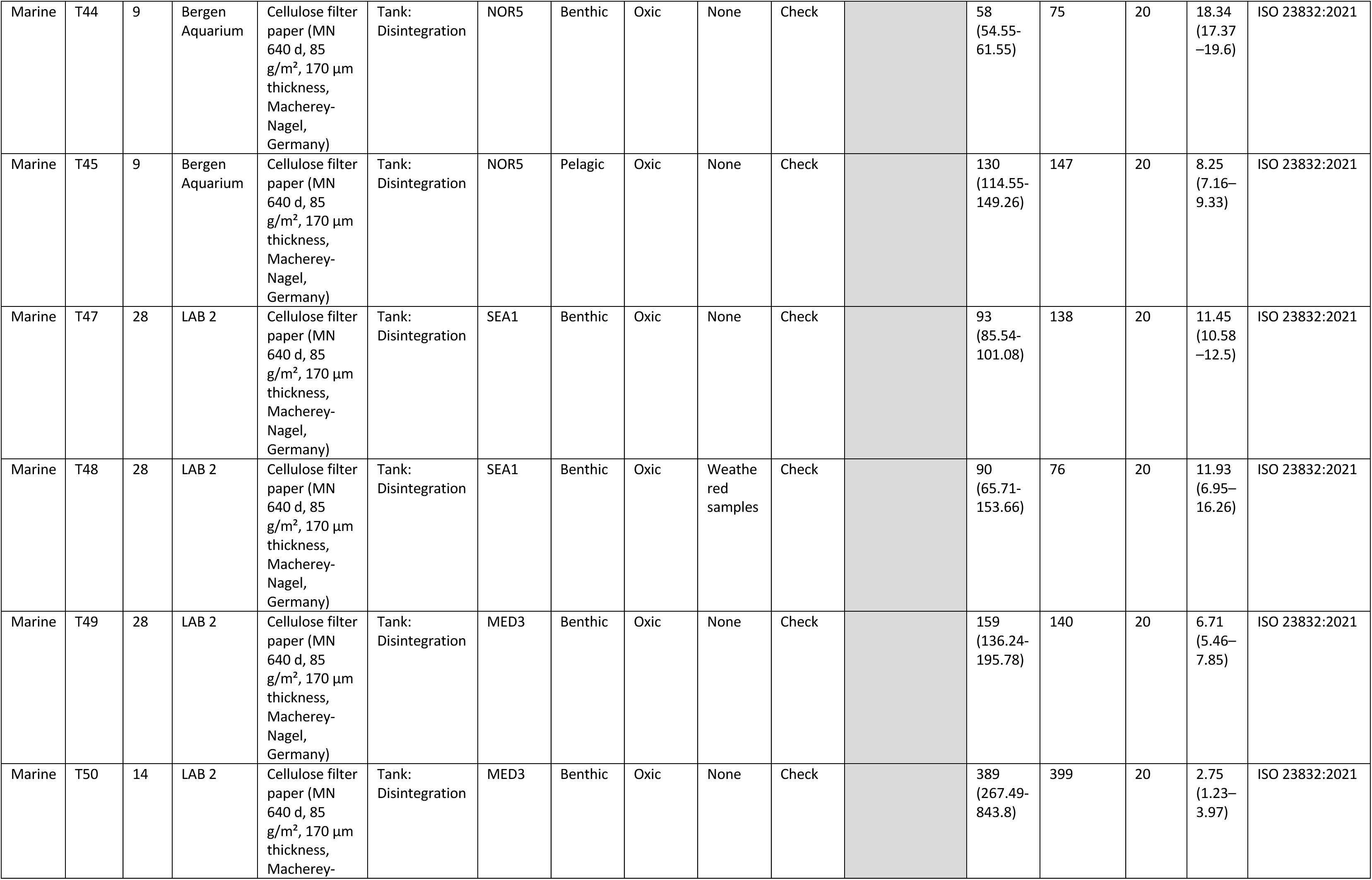
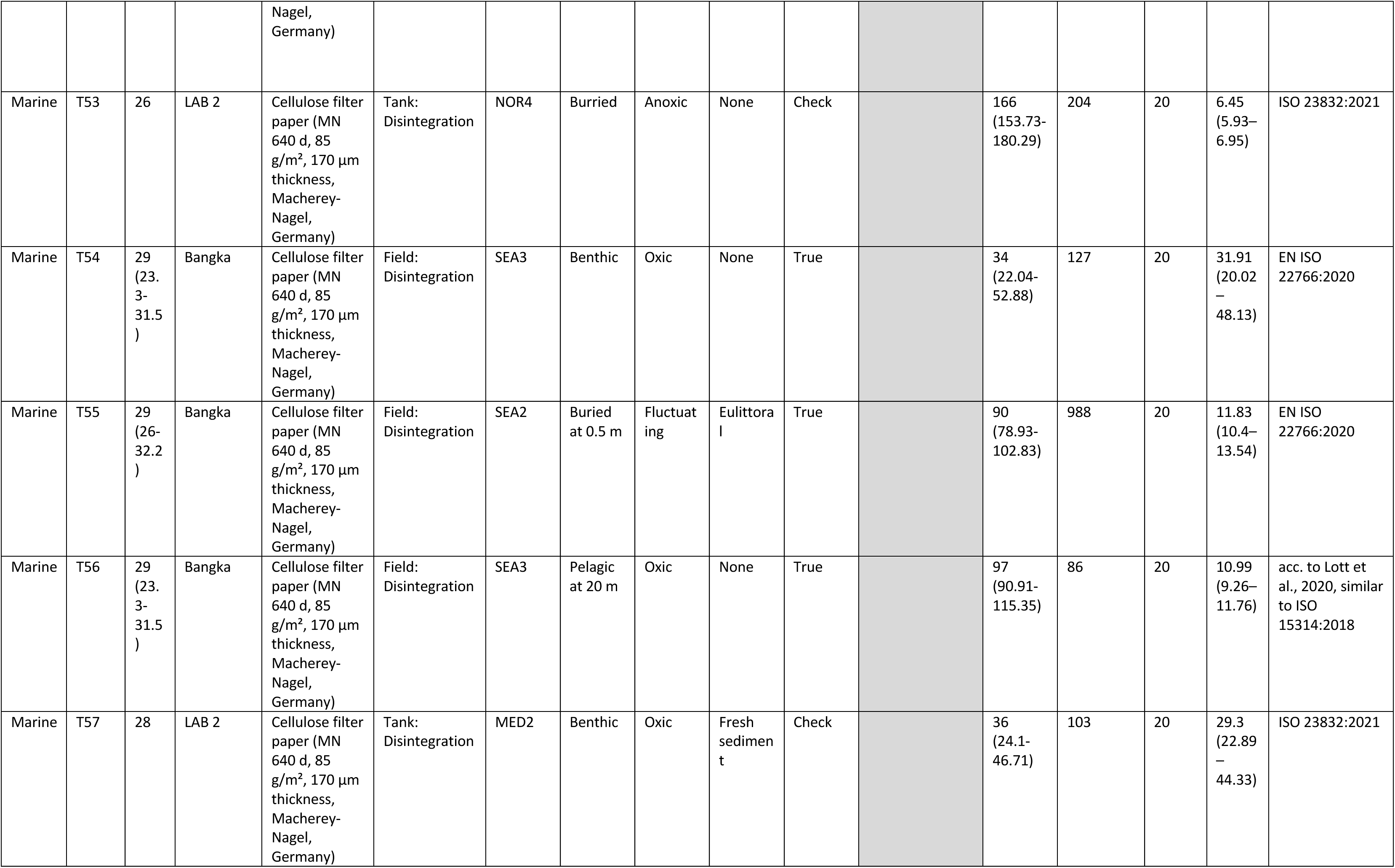
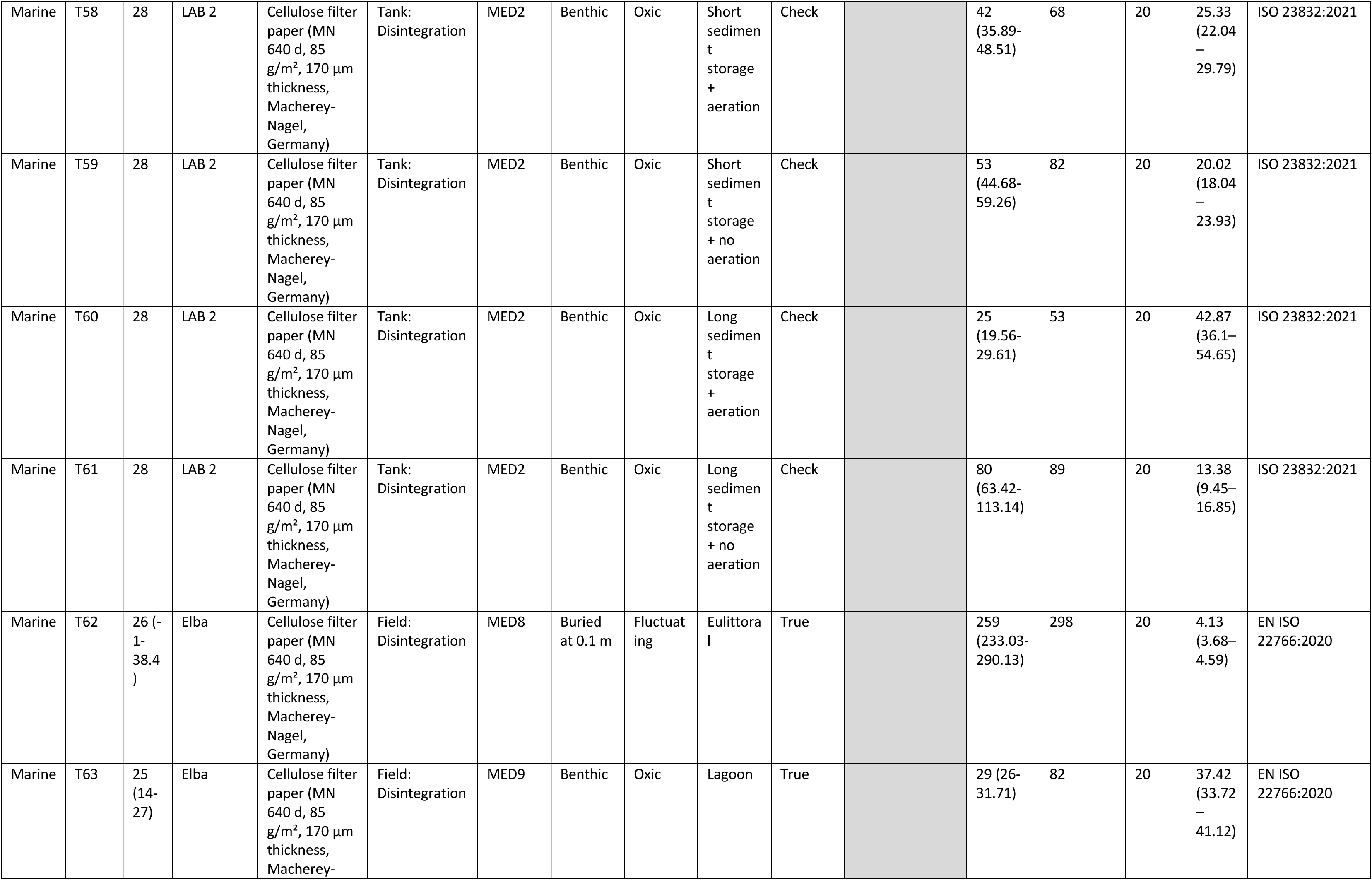
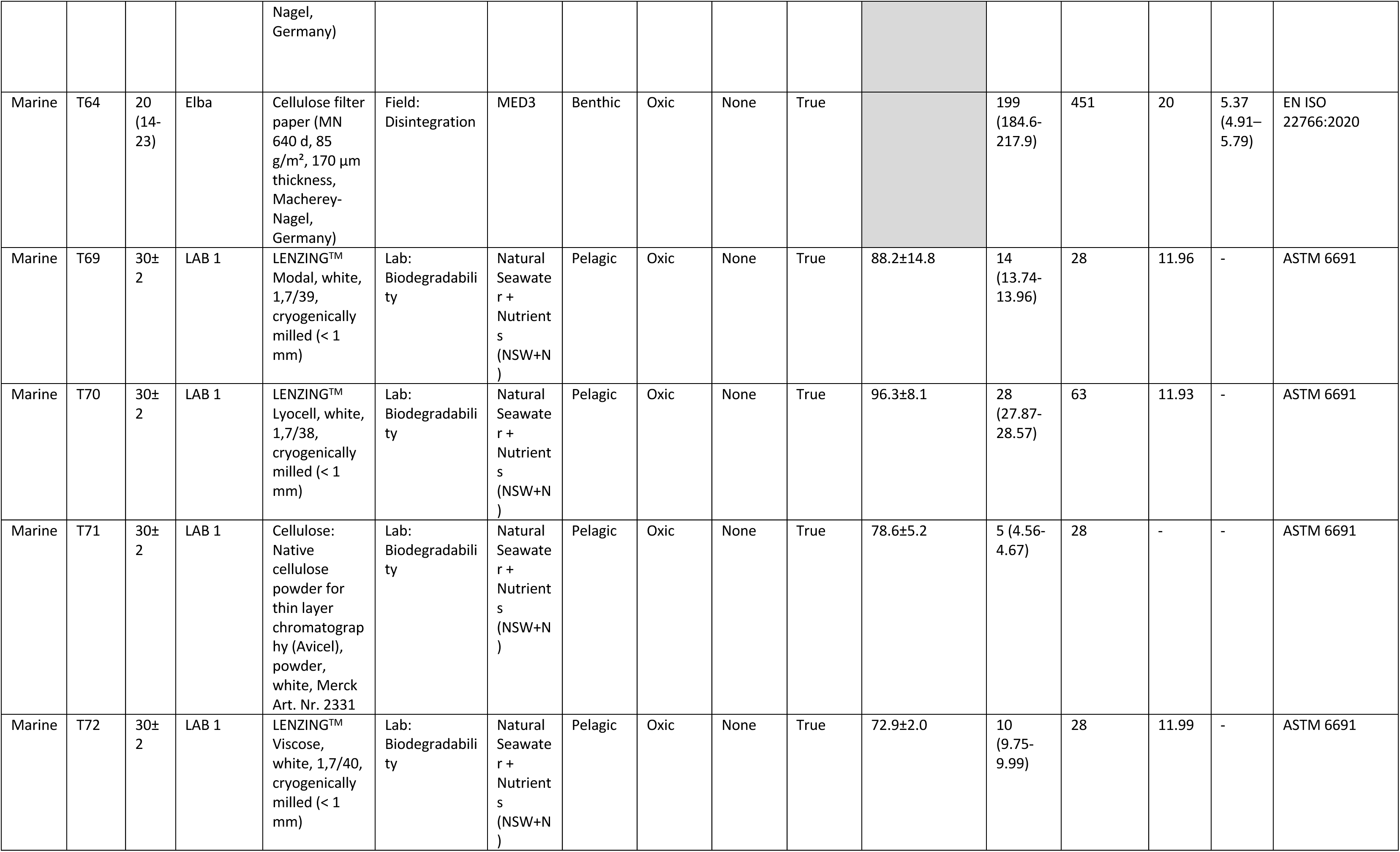
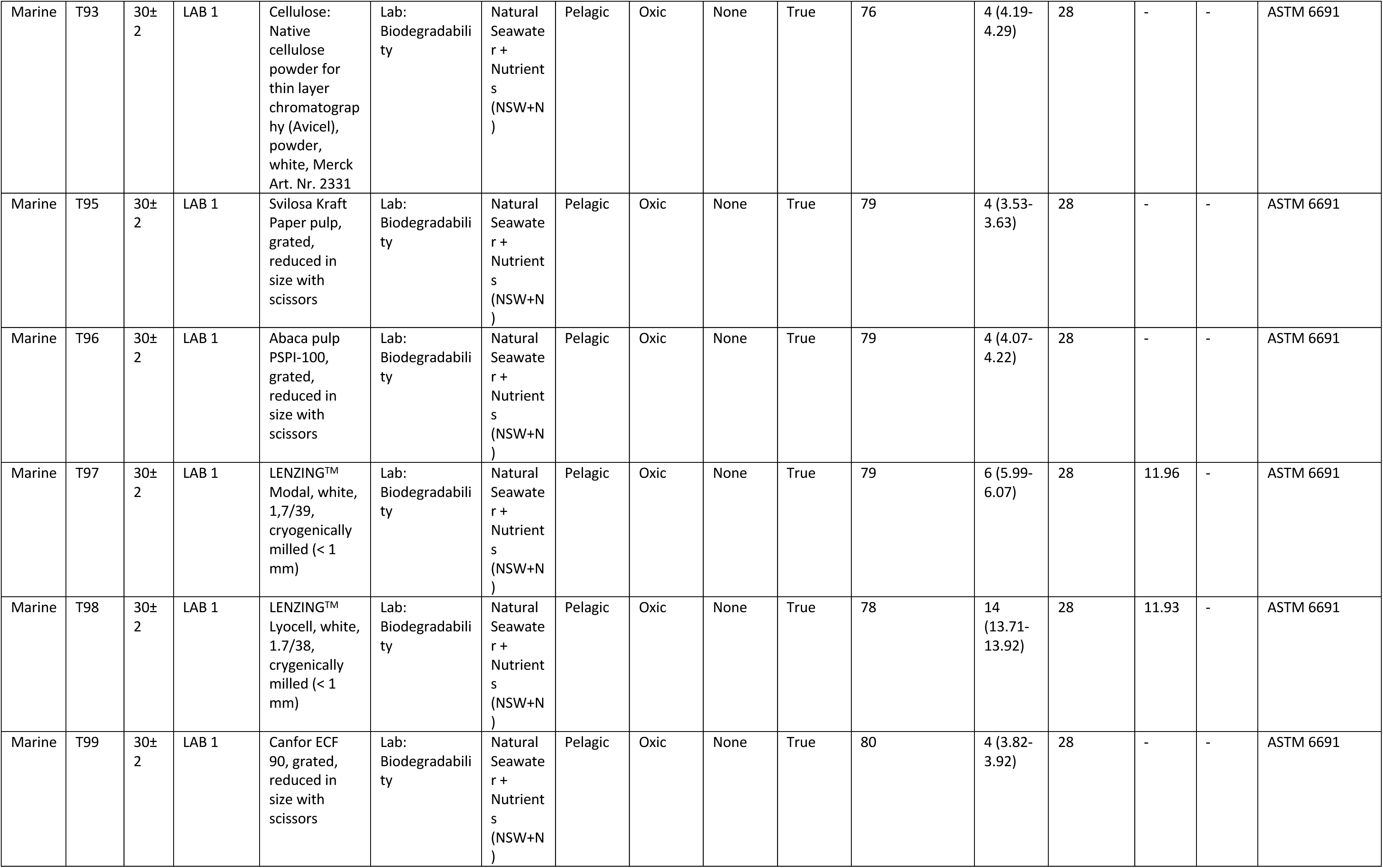
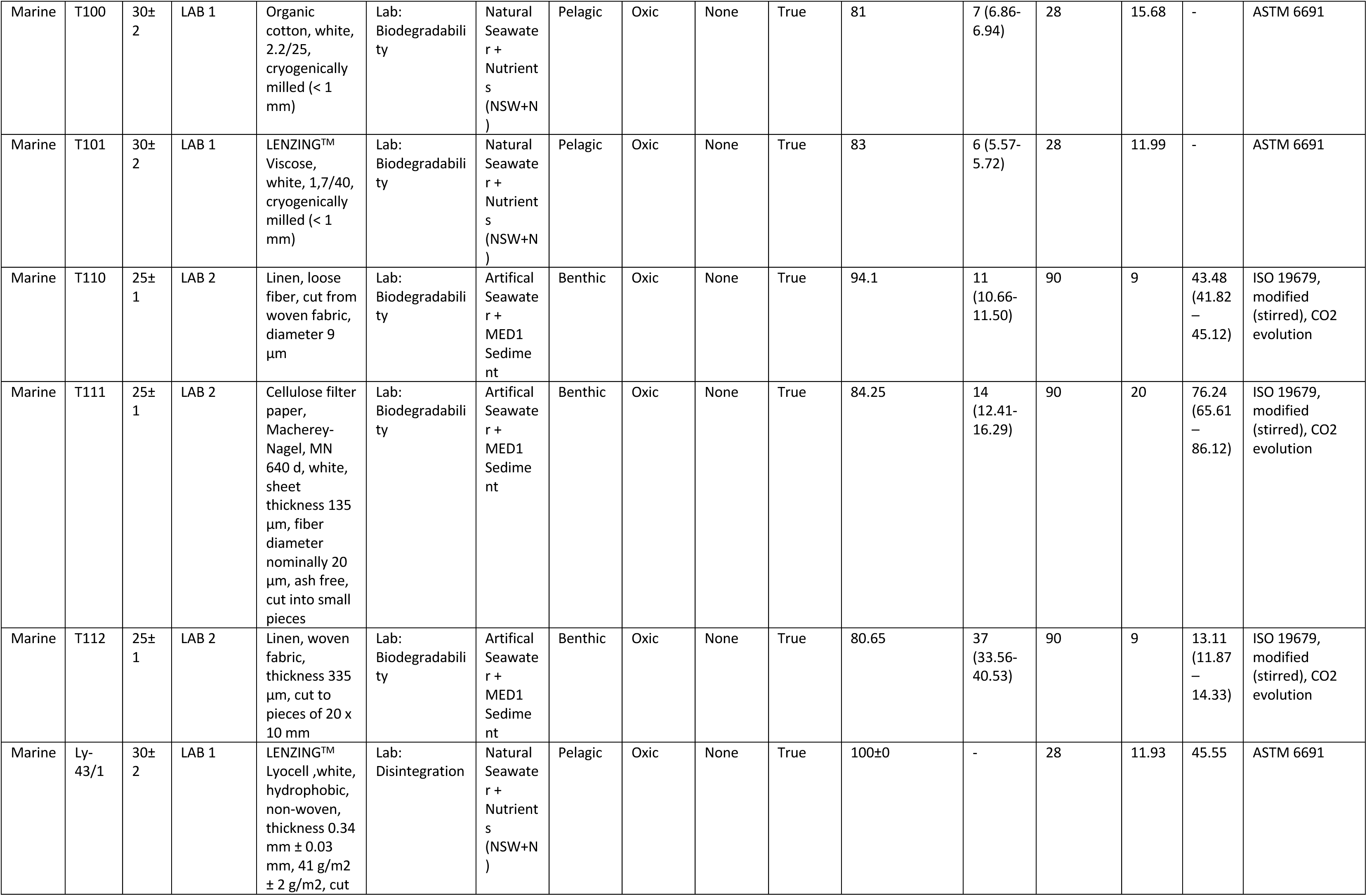
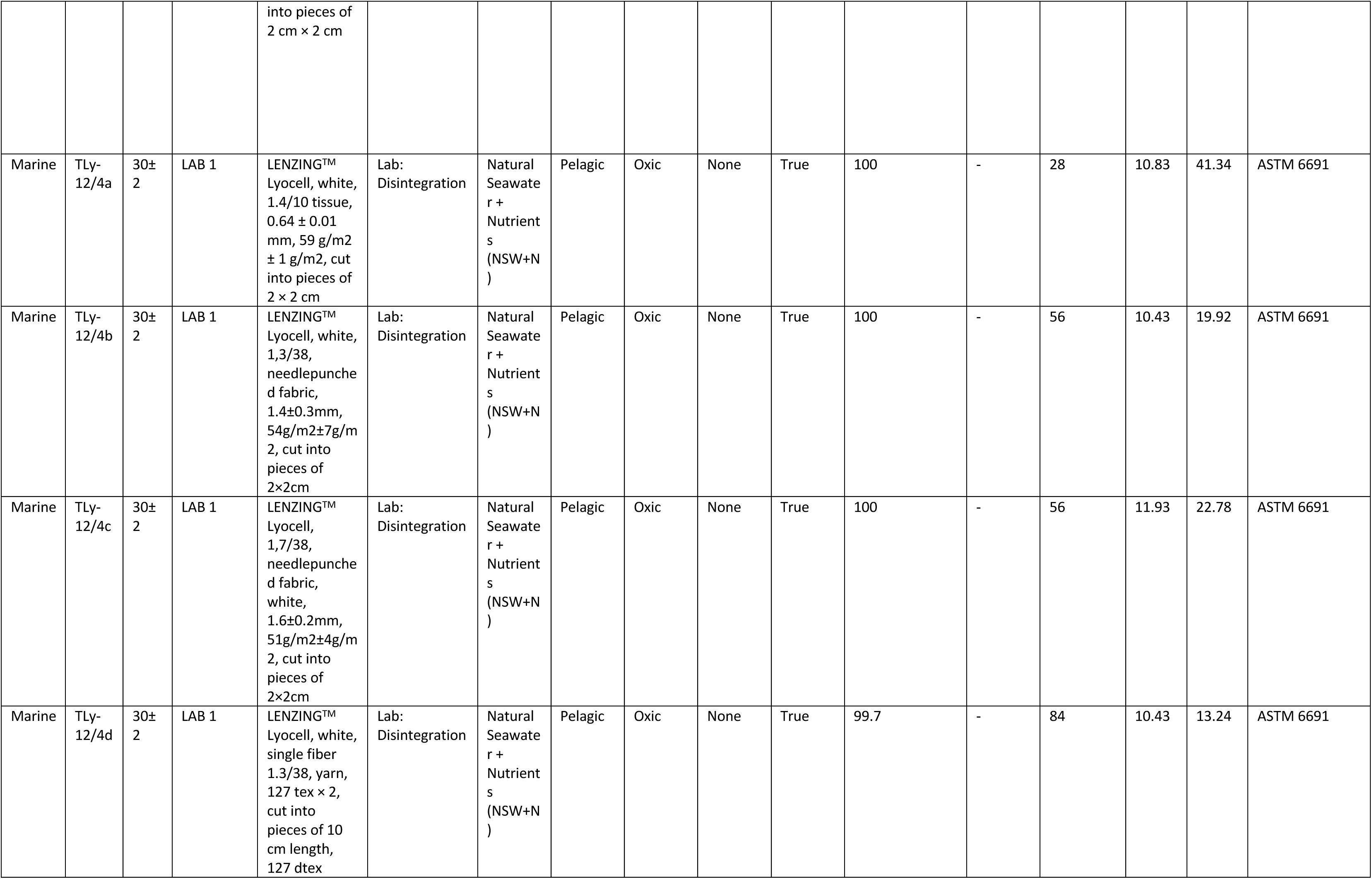
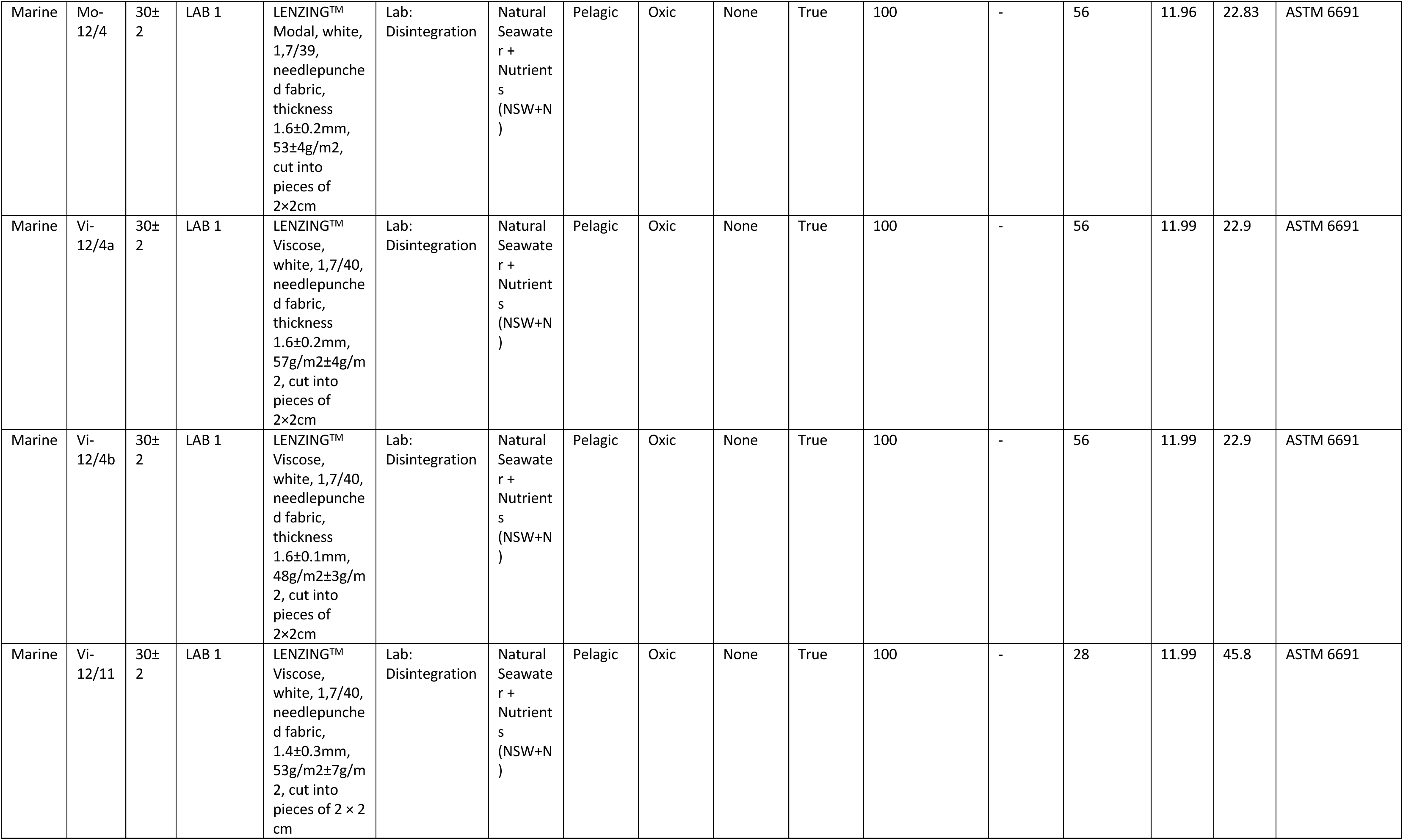
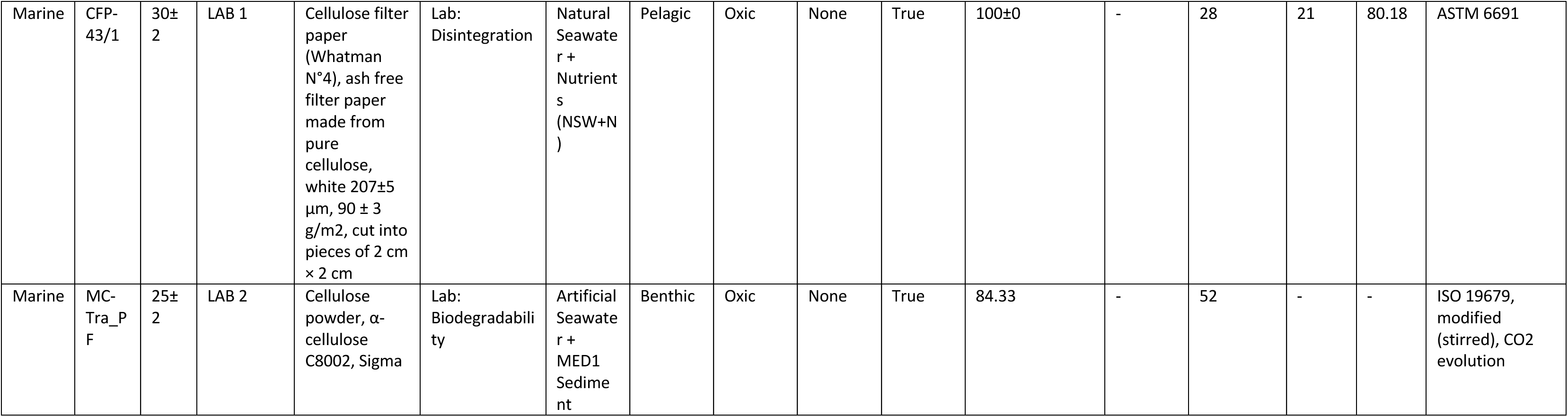
Overview of materials and experimental treatments under marine conditions. The materials were exposed to various marine environments to investigate the effects of environmental factors on their degree of disintegration or biodegradation (%). Each treatment (coded as Tn) represented a unique combination of temperature (°C), location, material, test type, matrix, vertical position, oxygen availability, particularities and sample type. The half-life and SSDR were determined where data quality allowed. Information on sample exposure (days), fiber diameter (µm) and the applied standard is also provided.

| Treat | Code | T [°C] | Location | Material | Test Type | Matrix | Vertical position | Oxygen | Particularities | Sample Type | Disintegration or Biodegradation [%] | Half-Life [d (CI)] | Exposure [d] | Fiber Ø [µm] | SSDR [µm yr<sup>-1</sup> (CI)] | Standard |
| --- | --- | --- | --- | --- | --- | --- | --- | --- | --- | --- | --- | --- | --- | --- | --- | --- |
| Marine | T11 | 23 | LAB 2 | Cellulose filter paper (MN 640 d, 85 g/m<sup>2</sup>, 170 µm thickness, Macherey-Nagel, Germany) | Tank: Disintegration | MED1 | Benthic | Oxic | None | Check |  | 68 (53.62-82.19) | 202 | 20 | 15.74 (13.01 – 19.94) | ISO 23832:2021 |
| Marine | T12 | 23 | LAB 2 | Cellulose filter paper (MN 640 d, 85 g/m<sup>2</sup>, 170 µm thickness, Macherey-Nagel, Germany) | Tank: Disintegration | MED2 | Benthic | Oxic | None | Check |  | 144 (124.41-162.73) | 202 | 20 | 7.43 (6.57–8.59) | ISO 23832:2021 |
| Marine | T13 | 23 | LAB 2 | Cellulose filter paper (MN 640 d, 85 g/m<sup>2</sup>, 170 µm thickness, Macherey-Nagel, Germany) | Tank: Disintegration | MED4 | Benthic | Oxic | None | Check | 0% after 202 d | - | 202 | 20 | - | ISO 23832:2021 |
| Marine | T14 | 23 | LAB 2 | Cellulose filter paper (MN 640 d, 85 g/m<sup>2</sup>, 170 µm thickness, Macherey- | Tank: Disintegration | MED6 | Benthic | Oxic | None | Check | 36% after 202 d | - | 202 | 20 | - | ISO 23832:2021 |
|  |  |  |  | Nagel,<br>Germany) |  |  |  |  |  |  |  |  |  |  |  |  |
| Marine | T15 | 23 | LAB 2 | Cellulose filter<br>paper (MN<br>640 d, 85<br>g/m<sup>2</sup>, 170 µm<br>thickness,<br>Macherey-Nagel,<br>Germany) | Tank:<br>Disintegration | MED5 | Benthic | Oxic | None | Check |  | 189<br>(157.28-238.42) | 202 | 20 | 5.66<br>(4.48–<br>6.8) | ISO 23832:2021 |
| Marine | T16 | 23 | LAB 2 | Cellulose filter<br>paper (MN<br>640 d, 85<br>g/m<sup>2</sup>, 170 µm<br>thickness,<br>Macherey-Nagel,<br>Germany) | Tank:<br>Disintegration | MED7 | Benthic | Oxic | None | Check | 44% after 202 d | - | 202 | 20 | - | ISO 23832:2021 |
| Marine | T17 | 23 | LAB 2 | Cellulose filter<br>paper (MN<br>640 d, 85<br>g/m<sup>2</sup>, 170 µm<br>thickness,<br>Macherey-Nagel,<br>Germany) | Tank:<br>Disintegration | NOR3 | Benthic | Oxic | None | Check |  | 58<br>(54.21-62.15) | 137 | 20 | 20.38<br>(16.73<br>–<br>25.53) | ISO 23832:2021 |
| Marine | T19 | 23 | LAB 2 | Cellulose filter<br>paper (MN<br>640 d, 85<br>g/m<sup>2</sup>, 170 µm<br>thickness,<br>Macherey-Nagel,<br>Germany) | Tank:<br>Disintegration | NOR1 | Benthic | Oxic | None | Check |  | 15<br>(13.57-16.9) | 22 | 20 | 23.37<br>(21.38<br>–<br>25.32) | ISO 23832:2021 |
| Marine | T20 | 23 | LAB 2 | Cellulose filter<br>paper (MN<br>640 d, 85<br>g/m<sup>2</sup>, 170 µm<br>thickness,<br>Macherey-Nagel,<br>Germany) | Tank:<br>Disintegration | NOR4 | Benthic | Oxic | None | Check |  | 34<br>(31.86-36.56) | 56 | 20 | 45.38<br>(21.01<br>–<br>71.22) | ISO 23832:2021 |
| Marine | T44 | 9 | Bergen Aquarium | Cellulose filter paper (MN 640 d, 85 g/m<sup>2</sup>, 170 µm thickness, Macherey-Nagel, Germany) | Tank: Disintegration | NOR5 | Benthic | Oxic | None | Check |  | 58 (54.55-61.55) | 75 | 20 | 18.34 (17.37-19.6) | ISO 23832:2021 |
| Marine | T45 | 9 | Bergen Aquarium | Cellulose filter paper (MN 640 d, 85 g/m<sup>2</sup>, 170 µm thickness, Macherey-Nagel, Germany) | Tank: Disintegration | NOR5 | Pelagic | Oxic | None | Check |  | 130 (114.55-149.26) | 147 | 20 | 8.25 (7.16-9.33) | ISO 23832:2021 |
| Marine | T47 | 28 | LAB 2 | Cellulose filter paper (MN 640 d, 85 g/m<sup>2</sup>, 170 µm thickness, Macherey-Nagel, Germany) | Tank: Disintegration | SEA1 | Benthic | Oxic | None | Check |  | 93 (85.54-101.08) | 138 | 20 | 11.45 (10.58-12.5) | ISO 23832:2021 |
| Marine | T48 | 28 | LAB 2 | Cellulose filter paper (MN 640 d, 85 g/m<sup>2</sup>, 170 µm thickness, Macherey-Nagel, Germany) | Tank: Disintegration | SEA1 | Benthic | Oxic | Weathered samples | Check |  | 90 (65.71-153.66) | 76 | 20 | 11.93 (6.95-16.26) | ISO 23832:2021 |
| Marine | T49 | 28 | LAB 2 | Cellulose filter paper (MN 640 d, 85 g/m<sup>2</sup>, 170 µm thickness, Macherey-Nagel, Germany) | Tank: Disintegration | MED3 | Benthic | Oxic | None | Check |  | 159 (136.24-195.78) | 140 | 20 | 6.71 (5.46-7.85) | ISO 23832:2021 |
| Marine | T50 | 14 | LAB 2 | Cellulose filter paper (MN 640 d, 85 g/m<sup>2</sup>, 170 µm thickness, Macherey- | Tank: Disintegration | MED3 | Benthic | Oxic | None | Check |  | 389 (267.49-843.8) | 399 | 20 | 2.75 (1.23-3.97) | ISO 23832:2021 |
|  |  |  |  | Nagel,<br>Germany) |  |  |  |  |  |  |  |  |  |  |  |  |
| Marine | T53 | 26 | LAB 2 | Cellulose filter<br>paper (MN<br>640 d, 85<br>g/m<sup>2</sup>, 170 µm<br>thickness,<br>Macherey-Nagel,<br>Germany) | Tank:<br>Disintegration | NOR4 | Burried | Anoxic | None | Check |  | 166<br>(153.73-180.29) | 204 | 20 | 6.45<br>(5.93–<br>6.95) | ISO 23832:2021 |
| Marine | T54 | 29<br>(23.3-31.5<br>) | Bangka | Cellulose filter<br>paper (MN<br>640 d, 85<br>g/m<sup>2</sup>, 170 µm<br>thickness,<br>Macherey-Nagel,<br>Germany) | Field:<br>Disintegration | SEA3 | Benthic | Oxic | None | True |  | 34<br>(22.04-52.88) | 127 | 20 | 31.91<br>(20.02<br>–<br>48.13) | EN ISO<br>22766:2020 |
| Marine | T55 | 29<br>(26-32.2<br>) | Bangka | Cellulose filter<br>paper (MN<br>640 d, 85<br>g/m<sup>2</sup>, 170 µm<br>thickness,<br>Macherey-Nagel,<br>Germany) | Field:<br>Disintegration | SEA2 | Buried<br>at 0.5 m | Fluctuat<br>ing | Eulittora<br>l | True |  | 90<br>(78.93-102.83) | 988 | 20 | 11.83<br>(10.4–<br>13.54) | EN ISO<br>22766:2020 |
| Marine | T56 | 29<br>(23.3-31.5<br>) | Bangka | Cellulose filter<br>paper (MN<br>640 d, 85<br>g/m<sup>2</sup>, 170 µm<br>thickness,<br>Macherey-Nagel,<br>Germany) | Field:<br>Disintegration | SEA3 | Pelagic<br>at 20 m | Oxic | None | True |  | 97<br>(90.91-115.35) | 86 | 20 | 10.99<br>(9.26–<br>11.76) | acc. to Lott et<br>al., 2020, similar<br>to ISO<br>15314:2018 |
| Marine | T57 | 28 | LAB 2 | Cellulose filter<br>paper (MN<br>640 d, 85<br>g/m<sup>2</sup>, 170 µm<br>thickness,<br>Macherey-Nagel,<br>Germany) | Tank:<br>Disintegration | MED2 | Benthic | Oxic | Fresh<br>sedimen<br>t | Check |  | 36<br>(24.1-46.71) | 103 | 20 | 29.3<br>(22.89<br>–<br>44.33) | ISO 23832:2021 |
| Marine | T58 | 28 | LAB 2 | Cellulose filter paper (MN 640 d, 85 g/m<sup>2</sup>, 170 µm thickness, Macherey-Nagel, Germany) | Tank: Disintegration | MED2 | Benthic | Oxic | Short sediment storage + aeration | Check |  | 42 (35.89-48.51) | 68 | 20 | 25.33 (22.04 – 29.79) | ISO 23832:2021 |
| Marine | T59 | 28 | LAB 2 | Cellulose filter paper (MN 640 d, 85 g/m<sup>2</sup>, 170 µm thickness, Macherey-Nagel, Germany) | Tank: Disintegration | MED2 | Benthic | Oxic | Short sediment storage + no aeration | Check |  | 53 (44.68-59.26) | 82 | 20 | 20.02 (18.04 – 23.93) | ISO 23832:2021 |
| Marine | T60 | 28 | LAB 2 | Cellulose filter paper (MN 640 d, 85 g/m<sup>2</sup>, 170 µm thickness, Macherey-Nagel, Germany) | Tank: Disintegration | MED2 | Benthic | Oxic | Long sediment storage + aeration | Check |  | 25 (19.56-29.61) | 53 | 20 | 42.87 (36.1–54.65) | ISO 23832:2021 |
| Marine | T61 | 28 | LAB 2 | Cellulose filter paper (MN 640 d, 85 g/m<sup>2</sup>, 170 µm thickness, Macherey-Nagel, Germany) | Tank: Disintegration | MED2 | Benthic | Oxic | Long sediment storage + no aeration | Check |  | 80 (63.42-113.14) | 89 | 20 | 13.38 (9.45–16.85) | ISO 23832:2021 |
| Marine | T62 | 26 (-1-38.4) | Elba | Cellulose filter paper (MN 640 d, 85 g/m<sup>2</sup>, 170 µm thickness, Macherey-Nagel, Germany) | Field: Disintegration | MED8 | Buried at 0.1 m | Fluctuating | Eulittoral | True |  | 259 (233.03-290.13) | 298 | 20 | 4.13 (3.68–4.59) | EN ISO 22766:2020 |
| Marine | T63 | 25 (14-27) | Elba | Cellulose filter paper (MN 640 d, 85 g/m<sup>2</sup>, 170 µm thickness, Macherey- | Field: Disintegration | MED9 | Benthic | Oxic | Lagoon | True |  | 29 (26-31.71) | 82 | 20 | 37.42 (33.72 – 41.12) | EN ISO 22766:2020 |
|  |  |  |  | Nagel,<br>Germany) |  |  |  |  |  |  |  |  |  |  |  |  |
| Marine | T64 | 20<br>(14-23) | Elba | Cellulose filter paper (MN 640 d, 85 g/m<sup>2</sup>, 170 µm thickness, Macherey-Nagel, Germany) | Field: Disintegration | MED3 | Benthic | Oxic | None | True |  | 199<br>(184.6-217.9) | 451 | 20 | 5.37<br>(4.91–5.79) | EN ISO 22766:2020 |
| Marine | T69 | 30±2 | LAB 1 | LENZING™ Modal, white, 1,7/39, cryogenically milled (< 1 mm) | Lab: Biodegradability | Natural Seawater + Nutrients (NSW+N) | Pelagic | Oxic | None | True | 88.2±14.8 | 14<br>(13.74-13.96) | 28 | 11.96 | - | ASTM 6691 |
| Marine | T70 | 30±2 | LAB 1 | LENZING™ Lyocell, white, 1,7/38, cryogenically milled (< 1 mm) | Lab: Biodegradability | Natural Seawater + Nutrients (NSW+N) | Pelagic | Oxic | None | True | 96.3±8.1 | 28<br>(27.87-28.57) | 63 | 11.93 | - | ASTM 6691 |
| Marine | T71 | 30±2 | LAB 1 | Cellulose: Native cellulose powder for thin layer chromatography (Avicel), powder, white, Merck Art. Nr. 2331 | Lab: Biodegradability | Natural Seawater + Nutrients (NSW+N) | Pelagic | Oxic | None | True | 78.6±5.2 | 5 (4.56-4.67) | 28 | - | - | ASTM 6691 |
| Marine | T72 | 30±2 | LAB 1 | LENZING™ Viscose, white, 1,7/40, cryogenically milled (< 1 mm) | Lab: Biodegradability | Natural Seawater + Nutrients (NSW+N) | Pelagic | Oxic | None | True | 72.9±2.0 | 10<br>(9.75-9.99) | 28 | 11.99 | - | ASTM 6691 |
| Marine | T93 | 30±<br>2 | LAB 1 | Cellulose:<br>Native<br>cellulose<br>powder for<br>thin layer<br>chromatograp<br>hy (Avicel),<br>powder,<br>white, Merck<br>Art. Nr. 2331 | Lab:<br>Biodegradabili<br>ty | Natural<br>Seawate<br>r +<br>Nutrient<br>s<br>(NSW+N<br>) | Pelagic | Oxic | None | True | 76 | 4 (4.19-4.29) | 28 | - | - | ASTM 6691 |
| Marine | T95 | 30±<br>2 | LAB 1 | Svilosa Kraft<br>Paper pulp,<br>grated,<br>reduced in<br>size with<br>scissors | Lab:<br>Biodegradabili<br>ty | Natural<br>Seawate<br>r +<br>Nutrient<br>s<br>(NSW+N<br>) | Pelagic | Oxic | None | True | 79 | 4 (3.53-3.63) | 28 | - | - | ASTM 6691 |
| Marine | T96 | 30±<br>2 | LAB 1 | Abaca pulp<br>PSPI-100,<br>grated,<br>reduced in<br>size with<br>scissors | Lab:<br>Biodegradabili<br>ty | Natural<br>Seawate<br>r +<br>Nutrient<br>s<br>(NSW+N<br>) | Pelagic | Oxic | None | True | 79 | 4 (4.07-4.22) | 28 | - | - | ASTM 6691 |
| Marine | T97 | 30±<br>2 | LAB 1 | LENZING™<br>Modal, white,<br>1,7/39,<br>cryogenically<br>milled (< 1<br>mm) | Lab:<br>Biodegradabili<br>ty | Natural<br>Seawate<br>r +<br>Nutrient<br>s<br>(NSW+N<br>) | Pelagic | Oxic | None | True | 79 | 6 (5.99-6.07) | 28 | 11.96 | - | ASTM 6691 |
| Marine | T98 | 30±<br>2 | LAB 1 | LENZING™<br>Lyocell, white,<br>1.7/38,<br>crygenically<br>milled (< 1<br>mm) | Lab:<br>Biodegradabili<br>ty | Natural<br>Seawate<br>r +<br>Nutrient<br>s<br>(NSW+N<br>) | Pelagic | Oxic | None | True | 78 | 14<br>(13.71-13.92) | 28 | 11.93 | - | ASTM 6691 |
| Marine | T99 | 30±<br>2 | LAB 1 | Canfor ECF<br>90, grated,<br>reduced in<br>size with<br>scissors | Lab:<br>Biodegradabili<br>ty | Natural<br>Seawate<br>r +<br>Nutrient<br>s<br>(NSW+N<br>) | Pelagic | Oxic | None | True | 80 | 4 (3.82-3.92) | 28 | - | - | ASTM 6691 |
| Marine | T100 | 30±2 | LAB 1 | Organic cotton, white, 2.2/25, cryogenically milled (< 1 mm) | Lab: Biodegradability | Natural Seawater + Nutrients (NSW+N) | Pelagic | Oxic | None | True | 81 | 7 (6.86-6.94) | 28 | 15.68 | - | ASTM 6691 |
| Marine | T101 | 30±2 | LAB 1 | LENZING™ Viscose, white, 1,7/40, cryogenically milled (< 1 mm) | Lab: Biodegradability | Natural Seawater + Nutrients (NSW+N) | Pelagic | Oxic | None | True | 83 | 6 (5.57-5.72) | 28 | 11.99 | - | ASTM 6691 |
| Marine | T110 | 25±1 | LAB 2 | Linen, loose fiber, cut from woven fabric, diameter 9 µm | Lab: Biodegradability | Artificial Seawater + MED1 Sediment | Benthic | Oxic | None | True | 94.1 | 11 (10.66-11.50) | 90 | 9 | 43.48 (41.82 – 45.12) | ISO 19679, modified (stirred), CO2 evolution |
| Marine | T111 | 25±1 | LAB 2 | Cellulose filter paper, Macherey-Nagel, MN 640 d, white, sheet thickness 135 µm, fiber diameter nominally 20 µm, ash free, cut into small pieces | Lab: Biodegradability | Artificial Seawater + MED1 Sediment | Benthic | Oxic | None | True | 84.25 | 14 (12.41-16.29) | 90 | 20 | 76.24 (65.61 – 86.12) | ISO 19679, modified (stirred), CO2 evolution |
| Marine | T112 | 25±1 | LAB 2 | Linen, woven fabric, thickness 335 µm, cut to pieces of 20 x 10 mm | Lab: Biodegradability | Artificial Seawater + MED1 Sediment | Benthic | Oxic | None | True | 80.65 | 37 (33.56-40.53) | 90 | 9 | 13.11 (11.87 – 14.33) | ISO 19679, modified (stirred), CO2 evolution |
| Marine | Ly-43/1 | 30±2 | LAB 1 | LENZING™ Lyocell, white, hydrophobic, non-woven, thickness 0.34 mm ± 0.03 mm, 41 g/m2 ± 2 g/m2, cut into pieces of<br>2 cm × 2 cm | Lab: Disintegration | Natural Seawater + Nutrients (NSW+N) | Pelagic | Oxic | None | True | 100±0 | - | 28 | 11.93 | 45.55 | ASTM 6691 |
| Marine | Tly-12/4a | 30±2 | LAB 1 | LENZING™<br>Lyocell, white,<br>1.4/10 tissue,<br>0.64 ± 0.01<br>mm, 59 g/m2<br>± 1 g/m2, cut<br>into pieces of<br>2 × 2 cm | Lab:<br>Disintegration | Natural<br>Seawater<br>+<br>Nutrient<br>s<br>(NSW+N<br>) | Pelagic | Oxic | None | True | 100 | - | 28 | 10.83 | 41.34 | ASTM 6691 |
| Marine | Tly-12/4b | 30±2 | LAB 1 | LENZING™<br>Lyocell, white,<br>1,3/38,<br>needlepunch<br>ed fabric,<br>1.4±0.3mm,<br>54g/m2±7g/m<br>2, cut into<br>pieces of<br>2x2cm | Lab:<br>Disintegration | Natural<br>Seawater<br>+<br>Nutrient<br>s<br>(NSW+N<br>) | Pelagic | Oxic | None | True | 100 | - | 56 | 10.43 | 19.92 | ASTM 6691 |
| Marine | Tly-12/4c | 30±2 | LAB 1 | LENZING™<br>Lyocell,<br>1,7/38,<br>needlepunch<br>ed fabric,<br>white,<br>1.6±0.2mm,<br>51g/m2±4g/m<br>2, cut into<br>pieces of<br>2x2cm | Lab:<br>Disintegration | Natural<br>Seawater<br>+<br>Nutrient<br>s<br>(NSW+N<br>) | Pelagic | Oxic | None | True | 100 | - | 56 | 11.93 | 22.78 | ASTM 6691 |
| Marine | Tly-12/4d | 30±2 | LAB 1 | LENZING™<br>Lyocell, white,<br>single fiber<br>1.3/38, yarn,<br>127 tex × 2,<br>cut into<br>pieces of 10<br>cm length,<br>127 dtex | Lab:<br>Disintegration | Natural<br>Seawater<br>+<br>Nutrient<br>s<br>(NSW+N<br>) | Pelagic | Oxic | None | True | 99.7 | - | 84 | 10.43 | 13.24 | ASTM 6691 |
| Marine | Mo-12/4 | 30±2 | LAB 1 | LENZING™ Modal, white, 1,7/39, needlepunched fabric, thickness 1.6±0.2mm, 53±4g/m<sup>2</sup>, cut into pieces of 2×2cm | Lab: Disintegration | Natural Seawater + Nutrients (NSW+N) | Pelagic | Oxic | None | True | 100 | - | 56 | 11.96 | 22.83 | ASTM 6691 |
| Marine | Vi-12/4a | 30±2 | LAB 1 | LENZING™ Viscose, white, 1,7/40, needlepunched fabric, thickness 1.6±0.2mm, 57g/m<sup>2</sup>±4g/m<sup>2</sup>, cut into pieces of 2×2cm | Lab: Disintegration | Natural Seawater + Nutrients (NSW+N) | Pelagic | Oxic | None | True | 100 | - | 56 | 11.99 | 22.9 | ASTM 6691 |
| Marine | Vi-12/4b | 30±2 | LAB 1 | LENZING™ Viscose, white, 1,7/40, needlepunched fabric, thickness 1.6±0.1mm, 48g/m<sup>2</sup>±3g/m<sup>2</sup>, cut into pieces of 2×2cm | Lab: Disintegration | Natural Seawater + Nutrients (NSW+N) | Pelagic | Oxic | None | True | 100 | - | 56 | 11.99 | 22.9 | ASTM 6691 |
| Marine | Vi-12/11 | 30±2 | LAB 1 | LENZING™ Viscose, white, 1,7/40, needlepunched fabric, 1.4±0.3mm, 53g/m<sup>2</sup>±7g/m<sup>2</sup>, cut into pieces of 2 × 2 cm | Lab: Disintegration | Natural Seawater + Nutrients (NSW+N) | Pelagic | Oxic | None | True | 100 | - | 28 | 11.99 | 45.8 | ASTM 6691 |
| Marine | CFP-43/1 | 30±2 | LAB 1 | Cellulose filter paper (Whatman N°4), ash free filter paper made from pure cellulose, white 207±5 µm, 90 ± 3 g/m2, cut into pieces of 2 cm × 2 cm | Lab: Disintegration | Natural Seawater + Nutrients (NSW+N) | Pelagic | Oxic | None | True | 100±0 | - | 28 | 21 | 80.18 | ASTM 6691 |
| Marine | MC-Tra_P F | 25±2 | LAB 2 | Cellulose powder, α-cellulose C8002, Sigma | Lab: Biodegradability | Artificial Seawater + MED1 Sediment | Benthic | Oxic | None | True | 84.33 | - | 52 | - | - | ISO 19679, modified (stirred), CO2 evolution |

**Table SI5:** Overview of materials and experimental treatments where only observational data were available. The materials were exposed to various environments to investigate the effects of environmental factors on their degree of disintegration (%). Each treatment (coded as Tn) represented a unique combination of temperature (°C), location, material, test type, matrix, vertical position, oxygen availability and sample type. The half-life was not determined due to insufficient data quality. Information on sample exposure (days) and the applied standard is also provided.

| Treat | T [°C] | Location | Material | Test Type | Matrix | Vertical position | Oxygen | Sample Type | Disintegration [%] | Half-Life [d (CI)] | Exposure [d] | Standard |
| --- | --- | --- | --- | --- | --- | --- | --- | --- | --- | --- | --- | --- |
| Marine: Deep-sea | 14 | Mediterranean Sea, Sicily Channel | Cellulose filter paper | Field: Disintegration | DPS1 | Pelagic (at 532 m) | Oxic | True | 100 | - | 612 | acc. to Lott et al., 2020, similar to ISO 15314:2018 |
| Marine: Deep-sea | 14 | Mediterranean Sea, Corsica Channel | Cellulose filter paper | Field: Disintegration | DPS2 | Pelagic (at 400 m) | Oxic | True | 100 | - | 707 | acc. to Lott et al., 2020, similar to ISO 15314:2018 |
| Marine: Deep-sea | 14 | Mediterranean Sea, Sicily Channel | Cotton fabric | Field: Disintegration | DPS1 | Pelagic (at 532 m) | Oxic | True | 95 | - | 612 | acc. to Lott et al., 2020, similar to ISO 15314:2018 |
| Marine: Deep-sea | 14 | Mediterranean Sea, Corsica Channel | Cotton fabric | Field: Disintegration | DPS2 | Pelagic (at 400 m) | Oxic | True | 100 | - | 707 | acc. to Lott et al., 2020, similar to ISO 15314:2018 |
| Marine: Deep-sea | 14 | Mediterranean Sea, Sicily Channel | Lyocell | Field: Disintegration | DPS1 | Pelagic (at 532 m) | Oxic | True | 80 | - | 612 | acc. to Lott et al., 2020, similar to ISO 15314:2018 |
| Marine: Deep-sea | 14 | Mediterranean Sea, Corsica Channel | Lyocell | Field: Disintegration | DPS2 | Pelagic (at 400 m) | Oxic | True | 100 | - | 707 | acc. to Lott et al., 2020, similar to ISO 15314:2018 |
| Marine | 4 - 35 | Mediterranean Sea, Isola d'Elba | Cellulose filter paper | Field: Disintegration | MED8 | Eulittoral (Mid-water line) | Oxic | True | 95 | - | 258 | EN ISO 22766:2020 |
| Marine | 4 - 35 | Mediterranean Sea, Isola d'Elba | Cellulose filter paper | Field: Disintegration | MED8 | Eulittoral (Mid-water line) | Oxic | True | 100 | - | 673 | EN ISO 22766:2020 |
| Marine | 4 - 35 | Mediterranean Sea, Isola d'Elba | Cotton fabric | Field: Disintegration | MED8 | Eulittoral (Mid-water line) | Oxic | True | 0 | - | 258 | EN ISO 22766:2020 |
| Marine | 4 - 35 | Mediterranean Sea, Isola d'Elba | Cotton fabric | Field: Disintegration | MED8 | Eulittoral (Mid-water line) | Oxic | True | 100 | - | 673 | EN ISO 22766:2020 |
| Marine | 4 - 35 | Mediterranean Sea, Isola d'Elba | Lyocell | Field: Disintegration | MED8 | Eulittoral (Mid-water line) | Oxic | True | 0 | - | 258 | EN ISO 22766:2020 |
| Marine | 4 - 35 | Mediterranean Sea, Isola d'Elba | Lyocell | Field: Disintegration | MED8 | Eulittoral (Mid-water line) | Oxic | True | 85 | - | 673 | EN ISO 22766:2020 |
| Marine | 17 - 27 | Mediterranean Sea, Isola d'Elba | Cellulose filter paper | Field: Disintegration | MED9 | Lagoon (at 1 m) | Oxic | True | 100 | - | 257 | EN ISO 22766:2020 |
| Marine | 14 - 25 | Mediterranean Sea, Isola d'Elba | Cellulose filter paper | Field: Disintegration | MED3 | Benthic (40 m) | Oxic | True | 100 | - | 415 | EN ISO 22766:2020 |
| Marine | 17 - 27 | Mediterranean Sea, Isola d'Elba | Cotton fabric | Field: Disintegration | MED9 | Lagoon (at 1 m) | Oxic | True | 100 | - | 257 | EN ISO 22766:2020 |
| Marine | 14 - 25 | Mediterranean Sea, Isola d'Elba | Cotton fabric | Field: Disintegration | MED3 | Benthic (40 m) | Oxic | True | 90 | - | 415 | EN ISO 22766:2020 |
| Marine | 17 - 27 | Mediterranean Sea, Isola d'Elba | Lyocell | Field: Disintegration | MED9 | Lagoon (at 1 m) | Oxic | True | 100 | - | 257 | EN ISO 22766:2020 |
| Marine | 14 - 25 | Mediterranean Sea, Isola d'Elba | Lyocell | Field: Disintegration | MED3 | Benthic (40 m) | Oxic | True | 100 | - | 415 | EN ISO 22766:2020 |
| Marine | 29 | Bangka | Cellulose filter paper | Field: Disintegration | SEA2 | Eulittoral (Buried at 50 cm) | Fluctuating | True | 10 | - | 103 | EN ISO 22766:2020 |
| Marine | 29 | Bangka | Cellulose filter paper | Field: Disintegration | SEA2 | Eulittoral (Buried at 50 cm) | Fluctuating | True | 100 | - | 207 | EN ISO 22766:2020 |
| Marine | 29 | Bangka | Cellulose filter paper | Field: Disintegration | SEA3 | Benthic (at 32 m) | Oxic | True | 30 | - | 102 | EN ISO 22766:2020 |
| Marine | 29 | Bangka | Cellulose filter paper | Field: Disintegration | SEA3 | Benthic (at 32 m) | Oxic | True | 100 | - | 205 | EN ISO 22766:2020 |
| Marine | 29 | Bangka | Cellulose filter paper | Field: Disintegration | SEA3 | Benthic (at 32 m) | Oxic | True | 100 | - | 1312 | EN ISO 22766:2020 |
| Marine | 29 | Bangka | Cotton fabric | Field: Disintegration | SEA2 | Eulittoral (Buried at 50 cm) | Fluctuating | True | 0 | - | 103 | EN ISO 22766:2020 |
| Marine | 29 | Bangka | Cotton fabric | Field: Disintegration | SEA2 | Eulittoral (Buried at 50 cm) | Fluctuating | True | 100 | - | 207 | EN ISO 22766:2020 |
| Marine | 29 | Bangka | Cotton fabric | Field: Disintegration | SEA3 | Benthic (at 32 m) | Oxic | True | 0 | - | 102 | EN ISO 22766:2020 |
| Marine | 29 | Bangka | Cotton fabric | Field: Disintegration | SEA3 | Benthic (at 32 m) | Oxic | True | 45 | - | 205 | EN ISO 22766:2020 |
| Marine | 29 | Bangka | Cotton fabric | Field: Disintegration | SEA3 | Benthic (at 32 m) | Oxic | True | 100 | - | 1312 | EN ISO 22766:2020 |
| Marine | 29 | Bangka | Organic cotton t-shirt | Field: Disintegration | SEA3 | Benthic (at 32 m) | Oxic | True | 70 | - | 146 | EN ISO 22766:2020 |
| Marine | 29 | Bangka | Organic cotton t-shirt | Field: Disintegration | SEA2 | Eulittoral (Buried at 50 cm) | Fluctuating | True | 53 | - | 146 | EN ISO 22766:2020 |
| Marine | 29 | Bangka | Lyocell | Field: Disintegration | SEA2 | Eulittoral (Buried at 50 cm) | Fluctuating | True | 0 | - | 103 | EN ISO 22766:2020 |
| Marine | 29 | Bangka | Lyocell | Field: Disintegration | SEA2 | Eulittoral (Buried at 50 cm) | Fluctuating | True | 100 | - | 207 | EN ISO 22766:2020 |
| Marine | 29 | Bangka | Lyocell | Field: Disintegration | SEA3 | Benthic (at 32 m) | Oxic | True | 40 | - | 102 | EN ISO 22766:2020 |
| Marine | 29 | Bangka | Lyocell | Field: Disintegration | SEA3 | Benthic (at 32 m) | Oxic | True | 95 | - | 205 | EN ISO 22766:2020 |
| Marine | 29 | Bangka | Lyocell | Field: Disintegration | SEA3 | Benthic (at 32 m) | Oxic | True | 100 | - | 1312 | EN ISO 22766:2020 |
| Marine | 29 | Bangka | Denim blue jeans | Field: Disintegration | SEA2 | Eulittoral (Buried at 50 cm) | Fluctuating | True | 2 | - | 149 | EN ISO 22766:2020 |
| Marine | 29 | Bangka | Denim blue jeans | Field: Disintegration | SEA2 | Eulittoral (Buried at 50 cm) | Fluctuating | True | 100 | - | 1467 | EN ISO 22766:2020 |
| Marine | 29 | Bangka | Denim<br>blue jeans | Field:<br>Disintegration | SEA3 | Benthic<br>(at 32 m) | Oxic | True | 10 | - | 146 | EN ISO 22766:2020 |
| Marine | 29 | Bangka | Wood,<br>oak<br>veneer | Field:<br>Disintegration | SEA3 | Benthic<br>(at 32 m) | Oxic | True | 0 | - | 133 | EN ISO 22766:2020 |
| Marine | 29 | Bangka | Wood,<br>oak<br>veneer | Field:<br>Disintegration | SEA3 | Benthic<br>(at 32 m) | Oxic | True | 2 | - | 279 | EN ISO 22766:2020 |
| Marine | 29 | Bangka | Wood,<br>oak<br>veneer | Field:<br>Disintegration | SEA3 | Benthic<br>(at 32 m) | Oxic | True | 4 | - | 400 | EN ISO 22766:2020 |
| Marine | 29 | Bangka | Wood,<br>oak<br>veneer | Field:<br>Disintegration | SEA3 | Benthic<br>(at 32 m) | Oxic | True | 6 | - | 487 | EN ISO 22766:2020 |
| Marine | 29 | Bangka | Wood,<br>oak<br>veneer | Field:<br>Disintegration | SEA3 | Benthic<br>(at 32 m) | Oxic | True | 98 | - | 1592 | EN ISO 22766:2020 |
| Marine | 29 | Bangka | Banana<br>leaf | Field:<br>Disintegration | SEA3 | Benthic<br>(at 32 m) | Oxic | True | 70 | - | 133 | EN ISO 22766:2020 |

**Table SI6:**
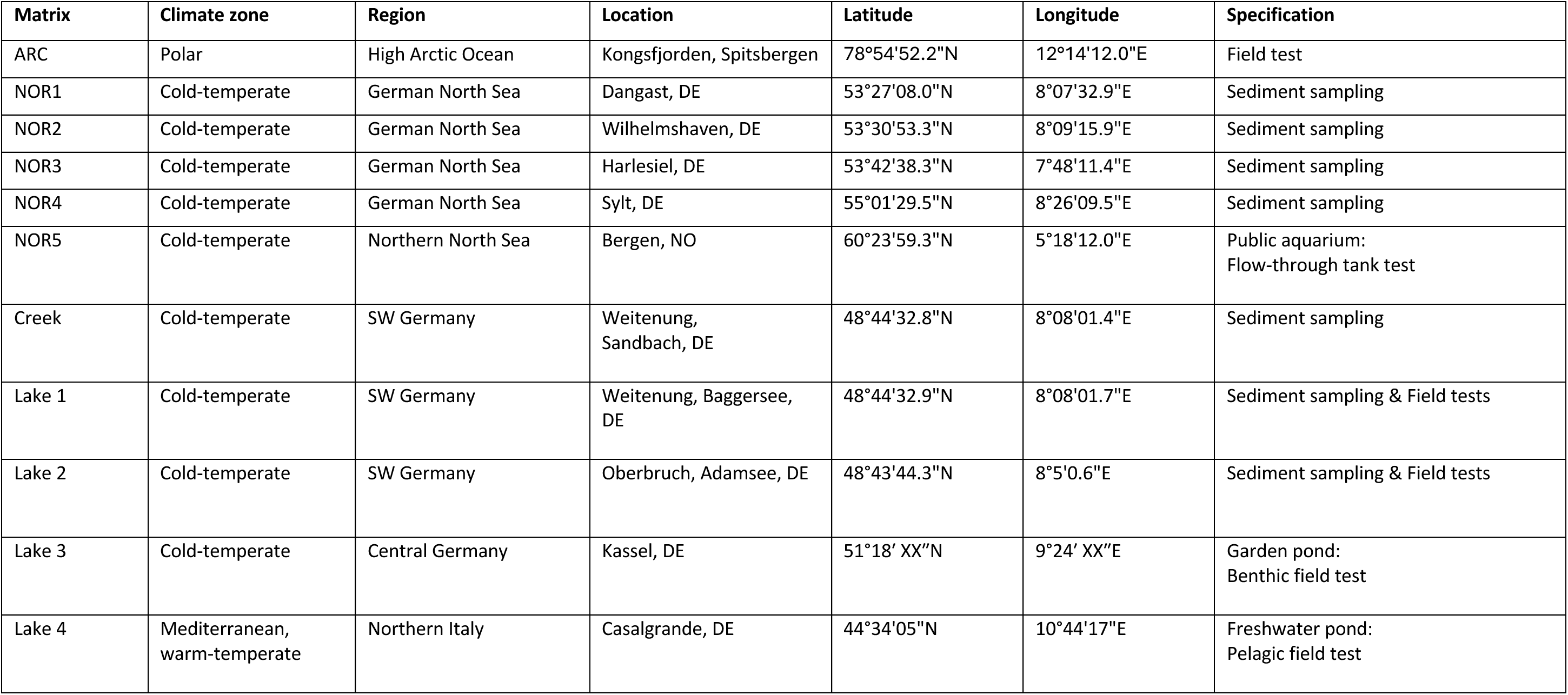
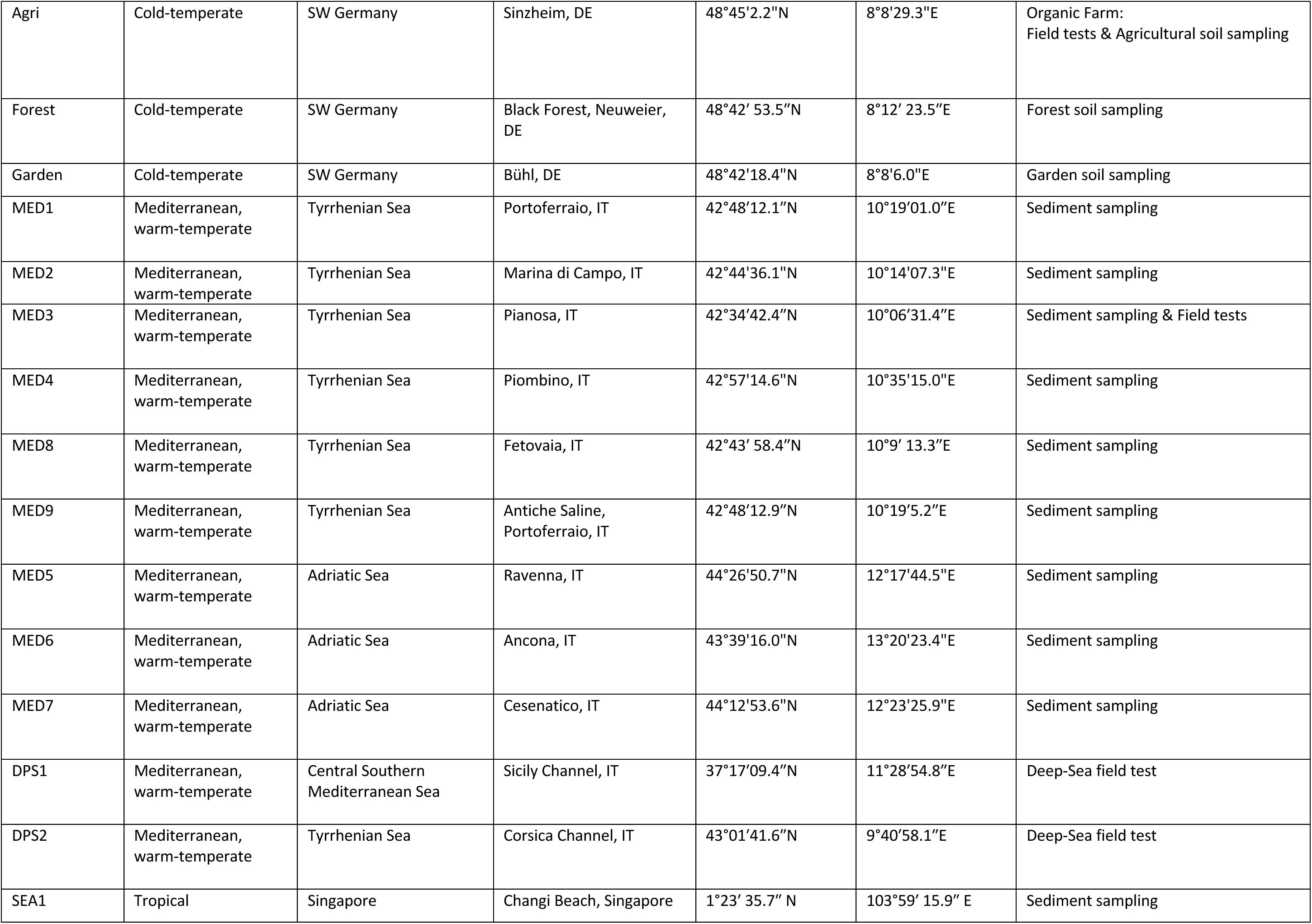
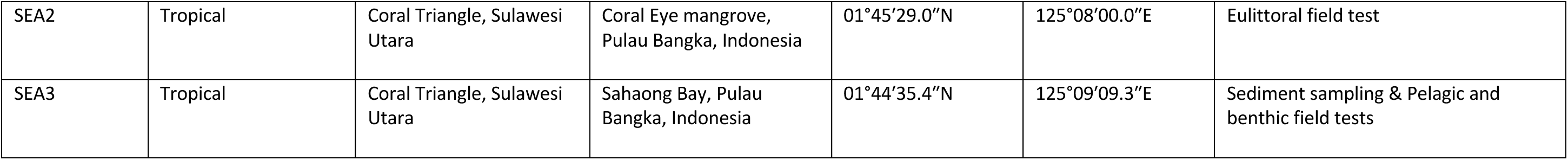
Locations of field tests and sediment sampling. Overview of sites used for field testing and sediment collection, including information on matrix, climate zone, region, specific location, and corresponding geographical coordinates (latitude and longitude). Additional site specifications are provided where relevant.

**Table SI7:** Temperatures of lab, tank and field tests. Overview of temperatures of the respective experiments, including the overall mean of daily temperatures, the mean minimum temperature, the mean maximum temperature and the absolute minimum and maximum temperatures.

|  | Habitat | Mean | Mean Min | Mean Max | Min | Max |
| --- | --- | --- | --- | --- | --- | --- |
| **Lab** | **All** | NA | 21±1 | 30±2 | 20 | 32 |
| **Lab** | **Compost** | 28±2 | NA | NA | 26 | 30 |
| **Lab** | **Soil** | 25±2 | NA | NA | 23 | 27 |
| **Lab** | **Freshwater** | NA | 21±1 | 28±1 | 20 | 29 |
| Lab | Freshwater 1 | 21±1 | NA | NA | 20 | 22 |
| Lab | Freshwater 2 | 28±1 | NA | NA | 27 | 29 |
| **Lab** | **Marine** | NA | 25±2 | 30±2 | 23 | 32 |
| Lab | Marine | 25±2 | NA | NA | 23 | 27 |
| Lab | Marine | 30±2 | NA | NA | 28 | 32 |
| **Field** | **All** | NA | 11 | 29 | -6,1 | 54,1 |
| **Field** | **Soil** | NA | 11 | 22 | -6,1 | 54,1 |
| Field | Greenhouse 1 | 16 | NA | NA | -6,1 | 47,7 |
| Field | Greenhouse 2 | 22 | NA | NA | 2,9 | 47,7 |
| Field | Greenhouse 3 | 15 | NA | NA | -2,2 | 31,6 |
| Field | Field1 | 14 | NA | NA | 2,6 | 28,3 |
| Field | Field2 | 20 | NA | NA | -2,1 | 54,1 |
| Field | Field3 | 14 | NA | NA | 2,6 | 28,3 |
| Field | Field4 | 13 | NA | NA | 5,1 | 31,6 |
| Field | Field5 | 11 | NA | NA | 2,6 | 21,9 |
| Field | Field6 | 14 | NA | NA | 2,6 | 28,3 |
| **Field** | **Freshwater** | 13,5 | 12 | 15 | 2 | 32,9 |
| Field | Lake 2 | 15 | NA | NA | 4,8 | 25,8 |
| Field | Lake 1 | 12 | NA | NA | 7 | 26,3 |
| Field | Lake 3 | 12 | NA | NA | 2 | 21,6 |
| Field | Lake 4 | 15 | NA | NA | 3,9 | 32,9 |
| **Field** | **Marine** | 25,8 | 14 | 29 | -1 | 38,4 |
| Field | SEA2 | 29 | NA | NA | 26 | 23,2 |
| Field | SEA3 | 29 | NA | NA | 23,2 | 31,5 |
| Field | MED8 | 26 | NA | NA | -1 | 38,4 |
| Field | MED9 | 25 | NA | NA | 14 | 27 |
| Field | MED3 | 20 | NA | NA | 14 | 23 |
| Field | MED3 | 19,5 | NA | NA | 14 | 25 |
| Field | MED9 | 22 | NA | NA | 17 | 27 |
| Field | MED8 | 19,5 | NA | NA | 4 | 35 |
| Field | DeepSea | 14 | NA | NA | NA | NA |
| **Mesocosms** | **All** | NA | 9 | 28 | 9 | 30,8 |
| **Mesocosms** | **Soil** | 23,5 | 23 | 24 | 13,6 | 30,8 |
| Mesocosms | Soil 1 | 24 | NA | NA | 13,6 | 30,8 |
| Mesocosms | Soil 4 | 23 | NA | NA | 13,9 | 26 |
| **Mesocosms** | **Freshwater** | NA | 18 | 28 | 18 | 28 |
| Mesocosms | Lake 2 | NA | 18 | 26 | 18 | 26 |
| Mesocosms | Creek | NA | 18 | 26 | 18 | 26 |
| Mesocosms | Lake 1 | NA | 26 | 28 | 26 | 28 |
| **Mesocosms** | **Marine** | 20 | 9 | 28 | 9 | 28 |
| Mesocosms | NOR5 | 9 | NA | NA | 9 | 9 |
| Mesocosms | MED3 | 14 | NA | NA | 14 | 14 |
| Mesocosms | MED1,2,4,5,6,7, NOR1,3,4 | 23 | NA | NA | 23 | 23 |
| Mesocosms | NOR4 | 26 | NA | NA | 26 | 26 |
| Mesocosms | SEA1, MED2,3 | 28 | NA | NA | 28 | 28 |

| Tank & Field |  |  |  |  |  |  |
| --- | --- | --- | --- | --- | --- | --- |
| Soil | All | NA | 11 | 22 | -6,1 | 54,1 |
| Freshwater | All | NA | 12 | 28 | 2 | 32,9 |
| Marine | All | NA | 9 | 29 | -1 | 38,4 |

### Part 2: PHOTO PANELS

**Figure SI11:**
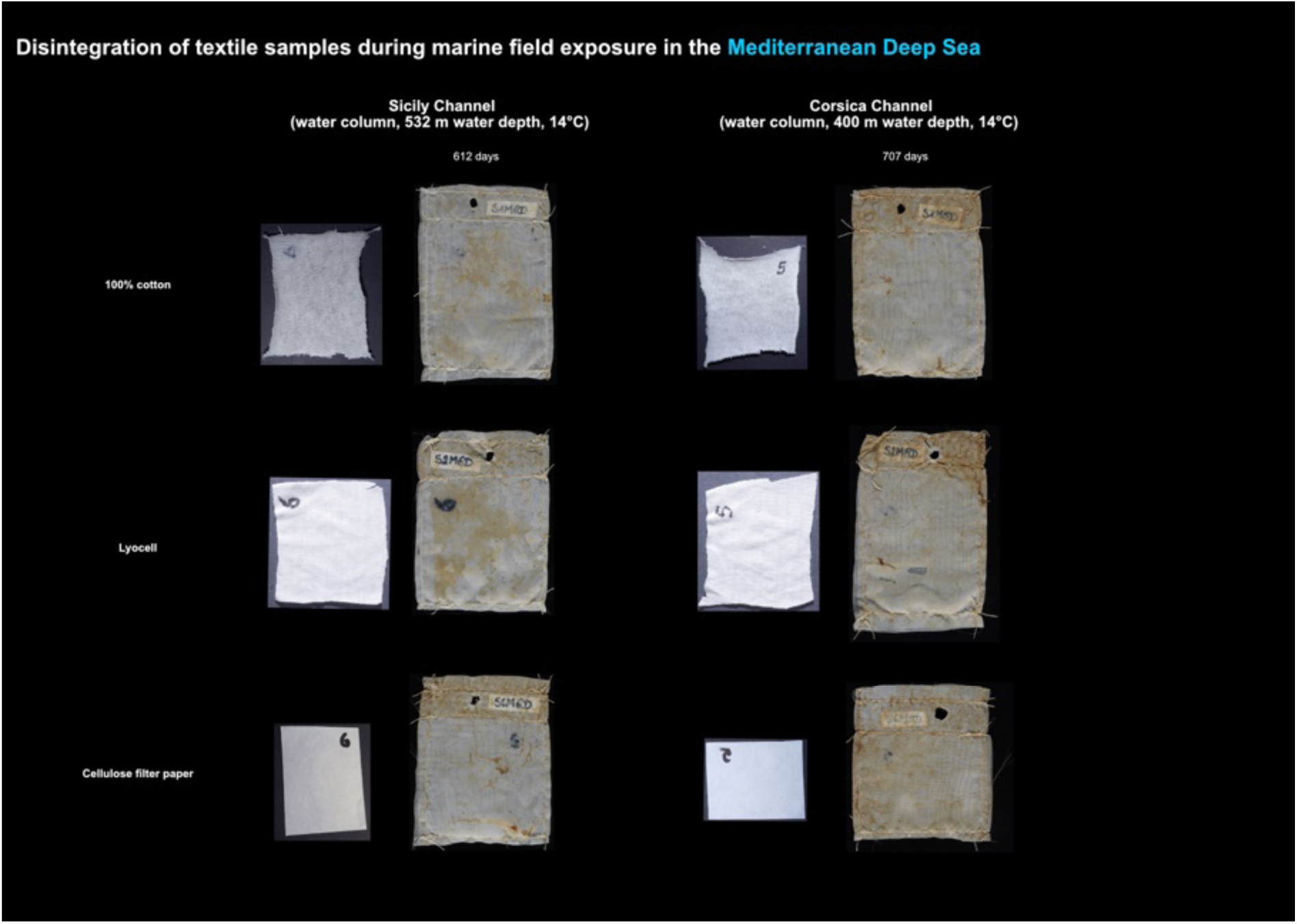
Woven fabric of cotton and lyocell compared to cellulose filter paper in a deep-sea water field test at 14°C. Note: The Mediterranean Sea has a special water circulation regime leading to minimum temperatures of ∼13-14°C even at greatest depth.

**Figure 12:**
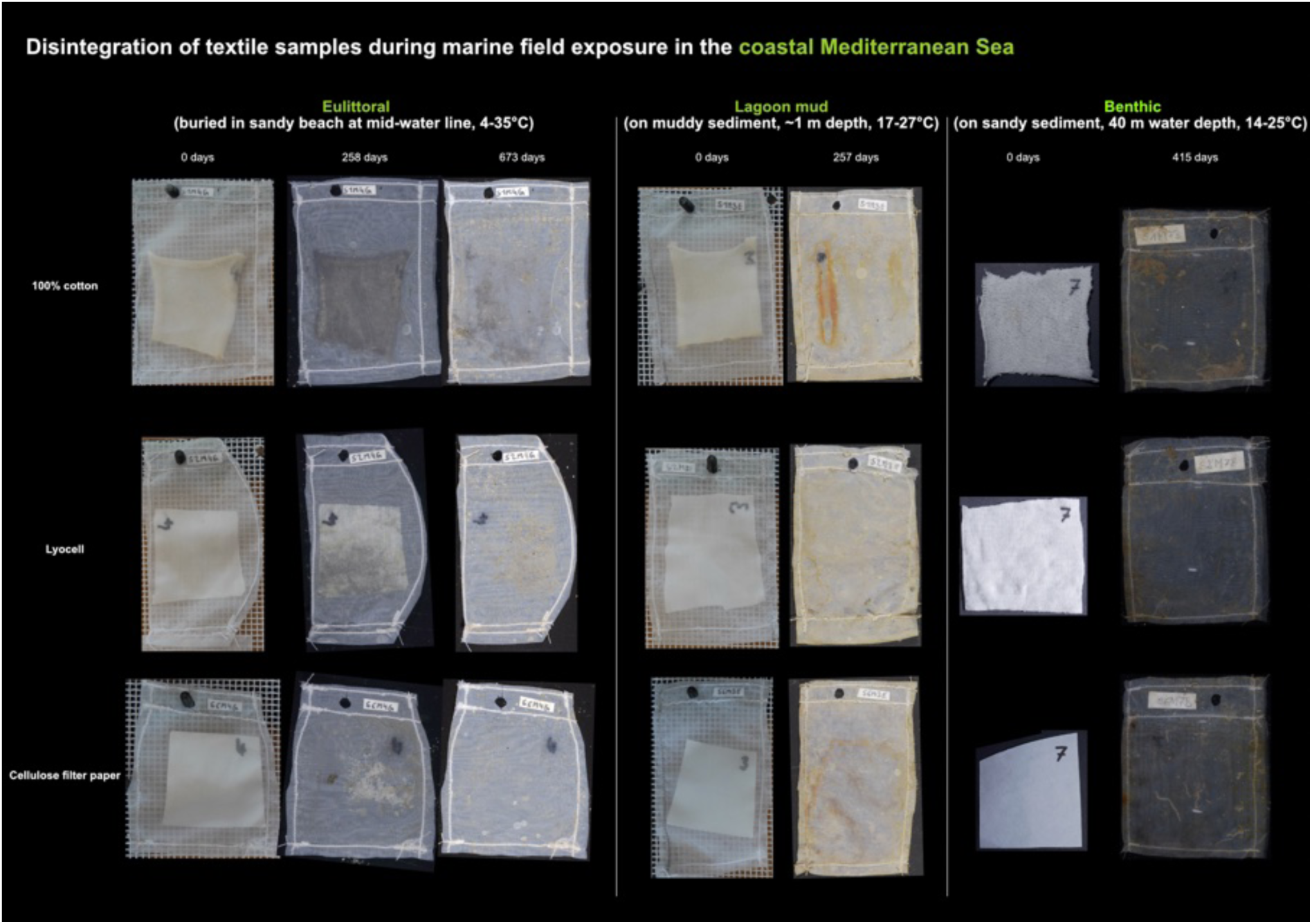
Woven fabric of cotton and lyocell compared to cellulose filter paper in beach sand (eulittoral, left), shallow-water lagoon mud (central), and on the seafloor (benthic, right) at 40 m depth in the Mediterranean Sea at ambient temperatures.

**Figure SI13:**
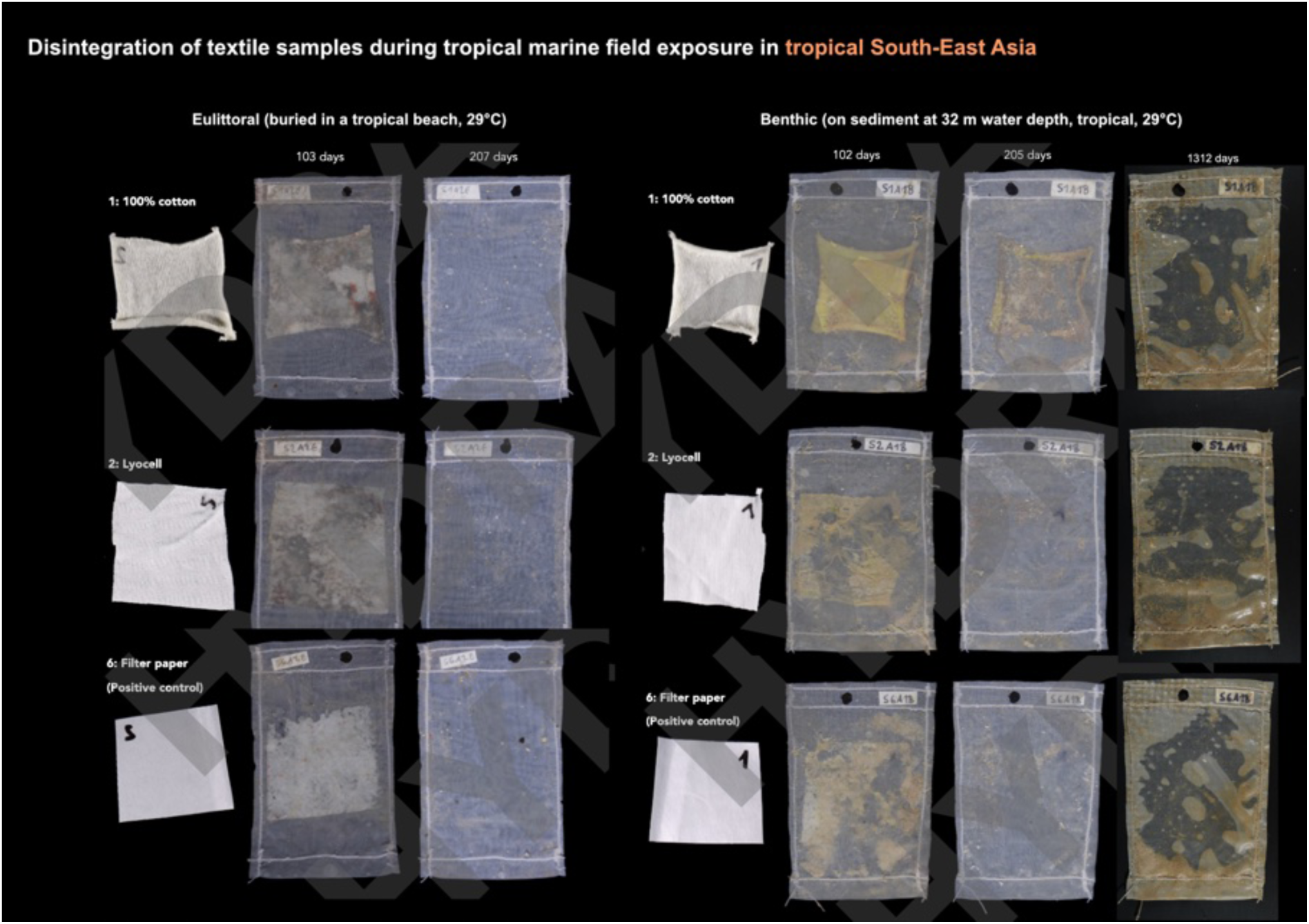
Disintegration of woven fabric of cotton and lyocell compared to cellulose filter paper in tropical beach (left) and seafloor (right) field tests at 29°C. Complete disintegration in the beach experiment after 207 d. The sediment in the beach experiments showed higher activity towards cotton than in the seafloor experiments.

**Figure SI14:**
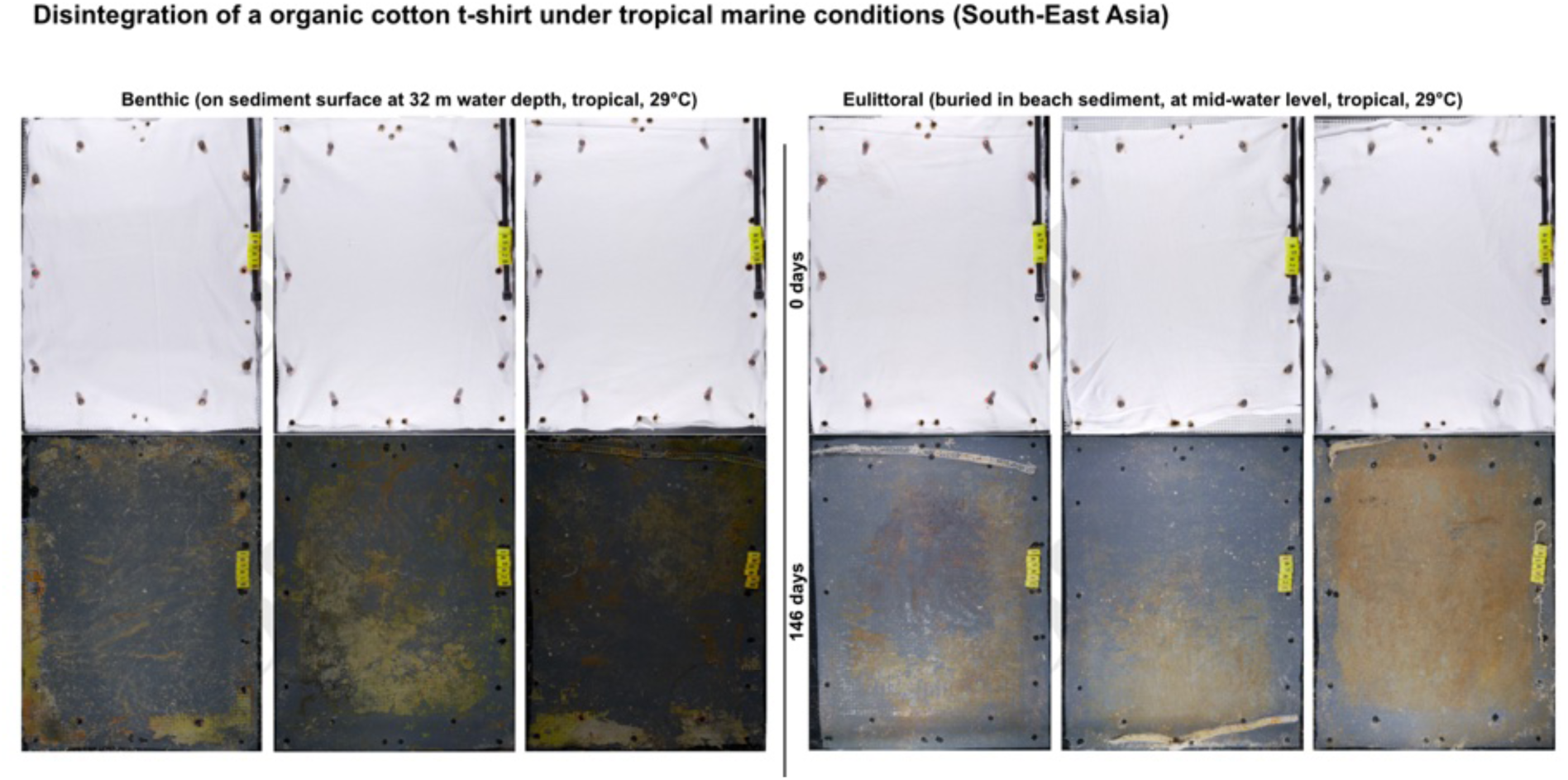
Woven fabric cut from an organic cotton t-shirt on the seafloor (benthic) at 32 m depth and in beach sand (eulittoral), at 29°C.

**Figure SI15:**
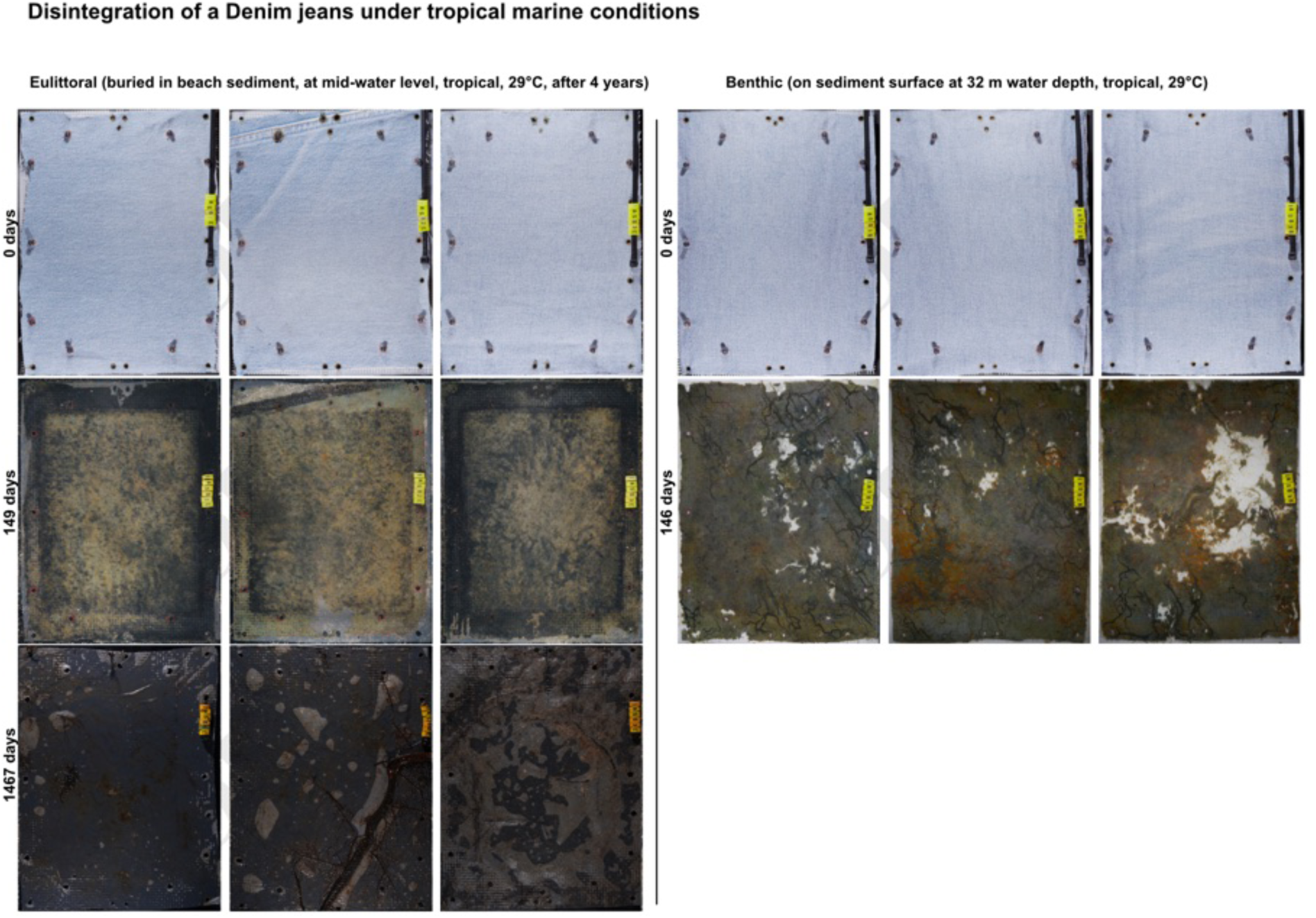
Woven fabric cut from Denim blue jeans exposed in beach sand (eulittoral, left) and on the seafloor (benthic, right) at 32 m depth, at 29°C.

**Figure SI16:**
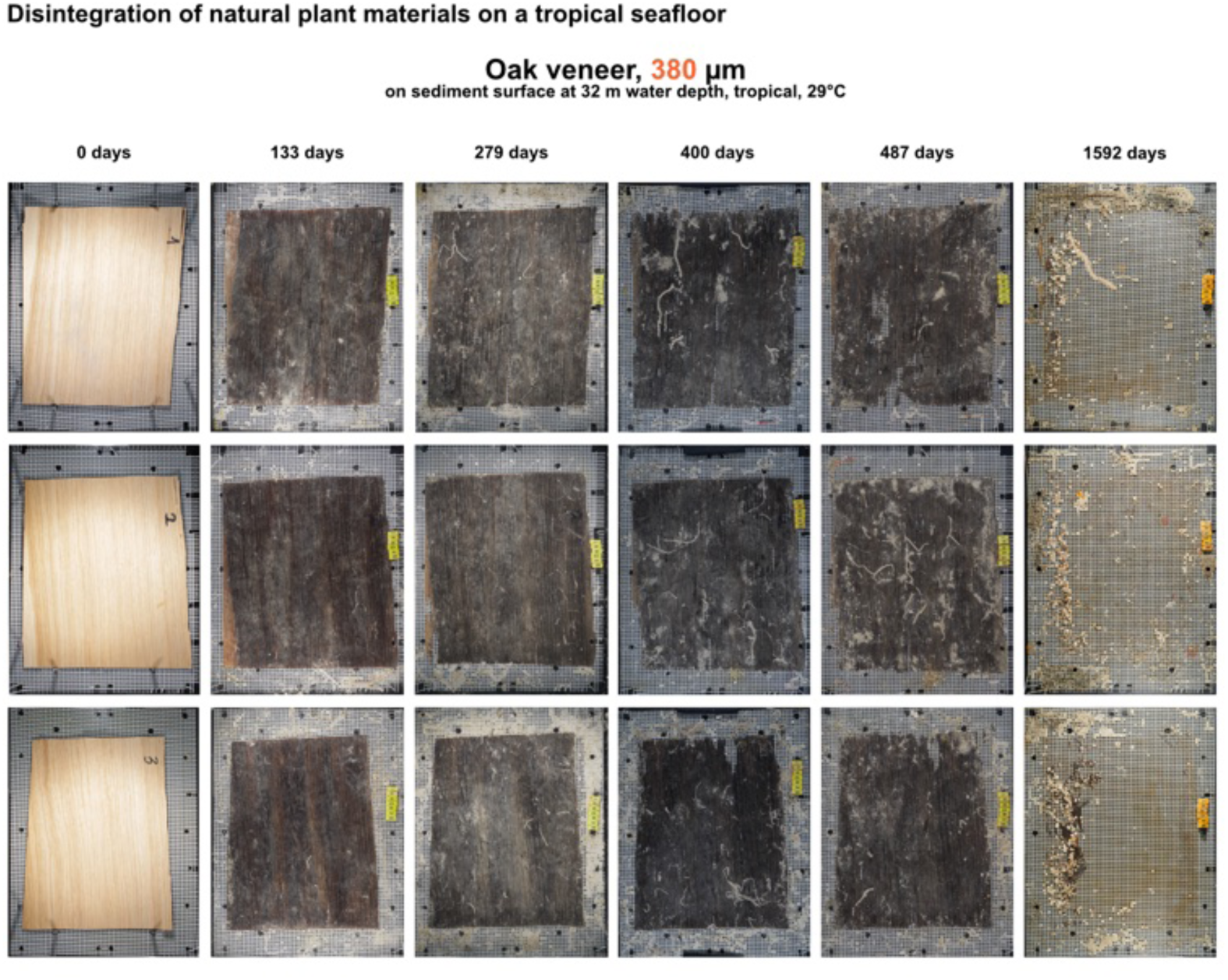
Oak veneer exposed at the seafloor at 32 m depth, at 29°C.

**Figure SI17:**
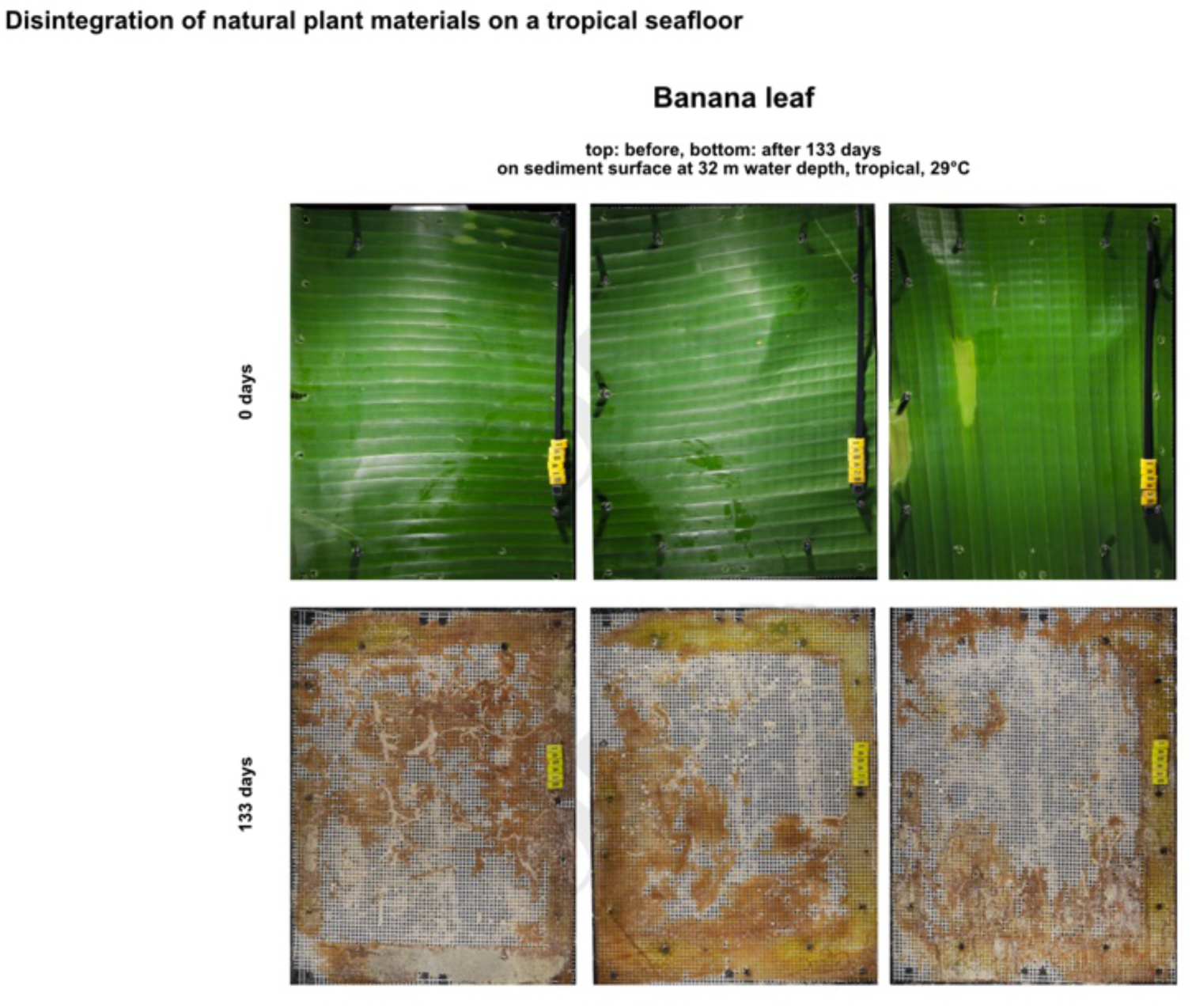
Fresh banana leaves exposed at the seafloor at 32 m depth, at 29°C.

**Figure SI18:**
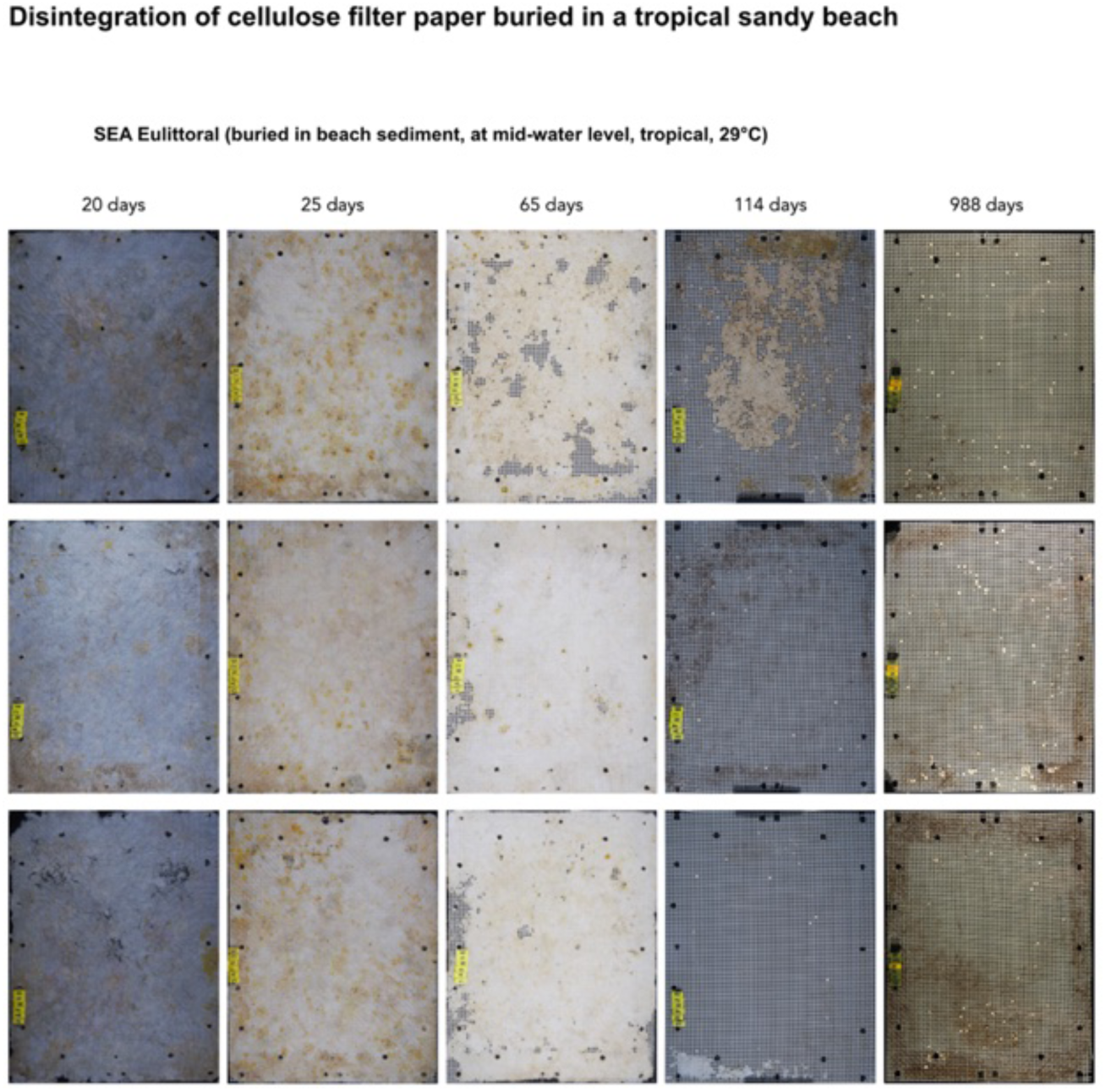
Cellulose Filter Paper exposed in tropical beach sand (eulittoral) in Southeast Asia, at 29°C, corresponding to treatment T55 (for details see tab. 6). Disintegration was just well noticeable but not completed yet after ∼2 months in the eulittoral. After ∼4 months, in two replicates, no trace of the material was left. After ∼3 years all material was gone. The half-live was modeled to t_0,5_ = 90 d (CI: 78.93-102.83).

**Figure SI19:**
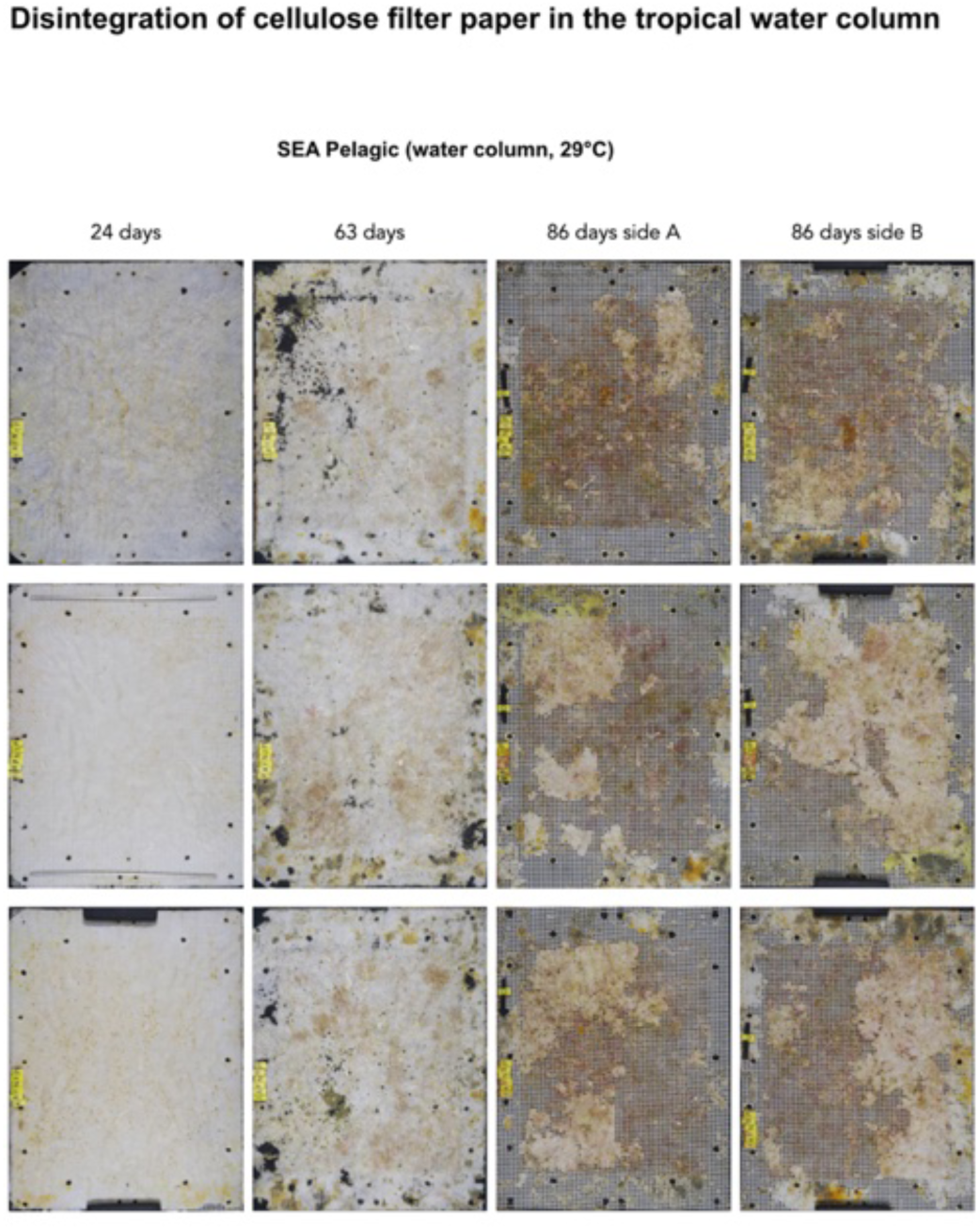
Cellulose Filter Paper exposed in the water column in Southeast Asia at 20 m depth, at 29°C, corresponding to treatment T56 (for details see tab. 6). Disintegration was just well noticeable but not completed yet after ∼3 months in the pelagic. The half-live was modeled to t_0,5_ = 97 d (CI: 90.91-115.35).

**Figure SI20:**
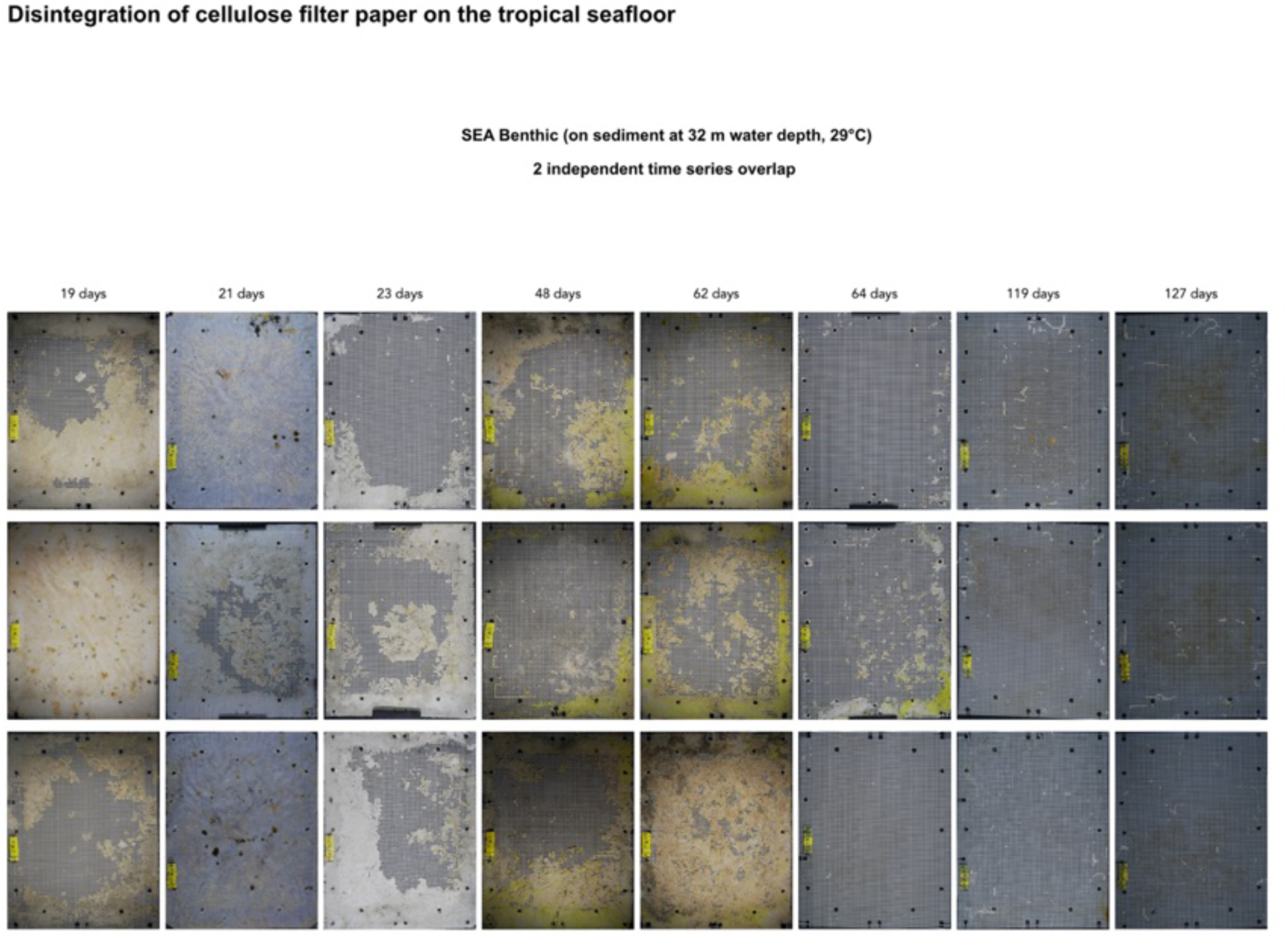
Cellulose Filter Paper exposed at the seafloor at 32 m depth in Southeast Asia, at 29°C, corresponding to treatment T54 (for details see tab. 6). Disintegration was heterogenous between replicates but already well noticeable after 3 weeks and completed after 4 months in the benthic. The photo panel (fig. SI20) represents the samples of 2 repeated experiments and shows the heterogeneity between experiments and replicate samples within one sampling interval. The half-live was modeled to t_0,5_ = 34 d (CI: 22.04-52.88).

